# Inter-species conservation of organisation and function between non-homologous regional centromeres

**DOI:** 10.1101/309815

**Authors:** Pin Tong, Alison L. Pidoux, Nicholas R.T. Toda, Ryan Ard, Harald Berger, Manu Shukla, Jesus Torres-Garcia, Carolin A. Mueller, Conrad A. Nieduszynski, Robin C. Allshire

**Author notes:** These authors made an equal contribution (Pin Tong, Alison L. Pidoux). Present addresses: UPMC CNRS, Roscoff Marine Station, Place Georges Teissier, 29680 Roscoff, France. Copenhagen Plant Science Centre, University of Copenhagen, Bülowsvej 34, 1870 Frederiksberg C, Denmark (Ryan Ard). Symbiocyte, Universität für Bodenkultur Wien, University of Natural Resources and Life Sciences, Vienna, Austria (Harald Berger).

## Abstract

Despite the conserved essential function of centromeres, centromeric DNA itself is not conserved^1–4^. The histone-H3 variant, CENP-A, is the epigenetic mark that specifies centromere identity^5–8^. Paradoxically, CENP-A normally assembles on particular sequences at specific genomic locations. To gain insight into the specification of complex centromeres we took an evolutionary approach, fully assembling genomes and centromeres of related fission yeasts. Centromere domain organization, but not sequence, is conserved between *Schizosaccharomyces pombe, S. octosporus* and *S. cryophilus* with a central CENP-A^Cnp1^ domain flanked by heterochromatic outer-repeat regions. Conserved syntenic clusters of tRNA genes and 5S rRNA genes occur across the centromeres of *S. octosporus* and *S. cryophilus*, suggesting conserved function. Remarkably, non-homologous centromere central-core sequences from *S. octosporus* are recognized in *S. pombe*, resulting in cross-species establishment of CENP-A^Cnp1^ chromatin and functional kinetochores. Therefore, despite the lack of sequence conservation, *Schizosaccharomyces* centromere DNA possesses intrinsic conserved properties that promote assembly of CENP-A chromatin. Thus, centromere DNA can be recognized and function over unprecedented evolutionary timescales.

---

Centromeres are the chromosomal regions upon which kinetochores assemble to mediate accurate chromosome segregation. Evidence suggests that both genetic and epigenetic influences define centromere identity^1,2,4,7,9^. *S. pombe*, a paradigm for dissecting complex regional centromere function, has demarcated centromeres (35-110 kb) with a central domain assembled in CENP-A^Cnp1^ chromatin, flanked by outer-repeat elements assembled in RNAi-dependent heterochromatin, in which histone-H3 is methylated on lysine-9 (H3K9)^10–13^. Heterochromatin is required for establishment but not maintenance of CENP-A^Cnp1^ chromatin^6,14^. We have proposed that it is not the sequence *per se* of *S. pombe* central-core that is key in its ability to establish CENP-A chromatin, but the properties programmed by it^15^. To investigate whether these properties are conserved we have determined the centromere sequences of other *Schizosaccharomyces* species and tested their cross-species functionality.

Long-read (PacBio) sequencing permitted complete assembly of the genomes across centromeres of *S. octosporus* (11.9 Mb) and *S. cryophilus* (12.0 Mb), extending genome sequences^16^ to telomeric or subtelomeric repeats or rDNA arrays (**Supplementary Figs. 1–3, Supplementary Tables 1,2**). Consistent with their closer evolutionary relationship^16,17^, *S. octosporus* and *S. cryophilus* (32 My separation, compared to 119 My separation from *S. pombe*) exhibit greatest synteny (**Fig. 1a**). Synteny is preserved adjacent to centromeres (**Fig. 1b**). Circos plots indicate a chromosome arm translocation occurred within two ancestral centromeres to generate *S. cryophilus cen2* (*S.cry-cen2*) and *S.cry-cen3* relative to *S. octosporus* and *S. pombe* (**Fig. 1b**). Despite centromere-adjacent synteny, *Schizosaccharomyces* centromeres lack detectable sequence homology (see below). All centromeres contain a central domain: central-core (*cnt*) surrounded by inverted repeat (*imr*) elements unique to each centromere (**Fig. 2, Supplementary Fig. 4, Supplementary Tables 3–6**). CENP-A^Cnp1^ localises to fission yeast centromeres (**Fig. 2a**) and ChIP-Seq indicates that central domains are assembled in CENP-A^Cnp1^ chromatin, flanked by various outer-repeat elements assembled in H3K9me2-heterochromatin (**Fig. 2b,c**). Despite the lack of sequence conservation, *S. octosporus* and *S. cryophilus* centromere organisation is strongly conserved with that of *S. pombe*, having CENP-A^Cnp1^-assembled central domains separated by clusters of tRNA genes from outer-repeats assembled in heterochromatin^10–11^ (**Supplementary Fig. 5, Supplementary Tables 7,8**). In contrast, our analyses of partially-assembled, transposon-rich centromeres of *S. japonicus* reveals the presence of heterochromatin on all classes of transposons and CENP-A on only two classes (**Supplementary Fig. 5, Supplementary Table 9**)^16^.

**Figure 1:**
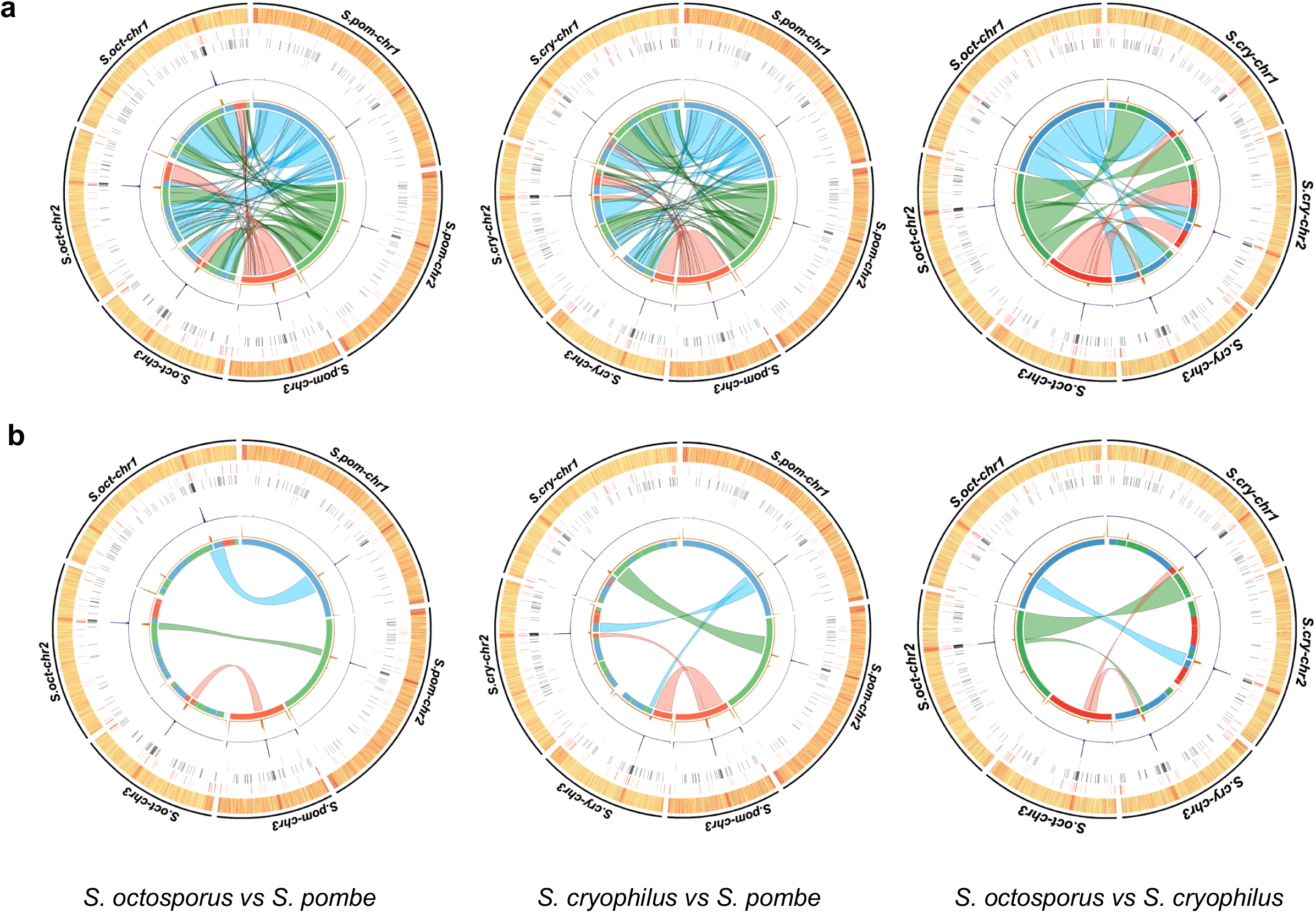
Genome organisation and synteny in *Schizosaccharomyces*. a) Circos plots depicting pairwise *S. pombe, S. octosporus* and *S. cryophilus* genome synteny. Rings from outside to inside represent: chromosomes; GC content (high: red, low: yellow); 5S rDNAs (red); tDNAs (black); LTRs (green); CENP-A^Cnp1^ ChIP-seq (purple); H3K9me2 ChIP-seq (orange); innermost ring and coloured connectors indicate regions of synteny between species. *S. pombe* chromosomes are indicated by blue (*S.pom-chr1*), green (*S.pom-chr2*), red (*S.pom-chr3*) in the left and right panels and regions of synteny on *S. octosporus* and *S. cryophilus* chromosomes, respectively, are indicated in corresponding colours. A similar designation is used for *S. octosporus* chromosomes in the middle panel. b) Circos plot isolating regions adjacent to centromeres highlighting preserved synteny and an intra-centromeric chromosome arm swap involving *S. cryophilus cen2* and *cen3* relative to *S. pombe* and *S. octosporus*.

**Figure 2:**
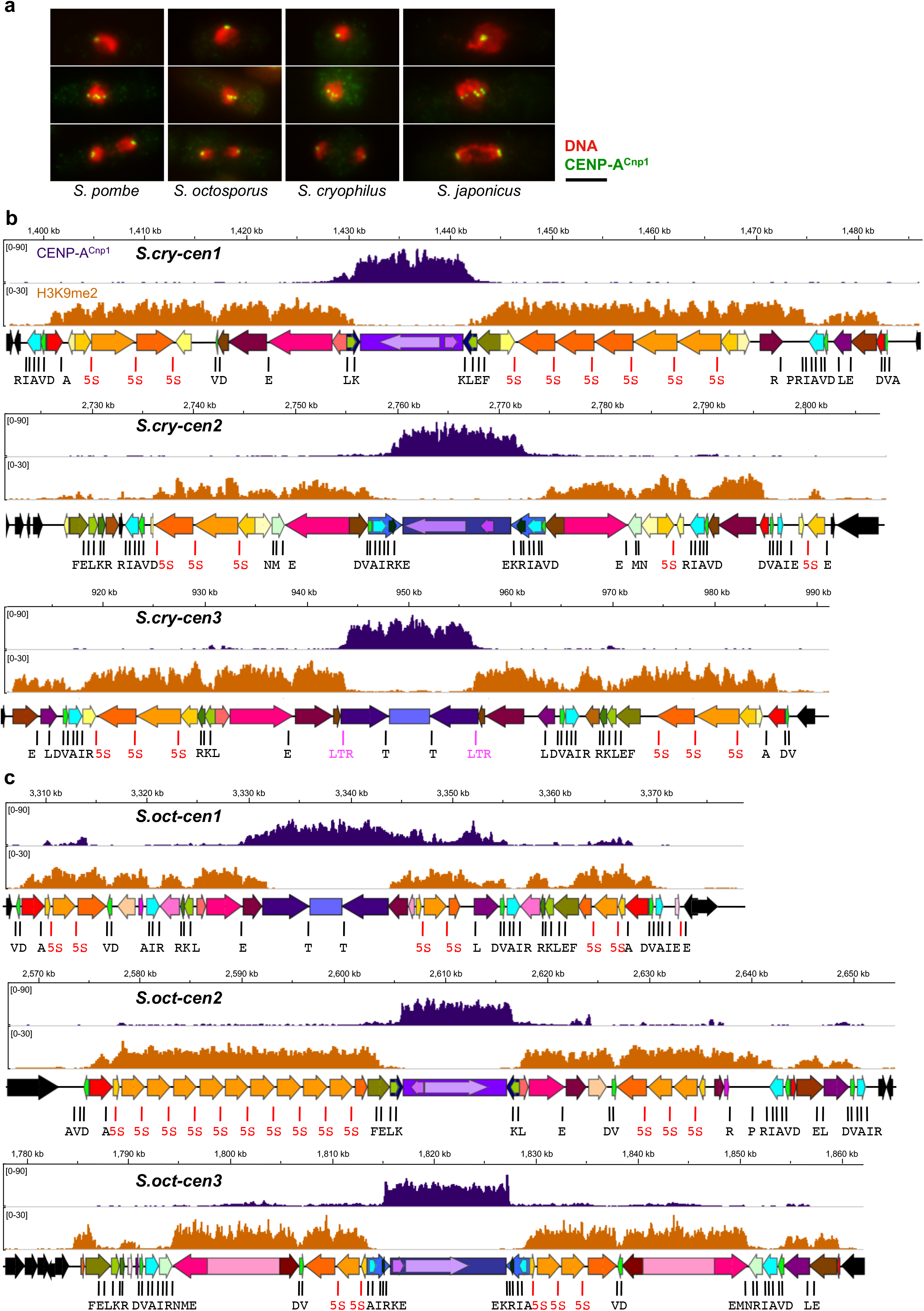
Domain organisation of *Schizosaccharomyces* centromeres. a) Immunostaining of centromeres in indicated *Schizosaccharomyces* species with anti-CENP-A^Cnp1^ antibody (green) and DNA staining (DAPI; red). Scale bar, 5 μm. (b) *S. cryophilus* centromere organisation indicating DNA repeat elements. ChIP-seq profiles for CENP-A^Cnp1^ (purple) and H3K9me2-heterochromatin (orange) are shown above each centromere. Positions of tDNAs (single-letter code of cognate amino acid; black), 5S-rDNAs (red), and solo LTRs are indicated (pink). Central cores (cnt – purples) inner-most repeats (imr – blue shades). 5S-associated repeats (cFSARs – orange shades); tDNA-associated repeats (TARs) containing clusters of tDNAs (green shades); heterochromatic repeats (cHR) and TARs associated with single tDNAs (various colours: brown/pink/red). cTAR-14s, containing retrotransposon remnants (deep pink). For details, including individual repeat annotation, see **Supplementary Fig. 4** and **Supplementary Tables 3,4**. (c) *S. octosporus* centromere organisation indicating DNA repeat elements. Labelling and shading as in (b). Only oTAR-14ex (pale pink part) contain retrotransposon remnants. Colouring is indicative of homology within each species but only of possible repeat equivalence (not homology) between species; see **Supplementary Table 5,6,19**.

Numerous 5S rRNA genes are located in the heterochromatic outer-repeats of *S. octosporus* and *S. cryophilus* centromeres (but not *S. pombe*) (**Fig. 1a, Supplementary Tables 10,11**). Almost all (25/26; 20/20) are within Five-S-Associated Repeats (FSARs; 0.6-4.2 kb) (**Fig. 3a**), encompassing ~35% of outer-repeat regions. FSARs exhibit 90% intra-class homology (**Supplementary Table 12**), but no interspecies homology. The three types of FSAR repeats almost always occur together, in the same order and orientation, but vary in copy number: *S. octosporus*: (oFSAR-1)_1_(oFSAR-2)_1-9_(oFSAR-3)_1_; *S. cryophilus*: (cFSAR-1)_1-3_(cFSAR-2)_1-2_(cFSAR-3)_1_. Both sides of *S. octosporus* and *S. cryophilus* centromeres contain at least one FSAR-1-2-3 array, except the right side of *S.cry-cen2* with two lone cFSAR-3 elements (**Fig. 3a, Supplementary Fig. 4**). *S. cryophilus* cFSAR-2 and cFSAR-3 repeats share ~400 bp homology (88% identity), constituting *hsp16* heat-shock protein ORFs (**Fig. 3a,b, Supplementary Table 13**) that are intact, implying functionality, selection and expression in some situations. Phylogenetic gene trees indicate that cFSAR-3-hsp16 genes are more closely related with each other than with those in subtelomeric regions or cFSAR-2s (**Fig. 3b**), consistent with repeat homogenisation^18–20^. cFSAR-1s contain an eroded ORF with homology to a small hypothetical protein and *S. octosporus* oFSAR-2s contain a region of homology with a family of membrane proteins (**Fig. 3a**). The functions of centromere-associated *hsp16* genes and other ORF-homologous regions remain to be explored.

**Figure 3:**
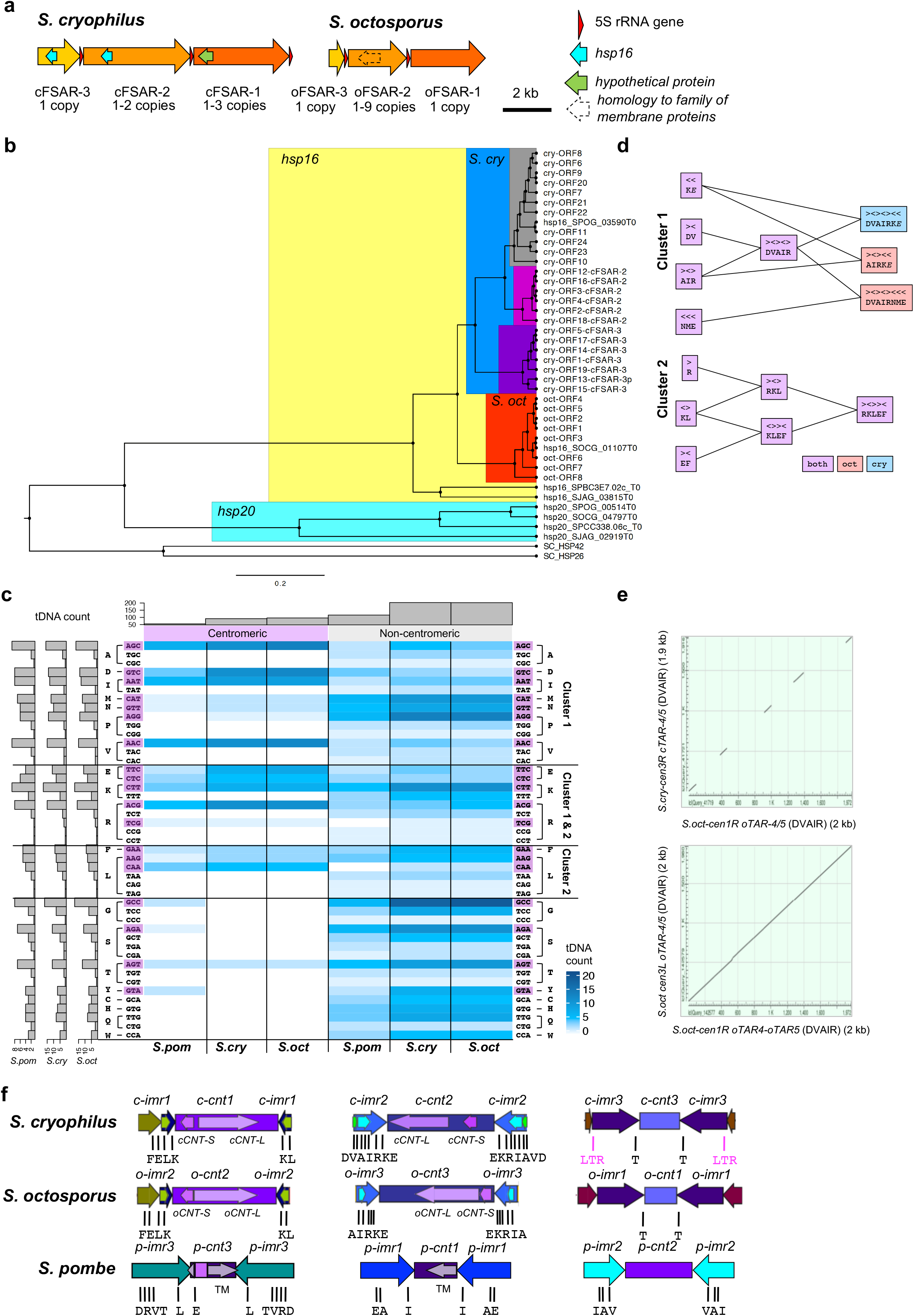
*S. cryophilus* and *S. octosporus* contain conserved clusters of tDNAs and similar non-homologous repeat elements. (a) Schematic of *S. cryophilus* and *S. octosporus* FSAR repeats, indicating positions of 5S-rDNAs, *hsp16* genes and other ORFs. Copy number of each FSAR within centromeric arrays is indicated. (b) Phylogenetic relationship of *S. cryophilus* centromeric *hsp16* genes with genomic *hsp16* and *hsp20* genes of *S. cryophilus, S. octosporus, S. pombe* and *S. japonicus* (c) Heat map of tDNA frequency at centromeric and non-centromeric sites (blue shades) for *S. pombe, S. cryophilus*, and *S. octosporus*. Anticodons and cognate amino acids indicated right (purple: present at centromeres). Clusters containing these tDNAs indicated. Histogram *(top):* total tDNA frequencies in centromeres and non-centromeric sites of indicated species. Histogram *(left):* tDNA frequencies in each species. (d) Depiction of centromeric tDNA clusters and sub-clusters. Combinations of 2 or 3 tDNAs subclusters present in both species (purple) or specific to *S. octosporus* (red) *or S. cryophilus* (blue) of are indicated (single-letter code of cognate amino-acid; arrows indicate plus or minus strand). (e) *Top*: Dot-plot alignment (MEGABLAST) showing synteny between oTAR-4/oTAR-5 (DVAIR-Cluster 1) from *S.oct-cen1R* (chr1:3355194-3357165) with oTAR-4/oTAR-5 (DVAIR-Cluster 1) from *S.cry-cen3R* (chr3:964707-966623). *Bottom*: Dot-plot of oTAR-4/oTAR-5 (DVAIR-Cluster 1) from *S.oct-cen1R* (chr1:3355194-3357165) and oTAR-4/oTAR-5 (DVAIR-Cluster 1) from *S.oct-cen3L* (chr3:1791072-1793051). (f) Schematic of central domain similarity between species. Central cores (purple shades), *imr* (blues), TARs containing tDNA clusters (greens). Long (*CNT-L*) and short (*CNT-S*) central core repeats are indicated. tDNAs indicated in single-letter amino acid code. Colours highlight similarity of organisation between species and indicates homology within, not between, species.

*S. cryophilus* heterochromatic outer-repeats contain additional repetitive elements, including a 6.2 kb element (cTAR-14) with homology to the retrotransposon *Tcry1* and transposon remnants at the mating-type locus^16^ (**Figs. 1a,2b, Supplementary Fig. 4** and **Supplementary Tables 3,4,14**). *Tcry1* is located in the chrIII-R subtelomeric region (**Supplementary Figs. 3,4, Supplementary Table 1**). Although no retrotransposons have been identified in *S. octosporus*, remnants are present in the mating-type locus and oTAR-14ex in *S.oct-cen3* outer-repeats (**Fig. 2c, Supplementary Figs. 1,4** and **Supplementary Tables 5,6,15**). Hence, transposon remnants, FSARs and other repeats are assembled in heterochromatin at *S. octosporus* and *S. cryophilus* centromeres and potentially mediate heterochromatin nucleation.

tDNA clusters occur at transitions between CENP-A and heterochromatin domains in two of three centromeres in *S. octosporus* (*S.oct-cen2, S.oct-cen3*) and *S. cryophilus (S.cry-cen1, S.cry-cen2)*, and are associated with low levels of both H3K9me2 and CENP-A^Cnp1^ (**Fig. 2b,c**), suggesting that they may act as boundaries, as in *S. pombe*^21–23^. No tDNAs demarcate the CENP-A/heterochromatin transition at *S.cry-cen3*. Instead, this transition coincides precisely with 270-bp LTRs (**Fig. 2b, Supplementary Tables 3,4,14**), which may also act as boundaries^24–26^. Like tDNAs, LTRs are regions of low nucleosome occupancy, which may counter spreading of heterochromatin^26,27^. In addition, tDNA clusters occur near the extremities of all centromeres in both species, separating heterochromatin from adjacent euchromatin. tDNAs and LTRs are thus likely to act as chromatin boundaries at fission yeast centromeres.

A high proportion (~32%) of tRNA genes in *S. pombe, S. octosporus*, and *S. cryophilus* genomes are located within centromere regions^28^ (**Figs. 1a,3c; Supplementary Tables 16–18**). Centromeric tDNAs are intact and are conserved in sequence with their genome-wide counterparts, indicating that they are functional genes. Two major, conserved tDNA clusters reside exclusively within *S. octosporus* and *S. cryophilus* centromeres (p-value<0.00001; q-value<0.05) (**Fig. 3c,d**). Cluster1 comprises several subclusters of 2-3 tDNAs in various combinations of up to 8 tDNAs, whilst Cluster2 contains up to 5 tDNAs (**Fig. 3d**); 17 different tDNAs (14 amino-acids) are represented, none of which are unique to centromeres (**Fig. 3c**). Intriguingly, the order and orientation of tDNAs within clusters is conserved between species, but intervening sequence is not (**Fig. 3d,e**). Strikingly, as well as local tDNA cluster conservation, inspection of centromere maps reveals synteny of tDNAs and clusters across large portions of *S. octosporus* and *S. cryophilus* centromeres. For example, the tDNA order AIR-RKL-E-T-T-L-DVAIR-RKLEF-A-DV (single-letter code) is observed at *S.oct-cen1* and *S.cry-cen3* (**Supplementary Fig. 6**). This synteny, together with both possessing small central-cores and long *imrs* suggests that these two centromeres are ancestrally related (**Fig. 3f**). Similarly, at *S.oct-cen3* and *S.cry-cen2*, tDNAs occur in the order NME-DV-AIRKE-EKRIA-VD-EMN-RIAVD, and at *S.oct-cen2* and *S.cry-cen1* the same tDNAs are present in the *imr* repeats and beyond (FELK-KL-E-DV). Central-cores have similar sizes and structures in the two species, each containing long (oCNT-L(6.4 kb); cCNT-L(6.0 kb)) and short (oCNT-S(1.2 kb); cCNT-S(1.3 kb)) species-specific repeats (**Fig.3f, Supplementary Tables 3–6,19**). CNT-repeats are arranged head-to-tail at one centromere and head-to-head at the other centromere in each species. Together, these similarities suggest ancestral relationships between *S.oct-cen2* and *S.cry-cen1, So-cen3* and *Scry-cen2*. Further, in places where synteny appears to break down, patterns of tDNA clusters suggest specific centromeric rearrangements occurred between the species. For instance, tDNA clusters at the edges of *S.cry-cen2R* and *S.cry-cen3L* are consistent with an inter-centromere arm translocation relative to *S.oct-cen1R* and *S.oct-cen2R*, indicated by gene synteny maps (**Figs. 1b, 4a** and **Supplementary Fig. 6**).

No central-core sequence homology was revealed between species using BLASTN. To identify potential underlying centromere sequence features, k-mer frequencies (5-mers), normalized for centromeric AT-bias, were used in Principal Component Analysis. CENP-A-associated regions of all three genomes group together, distinct from the majority of non-centromere sequences (p-value, 9.3 × 10^−7^) (**Fig. 4b,c**). Interestingly, *S. pombe* neocentromere-forming regions^29^ also cluster separately from other genomic regions, sharing sequence features with centromeres.

**Figure 4:**
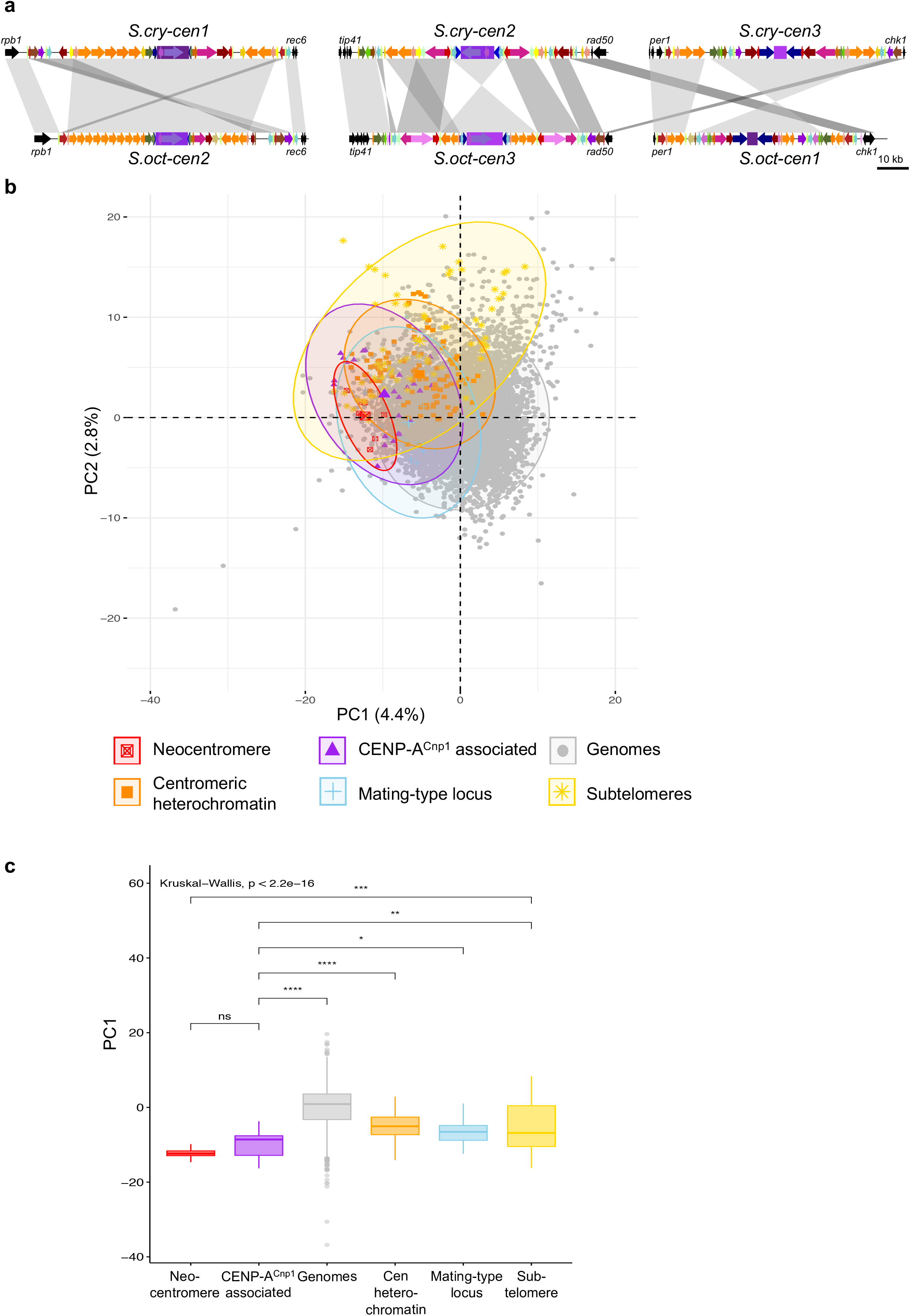
*Schizosaccharomyces* centromeres share ancestry and sequence features. (a) Structural alignment of putatively equivalent centromere repeat elements of *S. cryophilus* and *S. octosporus* to highlight potential centromere rearrangements during evolution (b) Principal Component Analysis PC1 and PC2 of 5-mer frequencies of three fission yeast genomes. Genome regions (12 kb window) were assigned to one of 5 specific annotated groups (CENP-A^Cnp1^-associated (purple), centromeric heterochromatin (orange), mating-type locus (blue), subtelomeres (yellow), neocentromere-forming regions^29^ (red), or other genome regions (grey). For each group the oval line encloses 95% of the data points. (c) Boxplot Principal Component PC1 of each group. Colours as in b. Mean comparison between groups was used (p-value: >0.05, ns; >0.01, *; >0.001, **; >0.0001, ***; <0.0001, ****)^60^. Centre line, medium; box limits, upper and lower quartiles; whiskers, 1.5 × interquartile range; points, outliers.

K-mer analysis and conserved centromeric organisation prompted us to investigate cross-species functionality of protein and DNA components of *Schizosaccharomyces* centromeres. GFP-tagged CENP-A^Cnp1^ protein from each species localised to *S. pombe* centromeres and complemented the *cnp1-1* mutant^30^ (**Fig. 5a-c**), indicating that heterologous CENP-A proteins assemble and function at *S. pombe* centromeres, despite normally assembling on non-homologous sequences in their respective organisms.

**Figure 5:**
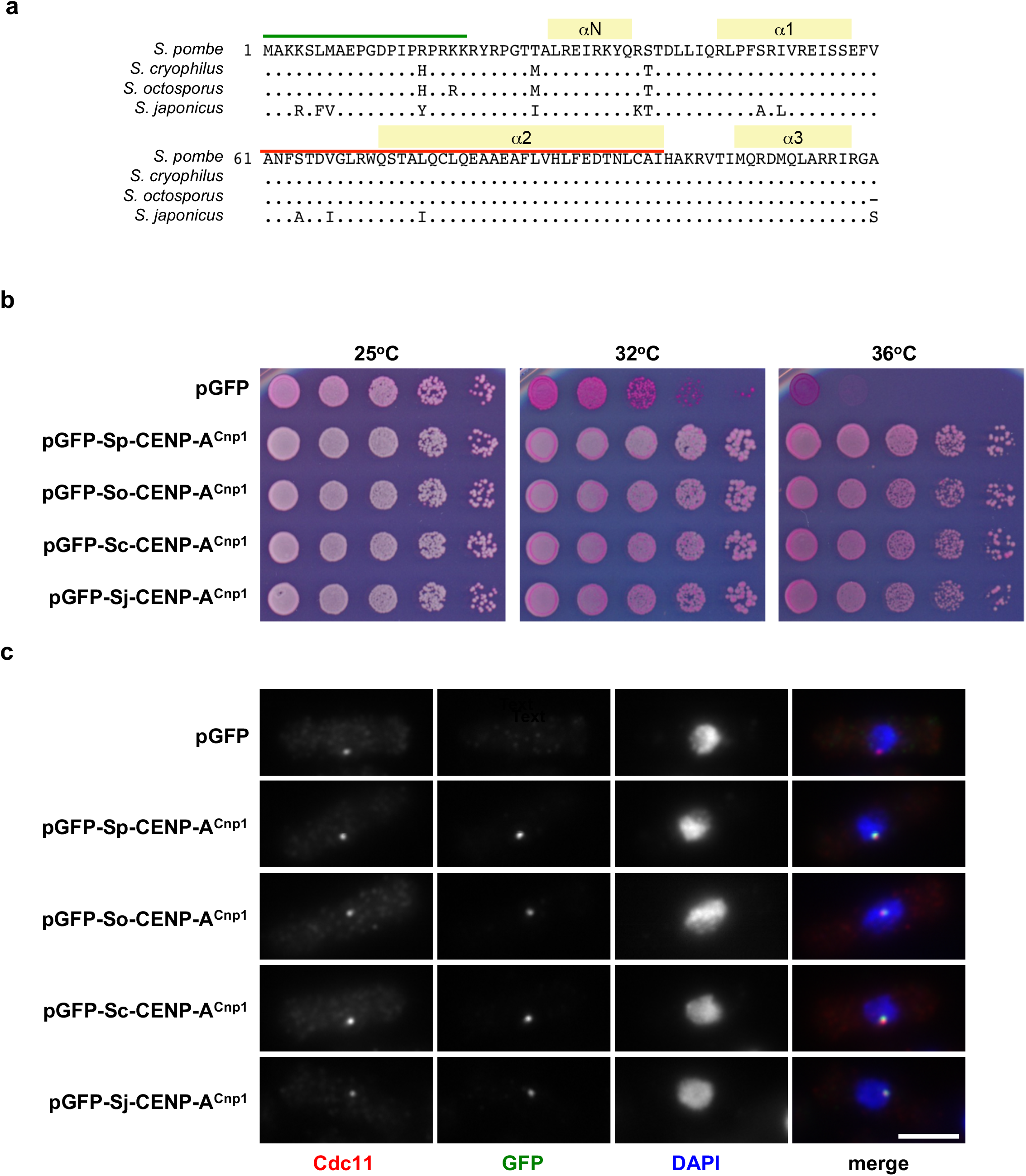
Cross-species functionality of CENP-A^Cnp1^ proteins. (a) Alignment of *Schizosaccharomyces* CENP-A^Cnp1^ proteins. Positions of alpha helices (yellow), N-terminal tail (green) and CENP-A-targeting domain (CATD; red) are indicated. (b) *S. pombe* temperature sensitive *cnp1-1* cells expressing plasmid-borne GFP-CENP-A^Cnp1^ from the indicated species (*Sp, S. pombe; So, S. octosporus; Sc, S. cryophilus; Sj, S. japonicus*), or GFP alone, were spotted on phloxine B-containing plates and incubated for 2-5 days at the indicated temperatures. (c) Localisation of GFP-tagged CENP-A^Cnp1^ from indicated *Schizosaccharomyces* species in *S. pombe*. Wild-type *S. pombe* cells bearing plasmids described in (a) were grown at 32°C before fixation and staining with anti-GFP (green), anti-Cdc11 (red, spindle-pole body) and DAPI (blue, DNA). Centromeres cluster at the spindle-pole body in *S. pombe*. Scale bar, 5 μm.

Introduction of *S. pombe* central-core (*S.pom-cnt*) DNA on minichromosomes into *S. pombe* results in the establishment and maintenance of CENP-A chromatin if *S.pom-cnt* is adjacent to heterochromatin, or if CENP-A is overexpressed^6,14,15,31^. *S.oct-cnt* regions (3.2-10 kb) or *S.pom-cnt2* (positive control) were placed adjacent to *S. pombe* outer-repeat DNA in mini-chromosome constructs (**Fig. 6a**) which were transformed into *S. pombe* cells expressing wild-type levels (wt-CENP-A) or overexpressing *S. pombe* GFP-CENP-A^Cnp1^ (hi-CENP-A^Cnp1^)^15^. Acquisition of centromere function is indicated by minichromosome retention on non-selective indicator plates (white/pale-pink colonies), and by the appearance of sectored colonies (**Fig. 6b,c**). The pHET-*S.pom-cnt2* minichromosome containing *S.pom-cnt2* established centromere function at high frequency immediately upon transformation in hi-CENP-A^Cnp1^ cells (90%) and at lower frequency in wt-CENP-A cells (15%; not shown). Centromere function was established on S.oct-cnt-containing minichromosomes in hi-CENP-A cells only (**Fig. 6d**). Centromere function was not due to minichromosomes gaining portions of *S. pombe* central-core DNA (data not shown). CENP-A^Cnp1^ ChIP-qPCR indicated that, for minichromosomes with established centromere function, CENP-A^Cnp1^ chromatin was assembled on non-homologous *S.oct-cnt* DNA, to levels similar to those at endogenous *S. pombe* centromeres and to *S.pom-cnt2* on a minichromosome (**Fig. 6e**). Minichromosomes containing *S.oct-cnt* provided efficient segregation function (**Fig. 6d**), no longer requiring CENP-A^Cnp1^ overexpression to maintain that function once established (**Fig. 6f**), consistent with the self-propagating ability of CENP-A chromatin^5,15^. These analyses indicate that *S.oct-cnt* is competent to establish CENP-A chromatin and centromere function in *S. pombe* when CENP-A^Cnp1^ is overexpressed, suggesting that *S. octosporus* central-core DNA has intrinsic properties that promote the establishment of CENP-A chromatin despite lacking sequence homology.

**Figure 6:**
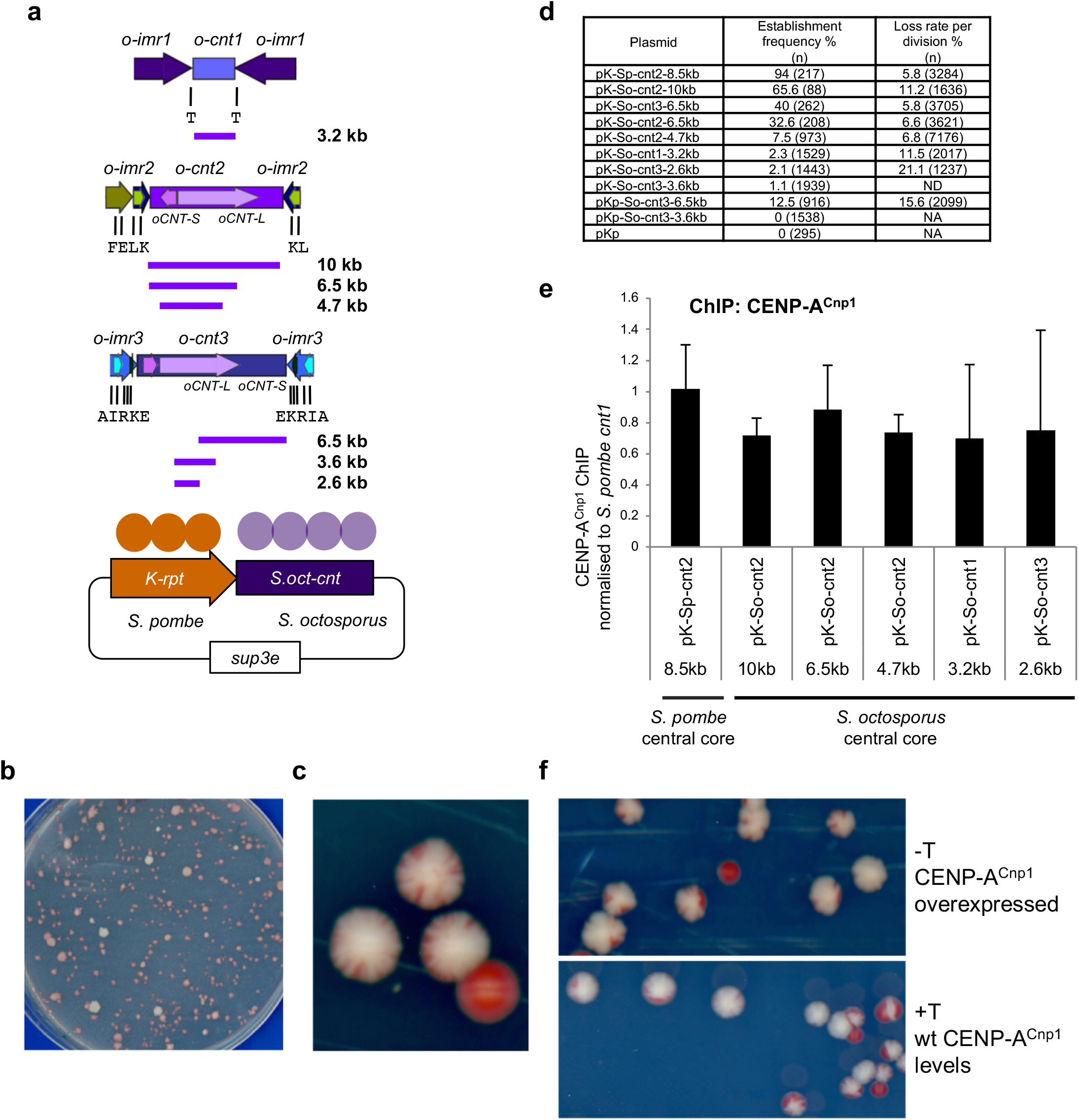
*S. octosporus* central core DNA establishes CENP-A^Cnp1^ chromatin upon introduction into *S. pombe*. (a) Indicated regions of *S. octosporus* central core DNA placed adjacent to a portion of *S. pombe* heterochromatin-forming outer repeat sequence on a plasmid. (b) *S. pombe* transformants containing minichromsome plasmids were replica-plated to low adenine non-selective plates: colonies retaining the chimeric minichromosome plasmid are white/pale-pink, those that lose it are red. Representative plate showing pKp-So-cnt3-6.5kb-containing colonies. (c) *S. pombe* cells containing pKp-So-cnt3-6.5kb chimeric minichromosome were streaked to single colonies. Red colour indicates loss of minichromosome; small red sectors indicate low frequency minichromosome loss and mitotic segregation function. (d) Establishment frequency of chimeric minichromosomes in *S. pombe* hi-CENP-A^Cnp1^ cells. Establishment frequency determined by replica plating of transformants (Methods) as shown in b (n=number of transformants analysed). Chromosome loss rate of established minichromosomes was determined by half-sector assay (Methods). Two transformants containing established centromeres were analysed for each minichromosome and the mean loss rate determined, n=number of colonies screened. (e) ChIP-qPCR for CENP-A^Cnp1^ on *S. pombe* hi-CENP-A^Cnp1^ cells containing chimeric minichromosomes with established centromere function. Three independent transfomants were analysed for each minichromosome. ChIP enrichment on *S.pom-cnt2* and S.oct-cnt-bearing minichromosomes is normalised to the level at endogenous *S. pombe cnt1*. Error bars, standard deviation. (f) Propagation of chimeric minichromosome stability. Cells containing pK(5.6kb)-So-cnt2-10kb were streaked on low adenine-containing plates with or without thiamine which results in repression or expression of high levels of *S. pombe* CENP-A^Cnp1^.

Based on conserved features, ancestral *Schizosaccharomyces* centromeres may have consisted of a CENP-A^Cnp1^-assembled central-core surrounded by tDNA clusters and 5S rDNAs. We surmise that RNAPIII promoters perhaps provided targets for transposon integration^32^, followed by heterochromatin formation to silence retrotransposons and preserve genome integrity^33,34^. The ability of heterochromatin to recruit cohesin^35,36^, benefitting chromosome segregation selected for heterochromatin maintenance^37,38^, rather than underlying sequence which evolved by repeat expansion and continuous homogenisation^18–20^. Because tDNAs performed important functions – as boundaries preventing heterochromatin spread into central-cores and perhaps in higher order centromere organisation and architecture – tDNA clusters were maintained^21^. In *S. pombe*, non-centromeric and centromeric tDNAs and 5S rDNAs cluster adjacent to centromeres in a TFIIIC-dependent manner^22,23^. The multiple tandem centromeric 5S rDNAs and tDNAs could contribute to a robust, highly-folded heterochromatin structure promoting optimum kinetochore configuration for co-ordinated microtubule attachments and accurate chromosome segregation^38^.

The lack of overt sequence conservation between centromeres of different species appears not to prevent functional conservation, which may be driven by underlying sequence features or properties such as the transcriptional landscape. Although maintenance of centromere function has been observed at a pre-established human centromere in chicken cells^39^ (310 My divergence), CENP-A establishment on human alpha-satellite in mouse cells^40^ (90 My divergence) is surpassed by the competence of *S. octosporus* central-core DNA to establish CENP-A chromatin in *S. pombe* from which it is separated by 119 My of evolution^16^ (equivalent to 383 My using a chordate molecular clock). Thus, our analyses extend the evolutionary timescale over which cross-species establishment of CENP-A chromatin has been demonstrated.

## Methods

### Cell growth and manipulation

Standard genetic and molecular techniques were followed. Fission yeast methods were as described^41^. Strains used in this study are listed in **Supplementary Table 20**. All *Schizosaccharomyces* strains were grown at 32°C in YES, except *S. cryophilus* which was grown at 25°C unless otherwise stated. *S. pombe* cells carrying minichromosomes were grown in PMG-ade-ura. For low GFP-tagged CENP-A^Cnp1^ protein expression from episomal plasmids, cells were grown in PMG-leu with thiamine.

### PacBio sequencing of genomic DNA

High molecular weight genomic DNA was prepared from *S. cryophilus, S. octosporus* and *S. japonicus* using a Qiagen Blood and Cell Culture DNA Kit (Qiagen), according to manufacturer’s instructions. Pacific Biosciences (PacBio) sequencing was carried out at the CSHL Cancer Center Next Generation Genomics Shared Resource. Samples were prepared following the standard 20 kb PacBio protocol. Briefly: 10-20 μg of genomic material was sheared via g-tube (Covaris) to 20 kb. Samples were damage repaired via ExoVII (PacBio), damage repair mix and end repair mix using standard PacBio 20 kb protocol. Repaired DNA underwent blunt-end ligation to add SMRTbell adapters. For some libraries: 10-50 kb molecules from 1-2 μg SMRTbell libraries were size selected using BluePippin (Sage Science) after which samples were annealed to Pacbio SMRTbell primers per the standard PacBio 20 kb protocol. Annealed samples were sequenced on the PacBio RSII instrument with P4/C3 chemistry. Magbead loading was used to load each sample at a concentration between 50 to 200 pM. Additional PacBio sequencing (without BluePippin) was performed by Biomedical Research Core Facilities, University of Michigan. There, the following kits were used: DNA Sequencing Kit XL 1.0, DNA Template Prep Kit 2.0 (3Kb - 10Kb)” and DNA/Polymerase Binding Kit P4. MagBead Standard Seq v2 sequencing was performed using 10,000 bp size bin with no Stage Start with a 2 hour observation time on a PacBio RSII sequencer. A summary of PacBio sequencing performed is listed in **Supplementary Table 21**.

### *De novo* whole genome assembly of PacBio sequence reads

PacBio reads were assembled using HGAP3 (The Hierarchical Genome Assembly Process version 3)^42^. Reads were first sorted by length, and the top 30% used as seed reads by HGAP3. All remaining reads of at least 1 kb in length were used to polish the seed reads. These polished reads were used to *de novo* assemble the genomes and Quiver software used to generate consensus genome contigs. Comparisons to the ChIP-seq input data and Broad Institute *Schizosaccharomyces* reference genomes^16^ showed very high agreement with these datasets. The *S. octosporus* and *S. cryophilus* chromosomes were named according to their sequence lengths, the longest chromosome being labelled as chromosome I in each case.

### *De novo* assembly of the *S. pombe* genome using nanopore technology

Genomic DNA was extracted as described previously^43^. DNA purity and concentration were assessed using a Nanodrop 2000 and the double-stranded high sensitivity assay on a Qubit fluorometer, respectively. Genomic DNA was sequenced using the MinION nanopore sequencer (Oxford Nanopore Technologies). Three sequencing libraries were generated using the 1D ligation kit SQK-LSK108, the 2D ligation kit SQK-NSK007 and the 1D Rapid sequencing kit SQK-RAD002, following manufacturers guidelines. Each library was sequenced on one MinION flow cell. Sequencing reads were base-called using Metrichor (1D and 2D ligation libraries) or Albacore (rapid sequencing library). The combined dataset incorporating reads from three flow cells was assembled using Canu v1.5. The assembly was computed using default Canu parameters and a genome size of 13.8 Mbp. QUAST v3.2 was used to evaluate the genome assembly.

### Genome annotation and chromosome structure

Genes were annotated onto the genome both *de novo*, using BLAST and the sequences of known genes, and by using liftover (https://genome-store.ucsc.edu) to carry over the previous gene annotation information from the Broad institute reference genomes (ref). CrossMap^44^ was then used to lift the chain files over to the new, updated genome. The locations of tDNAs were predicted using tRNAscan^45,46^. Dfam 2.0^47^ was used to annotate repetitive DNA elements. MUMmer3.23^48^ was used to compare the genomes and annotate repeat elements and tandem repeat sequences, including those located in centromeric domain and telomere sequences. Centromeric repeat elements were manually identified using BLASTN and MEGABLAST (https://blast.ncbi.nlm.nih.gov). Each repeat element was named according to their sequence features (association with tDNA & rDNAs) and locations. The sequence of the wild-type (h^90^) *S. pombe* mating-type locus was obtained by manually merging nanopore and PacBio contigs using available data^16^, Supplementary Figure 10 and information at *www.pombase.org/status/mating-type-region*. Genome synteny alignment analysis was carried out using syMAP42^49,50^, based on orthologous genes among the three genomes.

### ChIP-qPCR

For analysis of CENP-A^Cnp1^ association with minichromosomes bearing *S. octosporus* central core DNA, three independent transformants with established centromere function (indicated by ability to form sectored colonies) for each minichromosome were grown in PMG-ade-ura cultures and fixed with 3.7% formaldehyde for 15 min at room temperature. Cells were lysed by bead-beating (Biospec) and ChIP was performed as previously described^51^. 10 μl anti-CENP-A^Cnp1^ sheep antiserum and 25 μl Protein-G-Agarose beads (Roche) were used per ChIP. qPCR was performed using a LightCycler 480 and reagents (Roche) and analysed using Light Cycler 480 Software 1.5 (Roche). Primers used in qPCR are listed in **Supplementary Table 22**. Mean %IP ChIP values for *Sp-cnt* or *So-cnt* on minichromsomes were normalised to %IP for endogenous *S. pombe cnt1*. Error bars represent standard deviation.

### ChIP-seq

A modified ChIP protocol was used. Briefly, pellets containing 7.5 × 10^8^ cells were lysed by four 1 min pulses of bead beating in 500 μl of lysis buffer (50 mM HEPES-KOH, pH 7.5, 140 mM NaCl, 1 mM EDTA, 1% Triton X-100, 0.1% sodium deoxycholate), with resting on ice in between. The insoluble chromatin fraction was pelleted by centrifugation at 6000 *g* and washed with 1 ml lysis buffer before resuspension in 300 μl lysis buffer containing 0.2% SDS. Chromatin was sheared by sonication using a Bioruptor (Diagenode) for 30 minutes (30 s on/off, high setting). 900 μl of lysis buffer (no SDS) was added and samples clarified by centrifugation at 17000 *g* for 20 minutes and the supernatant used for ChIP. 6 μl anti-H3K9me2 mouse monoclonal mAb5.1.1^52^ (kind gift from Takeshi Urano) or 30 μl sheep anti-CENP-A^Cnp1^ antiserum^51^ were used, along with protein G-dynabeads (ThermoFisher Scientific) or Protein-G agarose beads (Roche), respectively. (For neocentromere strains, cells were first treated with Zymolyase 100T, washed in sorbitol and permeablized. Chromatin was fragmented with incubation with micrococcal nuclease. Cell suspensions were adjusted to standard ChIP buffer conditions and extracted chromatin was processed as per standard ChIP.) Immunoprecipitated DNA was recovered using Qiagen PCR purification columns. ChIP-Seq libraries were prepared with 1-5 ng of ChIP or 10 ng of input DNA. DNA was end-repaired using NEB Quick blunting kit (E1201L). The blunt, phosphorylated ends were treated with Klenow-exo^-^ (NEB, M0212S) and dATP. After ligation of NEXTflex adapters (Bioo Scientific) DNA was PCR amplified with Illumina primers for 12-15 cycles and library fragments of ~300 bp (insert plus adaptor sequences) were selected using Ampure XP beads (Beckman Coulter). The libraries were sequenced following Illumina HiSeq2000 work flow (or as indicated in **Supplementary Table 21**).

### Defining fission yeast centromeres

CENP-A^Cnp1^ and H3K9me2 ChIP-seq data was generated to identify centromere regions. ChIP-Seq reads with mapping qualities lower than 30, or read pairs that were over 500-nt or less than 100-nt apart, were discarded. ChIP-seq data was normalized with respect to input data. Paired-end ChIP-seq data (single-end for *S. japonicus*) was aligned to the updated genome sequences using Bowtie2^53^. Samtools^54^, Deeptools^55^ and IGV^56^ were subsequently used to generate sequence data coverage files and to visualize the data. MACS2^57^ was used to detect CENP-A^Cnp1^ and heterochromatin-enriched regions of the genome.

### Centromere tDNA cluster analysis

To test for the enrichment of tDNA clusters at centromere regions a greedy search approach was used to identify potential clusters. All tDNAs less than 1000 bp apart were grouped into clusters. To test for significant clustering of tDNAs at the centromere the locations of tDNAs across the genome were shuffled 1000 times. For each cluster observed in the real genome the proportion of permutations where the same cluster was observed at least as many times was calculated to provide estimates of significance. Following conversion of these p-values to q values to account for multiple testing, the centromere tDNA clusters each exhibited a q-value less than 0.005.

### Hsp16 gene tree analysis

*hsp16* paralogs from *S. octosporus* and *S. cryophilus* genomes were predicted using BLASTP. The predicted protein sequences from *hsp16* genes across all four fission yeasts were aligned together with those from *S. cerevisiae* using Clustal Omega. BEAST (Bayesian Evolutionary Analysis Sampling Trees)^58^ and FigTree (http://tree.bio.ed.ac.uk/software/figtree/) was used to generate and view the *hsp16* gene phylogenetic tree.

### 5-mer frequency PCA analysis

The CENP-A^Cnp1^-associated sequences in the *S. pombe, S. cryophilus* and *S. octosporus* genomes are all approximately 12 kb in length. Each genome was therefore split into 12 kb sliding windows with a 4.5 kb overlap. The frequencies of each 5-mer was calculated in each window using Jellyfish^59^. CENP-A^Cnp1^-associated regions showed a general enrichment of AT base pairs relative to the genome as a whole. To normalize for GC content amongst the windows, all base pairs were randomized in each sequence window to generate 1000 artificial sequences with the same GC content. 5-mer frequencies were then recalculated for each of these 1000 artificial sequences and the true original 5-mer frequencies compared to these background frequencies by calculating a z-score. Consequently, these enrichment scores represent the k-mer enrichments in a given sequence normalized for GC content. Genome windows were split into 6 groups: CENP-A^Cnp1^-associated sequences (CENP-A^Cnp1^ peaks covering more than 6 kb of sequence); outer repeat heterochromatin regions (more than half the window covered by H3K9me2 peaks adjacent to CENP-A domains); sub-telomeric regions (more than half the window covered by H3K9me peaks and close to the end of a chromosome); Mating-type locus, neo-centromere regions (identified using CENP-A^Cnp1^ ChIP-seq data on *S. pombe* neocentromere-containing strains^29^) and remaining genome sequences. Logistic regression and mean comparison were used to determine whether principal components were linked to the probability of a sequence belonging to a particular sequence group^60^. Logistic regression and mean comparison were used to determine whether principal components (FactoMineR) were linked to the probability of a sequence belonging to a particular sequence group.

### Construction of minichromosomes

Regions of *S. octosporus* central core regions were amplified with primers indicated in **Supplementary Table 22**. Fragments were digested with *Bgl*II, *Nco*l or *Bam*HI, *Nco*l and ligated into *Bgl*II-*Nco*l-digested plasmid pK(5.6kb)-MCS-ΔBam which contains a 5.6 kb fragment of the *S. pombe* K (*dg*) outer repeat. To create plasmid pK-So-cnt2-10kb, an additional 3.6 kb region from *S.oct-cnt2* was inserted as a *Bam*HI-*Sal*I fragment into *Bgl*II-*Sal*I-digested pK-So-cnt2-6.5kb to make a 10 kb region of *S. octosporus* central core. For pKp plasmids, *S. octosporus* central core regions were by inserted as *Bgl*II-*Nco*l or S*al*I-*Bam*HI fragments into *Sal*I-*Bam*HI or NcoI-BamHI digested plasmid pKp (pMC91) which contains 2 kb region from *S. pombe* K(*dg*) outer repeat. Plasmids are listed in **Supplementary Table 23**.

### Centromere establishment assay

Strains A7373 or A7408, which contains integrated nmt41-GFP-CENP-A^Cnp1^ to allow high level expression of CENP-A^15^,were grown in PMG-complete medium and transformed using sorbitol-electroporation method^61^. Cells were plated on PMG-uracil-adenine plates and incubated at 32°C for 5-10 days until medium-sized colonies had grown. Colonies were replica-plated to PMG low adenine (10 μg/ml) plates to determine the frequency of establishment of centromere function. These indicator plates allow minichromosome loss (red) or retention (white/pale pink) to be determined. Minichromosome retention indicates that centromere function has been established and that minichromosomes segregate efficiently in mitosis. In the absence of centromere establishment, minichromosomes behave as episomes that are rapidly lost. Minichromosomes occasionally integrate giving a false positive white phenotype. To assess the frequency of such integration events and to confirm establishment of centromere segregation function, a proportion of colonies giving the white/pale-pink phenotype upon replica plating were re-streaked to single colonies on low-adenine plates – sectored colonies are indicative of segregation function with low levels of minichromosome loss, whereas pure white colonies are indicative of integration into endogenous chromosomes – and the establishment frequency adjusted accordingly.

### Minichromosome stability assay

Minichromosome loss frequency was determined by half-sector assay. Briefly, transformants containing minichromsomes with established centromere function were grown in PMG-ade-ura to select for cells containing the minichromosome. Two transformants were analysed per minichromosome (four for pK-So-cnt2-4.7kb). Cells were plated on low-adenine containing plates and allowed to grow non-selectively for 4-7 days. Minichromosome loss is indicated by red sectors and retention by white sectors. To determine loss rate per division, all colonies were examined with a dissecting microscope. All colonies – except pure reds – were counted to give total number of colonies. Pure reds were checked for the absence of white sectors and were excluded because they had lost the minichromosome before plating. To determine colonies that lost the minichromosome in the first division after plating, ‘half-sectored’ colonies were counted. This included any colony that was 50% or greater red (including those with only a tiny white sector). Loss rate per division is calculated as the number of half-sectored colonies as a percentage of all (non-pure-red) colonies.

### Immunolocalisation

For localisation of CENP-A^Cnp1^, *Schizosaccharomyces* cultures were fixed with 3.7% formaldehyde for 7 min, before processing for immunofluorescence as described^51^. Anti-CENP-A^Cnp1^ sheep antiserum^51^ (raised to the N-terminal 19 amino-acids of *S. pombe* CENP-A^Cnp1^) was used at 1:1000 dilution, and Alexa-488-coupled donkey-anti-sheep secondary antibody (A11015; Invitrogen) at 1:1000 dilution. Cells were stained with DAPI and mounted in Vectashield. Microscopy was performed with a Zeiss Imaging 2 microscope (Zeiss) using a 100x 1.4NA Plan-Apochromat objective, Prior filter wheel, illumination by HBO100 mercury bulb. Image acquisition with a Photometrics Prime sCMOS camera (Photometrics, https://www.photometrics.com) was controlled using Metamorph software (Universal Imaging Corporation). Exposures were 1500 ms for FITC/Alexa-488 channel and 300-1000 ms for DAPI. Images shown in Figure 2a are autoscaled.

To express GFP-tagged versions of *Schizosaccharomyces* CENP-A^Cnp1^ proteins in *S. pombe*, ORFs were amplified from relevant genomic DNA using primers listed in **Supplementary Table 22**. Fragments were digested with *Nde*l-*Bam*HI or *Nde*l-*Bgl*II and ligated into *Nde*l-*Bam*HI digested pREP41X-GFP vector^62^ (**Supplementary Table 23**). For detection of GFP-tagged versions of *Schizosaccharomyces* CENP-A^Cnp1^ proteins in *S. pombe*, cells containing pREP41X-GFP-CENP-A^Cnp1^ episomal plasmids (variable copy number) were grown in PMG-leu + thiamine to allow low GFP-CENP-A^Cnp1^ expression. Cells were fixed, processed for immunolocalisation and microscopy as above. Anti-GFP antibody (A11122; Invitrogen) was used at 1:300, anti-Cdc11^51^ (a spindle-pole body marker; gift from Ken Sawin) was used at 1:600. Secondary antibodies were, respectively, Alexa-488 coupled chicken anti-rabbit (A21441; Invitrogen) and Alexa-594 coupled donkey anti-sheep (A11016; Invitrogen) both at 1:1000. Exposures were FITC/488 channel: 1500 ms, TRITC/594 1000 ms, DAPI 500-1000 ms. For display of images in Figure 5C, TRITC/594 and FITC/488 images are scaled relative to the maximum intensity in the set of images, whilst DAPI images are autoscaled.

## Data Availability

All data generated in this study have been submitted to GEO under accession number: GSE112454. SRA submission number for *S. pombe* nanopore sequencing data: SUB3761672. The following figures have associated raw data: 1, 2, S1, S2, S3, S5.

## Acknowledgments

We thank Alastair Kerr, Shaun Webb and Daniel Robertson for bioinformatics support, David Kelly for microscopy support, and Ken Sawin and Takeshi Urano for antibodies, and Kojiro Ishii, Ken Sawin and Nick Rhind for yeast strains. We thank Robert Lyons, Joe Washburn, Christina McHenry (University of Michigan) and Greg J. Hannon, Richard McCombie, Eric Antoniou and Sara Goodwin (CSHL) for PacBio sequencing. We are grateful to Chris Ponting for advice and comments on the manuscript and Sandra Catania and other members of the Allshire and Heun labs for helpful discussions. N.R.T.T., R.A. and J.T-G. were supported by the Darwin Trust of Edinburgh. The Darwin Trust and a Principal’s Career Development scholarship supported N.R.T.T. P.T. was partly supported by funding from the European Commission Network of Excellence EpiGeneSys-(HEALTH-F4-2010-257082) and a Wellcome Enhancement Award (095021) to R.C.A. R.C.A. is a Wellcome Principal Research Fellow (095021, 200885); the Wellcome Centre for Cell Biology is supported by core funding from Wellcome (203149). C.A.M. and C.A.N. are supported by Biotechnology and Biological Sciences Research Council (BBSRC) grant BB/N016858/1 and Wellcome Investigator Award 110064/Z/15/Z.Pacific Biosciences (PacBio) sequencing carried out at the CSHL Cancer Center Next Generation Genomics Shared Resource, which is supported by the Cancer Center Support Grant 5P30CA045508 was paid for by a kind gift from Kathryn W. Davis to GJH.

## Author Contributions

R.C.A. and A.L.P. designed the study. P.T. performed the PacBio genome assemblies and bioinformatics, ChIP-seq analysis and PCA analysis. C.M. performed the nanopore sequencing of *S. pombe* supervised by C.N. H.B., N.R.T.T. J.T.-G. and R.A. generated ChIP-seq data with contribution from M.S. A.L.P. performed cytology, analysis of repetitive regions, and experiments on cross-species functionality. R.C.A. supervised the study. A.L.P. wrote the manuscript with contributions from P.T., R.C.A. and other authors. All authors read and approved the final version of the manuscript.

## Supplementary Figure Legends

**Figure S1:**
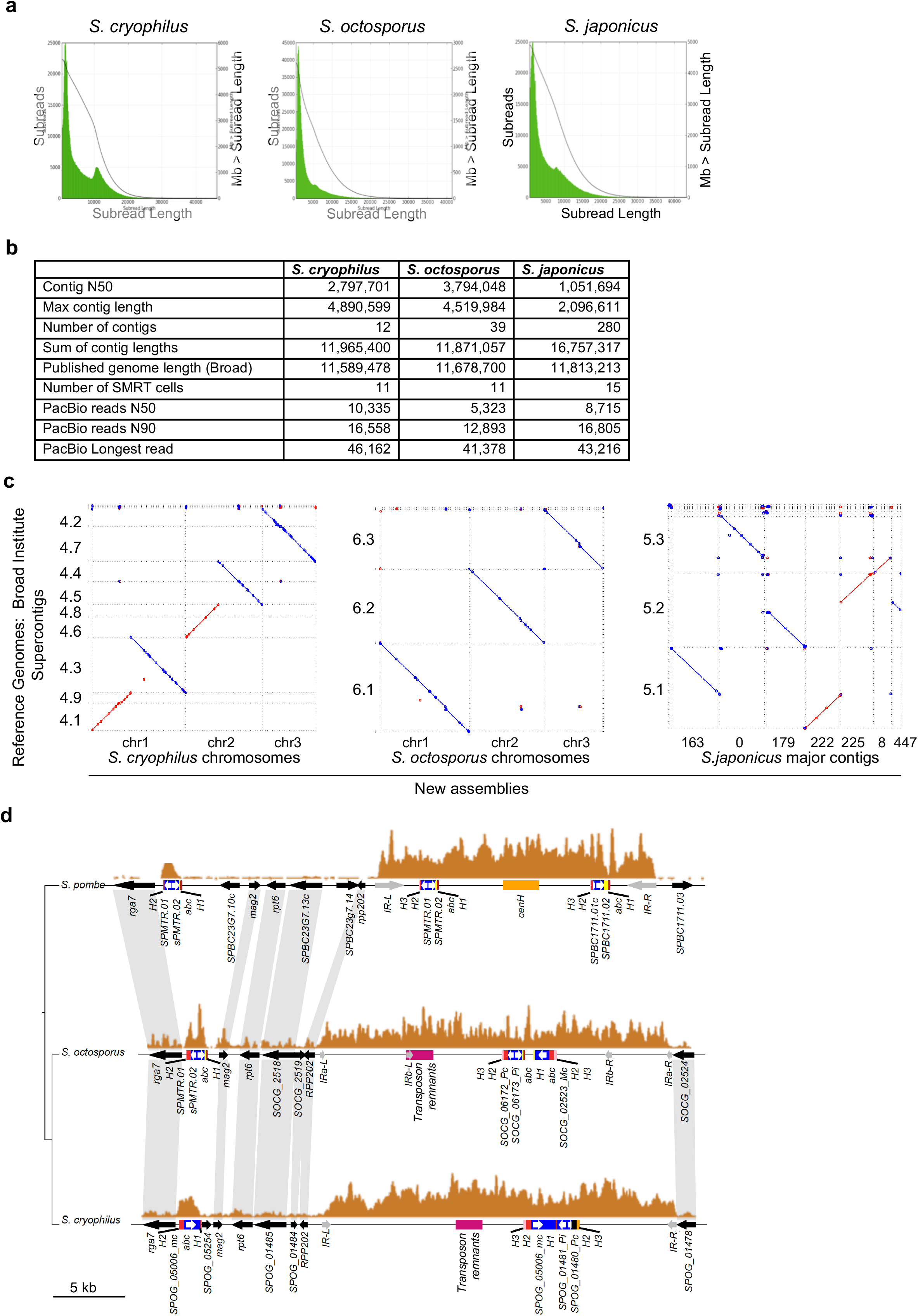
*S. octosporus* and *S. cryophilus* genome assembly statistics. a) Histograms of SMRT cell subread lengths (green) and the sum of subread length (black) for the indicated genomes. b) Summary of PacBio subreads and final assemblies. c) Dot plot comparison of new assemblies with previously published assemblies for *S. octosporus, S. cryophilus and S. japonicus*^16^. d) Organisation of mating-type loci in *S. pombe, S. octosporus* and *S. cryophilus*^16,22^. ChIP-seq profiles for H3K9me2-heterochromatin (orange) are shown. Positions of mating-type loci (blue) and mating-type genes (white), mating-type associated repeat elements (H1: dark red; H2: red; H3: pink; abc: yellow); transposon remnants (pink), inverted IR repeats (grey) and other genes (black) are indicated. *cenH* region (orange) homologous to *S. pombe* centromeric *dg/dh* repeats is indicated. Blue shading indicates homologous genes between species.

**Figure S2:**
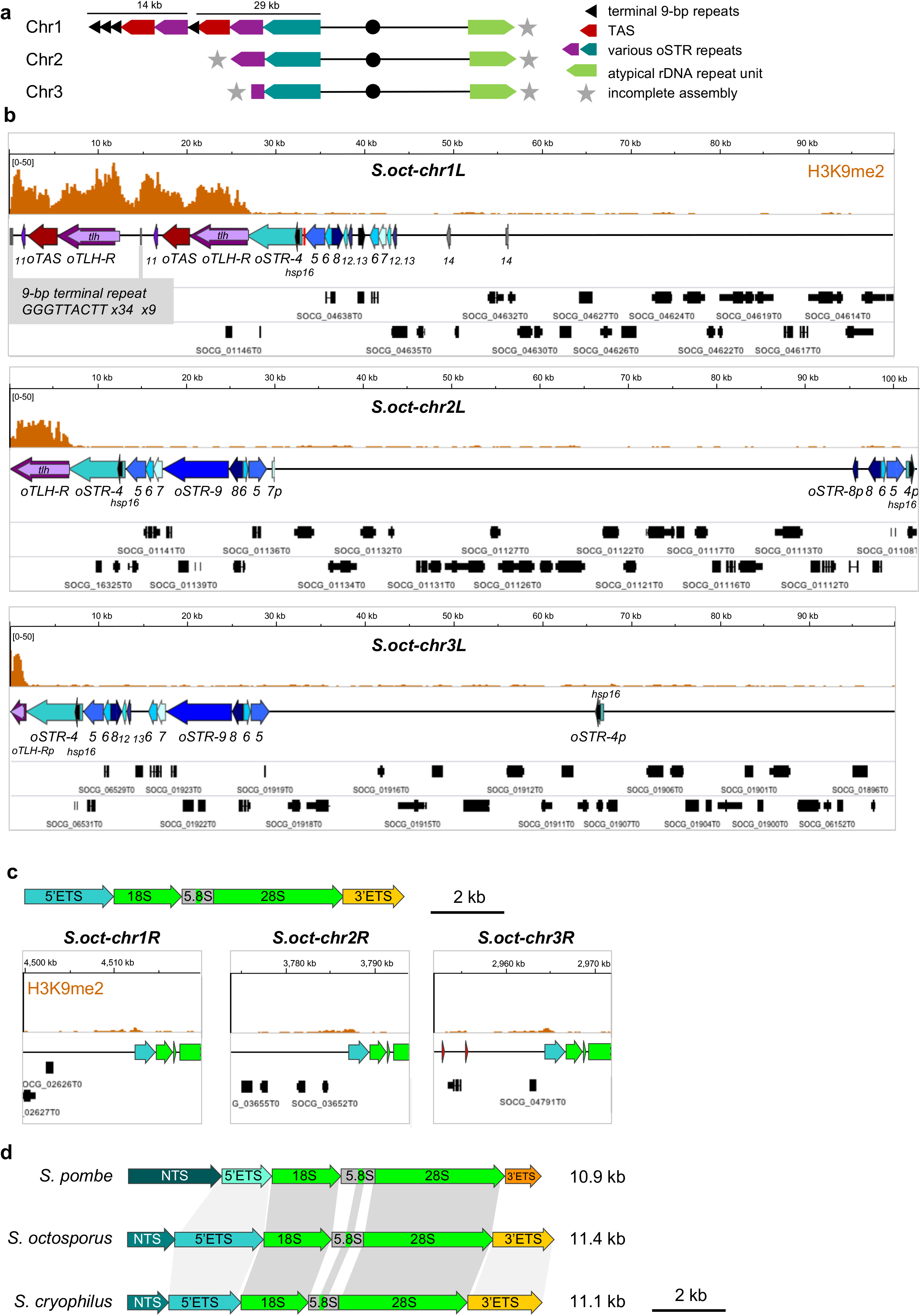
Structure of *S. octosporus* chromosome ends. a) Overview of *S. octosporus* chromosomes, indicating organisation of subtelomeres. Multiple copies of terminal repeats (black arrows; GGGTTACTT) are detected at the end of chr1L (and internally). Combinations of subtelomeric repeats, including telomere-associated sequences (oTAS, dark red); RecQ type DNA helicase genes (*tlh*) and associated repeats (oTLH-R) and other subtelomeric repeats (oSTR-4 etc; turquoise); details in (b). A partial atypical rDNA repeat is detected at: chr1R, 2R and 3R (light green arrow). Due to repetitive nature of this region, assemblies are incomplete at all chromosome ends (denoted by grey star), except chr1L. b) Details of terminal 100 kb of chr1L, chr2L and chr3L. Two copies of telomere-associated sequences (oTAS; dark red), and oTLH-R (containing RecQ type DNA helicase genes (*tlh*)) are present at chr1L, along with multiple copies of GGGTTACTT repeats. Numerous other subtelomeric repeats, designated oSTR-4 etc (blue/turquoise) are indicated, mostly by number designation only due to space constraints. c) Top, structure of atypical rDNA repeat unit (lacking the full NTS seen in typical rDNA repeats, see (d). Atypical rDNA repeat units are present at centromere-proximal side of chromosome ends: chr1R, 2R, 3R. Due to repetitive nature of these regions, full assembly was not achieved and the number of rDNA repeat units present at each chromosome end is unknown. From ChIP input read counts the total number of rDNA repeat units is estimated to be approximately 150 copies. d) Homology of rDNA repeat unit between *Schizosaccharomyces* species. *S. pombe, S. cryophilus* and *S. octosporus* rDNA repeats are shown. *S. pombe* elements were previously defined^63^. Homology indicated by grey blocks: darker grey indicates higher homology (65%-92%).

**Figure S3:**
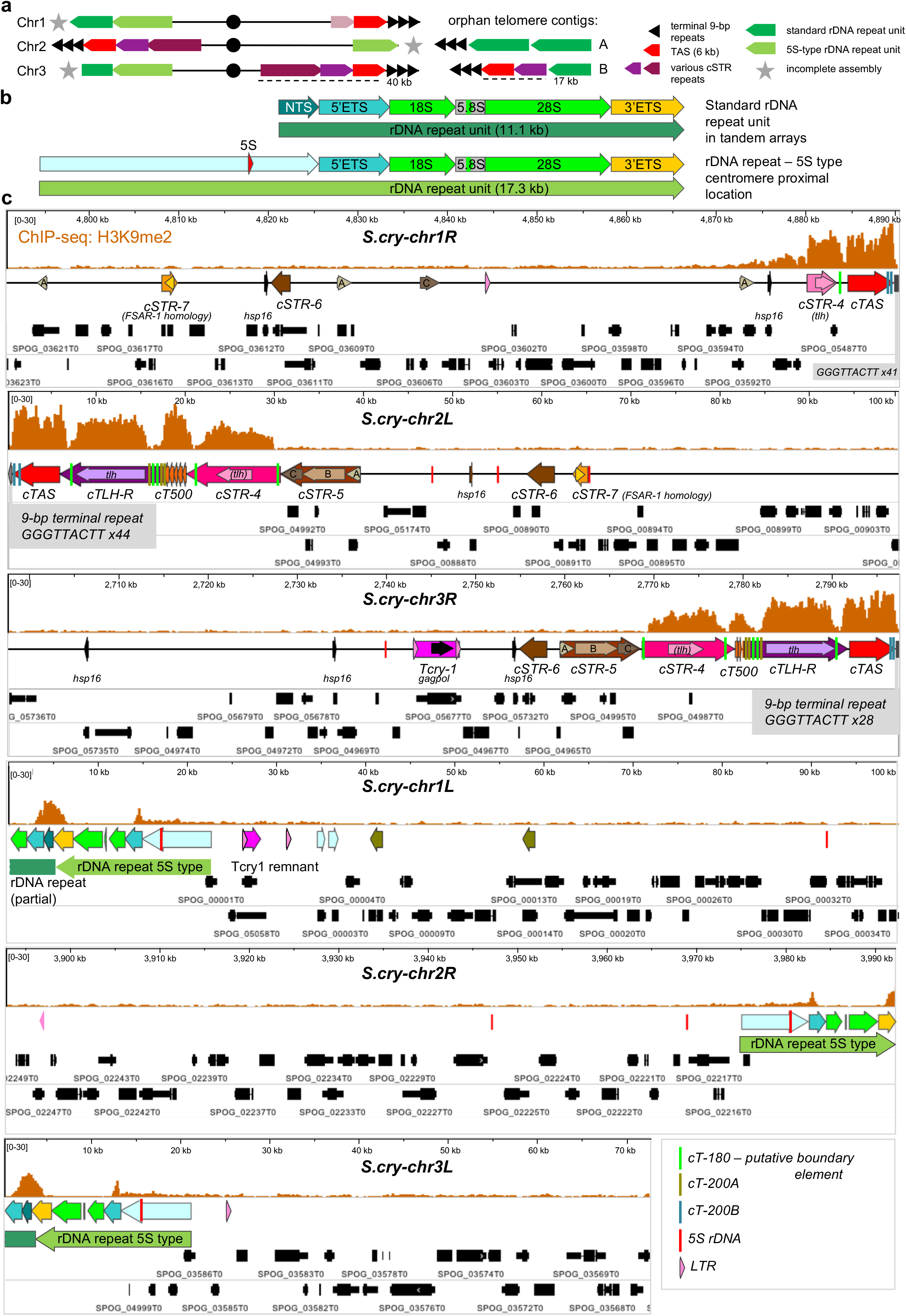
Structure of *S. cryophilus* chromosome ends. a) *Left*, overview of *S. cryophilus* chromosomes, indicating organisation of subtelomeres. Multiple copies of terminal repeats (black arrows; GGGTTACTT) are present at the ends of 1R, 2L and 3R, along with combinations of subtelomeric repeats, including telomere-associated sequences (cTAS, red); RecQ type DNA helicase genes (*tlh*) and associated repeats (cTLH-R) and other subtelomeric repeats (cSTR-4 etc; shades of pink/brown); details in c. rDNA repeats are located at: 1L, 2R and 3L. Centromere-proximal rDNA repeat is atypical (light green arrow) and associated with a 5S rDNA (details in b). Distal to that, assemblies of chromosome 1 and 3 indicate a partial standard rDNA repeat (no associated 5S rDNA; dark green). Due to repetitive nature of this region, assemblies are incomplete at 1L, 2R and 3L (denoted by grey star). *Middle*, two classes of terminal rDNA-containing contigs were also identified. Both types contain multiple copies of the terminal GGGTTACTT repeat. In class A these directly abut rDNA repeat units. In class B, cTAS and cTLH-R elements are located between the terminal repeats and rDNA repeat unit. A similar arrangement has recently been described for the rDNA-containing ends of *S. pombe* chromosome 3^64^. *Right*, key to repeat elements. b) Two types of rDNA repeat are present in *S. cryophilus. Top*, standard repeat of 11.1 kb, present in tandem arrays. 28S, 18S and 5.8S genes are indicated (green), along with putative associated elements external transcribed spacers (5’ETS, turquoise; 3’ETS, yellow) and non-transcribed spacer (NTS, teal). *Bottom*, atypical rDNA repeat located at centromere-proximal location of all three rDNA-containing chromosome ends. In place of standard NTS element it is associated with a 7.5 kb repeat (pale blue) containing a 5S rDNA gene (red arrow). c) *Upper 3 panels*: Subtelomere-repeat-containing chromosome ends are assembled in heterochromatin. Terminal 100 kb of arms 1R, 2L and 3R are shown. Location of repeat elements are indicated, along with positions of *hsp16* genes. Smaller repeat elements are indicated by vertical bars (see key, bottom right). H3K9me2 ChIP-seq profile is shown (orange). cT-180 repeats (green bars) coincide with deep troughs in H3K9me2 reads, suggesting that these elements could perform a boundary function. cTLH-R contains intact *tlh* genes, whereas cSTR-4 elements contain degraded copies of *tlh*. The partial cSTR-4 element has weak homology to cSTR-4 at 2L and 3R and a highly degraded copy of *tlh*. Location of genes indicated in black. The non-heterochromatic portion of the subtelomeres contain several paralogous genes, homologues of which are also found in the subtelomeric regions of *S. octosporus*. The intact retrotransposon *Tcry1* (magenta; LTRs, pink arrows) is located in the subtelomeric region of 3R. *Lower 3 panels*: Chromosome ends with rDNA repeat units: terminal 100 kb of 1L, 2R, 3L shown. Feature colours as in a, b. H3K9me2 ChIP-seq profile (orange) indicates enrichment over the nontranscribed spacer. Note that, as with all repetitive regions, numbers of ChIP-seq reads mapped represent an average over all repeats. Locations of 5S rDNAs, LTRs (pink arrow) and a partial *Tcry1* retrotransposon (magenta) are indicated. Assembly of full rDNA-containing chromosome ends was not possible due to its repetitive nature, consequently the number of rDNA repeat units present at each chromosome end remains unknown. However, the total number of rDNA repeat units is estimated to be 150 from ChIP input read counts.

**Figure S4:**
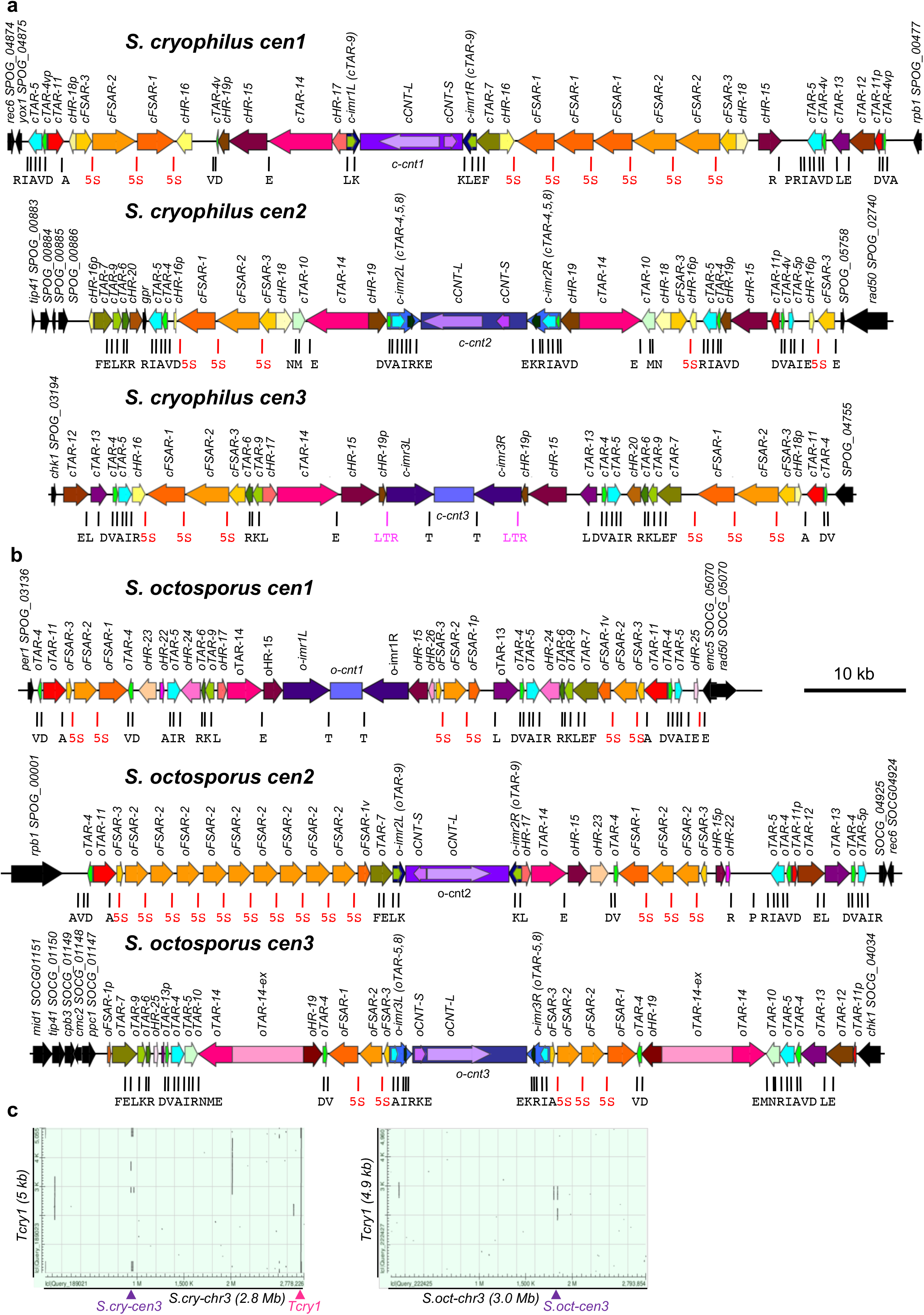
Centromere repeat organisation in *S. cryophilus* and *S. octosporus*. Structure and organisation of *S. cryophilus* (a) and *S. octosporus* (b) centromeres are shown. Location and names of centromeric repeats indicated. Repeat colour indicates identity/high degree of homology within species. Repeats with the same colour between species do not show sequence homology, but are present in similar contexts with respect to chromatin status, association with particular tDNAs or occurrence and location within centromeres. The only detectable homology is between tDNAs themselves, and between cTAR-14 (pink; all *S.cry-cens*) and extended oTAR-14-ex (*S.oct-chr3*) elements which have weak homology to the retrotransposon *Tcry1* and retransposon remnants at the mating-type loci of both species and at other locations in the genomes (see **Supplementary Tables 14,15,19**). Central core regions are indicated in shades of purple, and positions of long (*cCNT-L* and *oCNT-L*) and short (*cCNT-S* and *oCNT-S*) repeats are indicated. Innermost (*imr*) inverted repeats are shown in shades of blue and in some cases contain smaller ‘boundary type’ repeat elements (greens). tDNAs are shown as vertical black bars and the cognate amino acid shown in single letter code below. Small repeat elements associated with clustered tDNAs which may have boundary function are shown in shades of green and turquoise (tDNA-associated repeats; TARs). 5S rDNAs are shown as vertical red bars. Heterochromatic Five-S-associated repeats (cFSARs and oFSARs) are shown in shades of orange (see **Figure 3a**). Longer repeats associated with single tDNAs (unlikely to have boundary function) are indicated in shades of red/brown/plum (TARs 11-14). Other heterochromatic repeats (HR) are indicated in shades of yellow/brown/pink. Genes flanking the centromeres are indicated in black. (c) Retrotransposon remnants are present within *S. cryophilus* and *S. octosporus* centromeres and other genomic locations. *Left panel*: Dot-plot alignment (BLASTN) of *Tcry1* retrotransposon^16^ with *S. cryophilus* chromosome 3. Pink triangle indicates position of *Tcry1* itself at subtelomeric region of chr3R. Purple triangle indicates position of *S.oct-cen3*. Homology lies within cTAR-14 elements (also present at *S. cryophilus cen1* and *cen3*). Other regions of homology on chromosome arms are due to partial copies of *Tcry1* retrotransposon. *Right panel:* Dot-plot alignment (BLASTN) of *Tcry1* retrotransposon with *S. octosporus* chromosome 3. Purple triangle indicates position of *S.oct-cen3*. Homology within centromere is with oTAR-14ex elements, present only at *S.oct-cen3*. Region of homology in subtelomere is ~135 kb from chromosome end.

**Figure S5:**
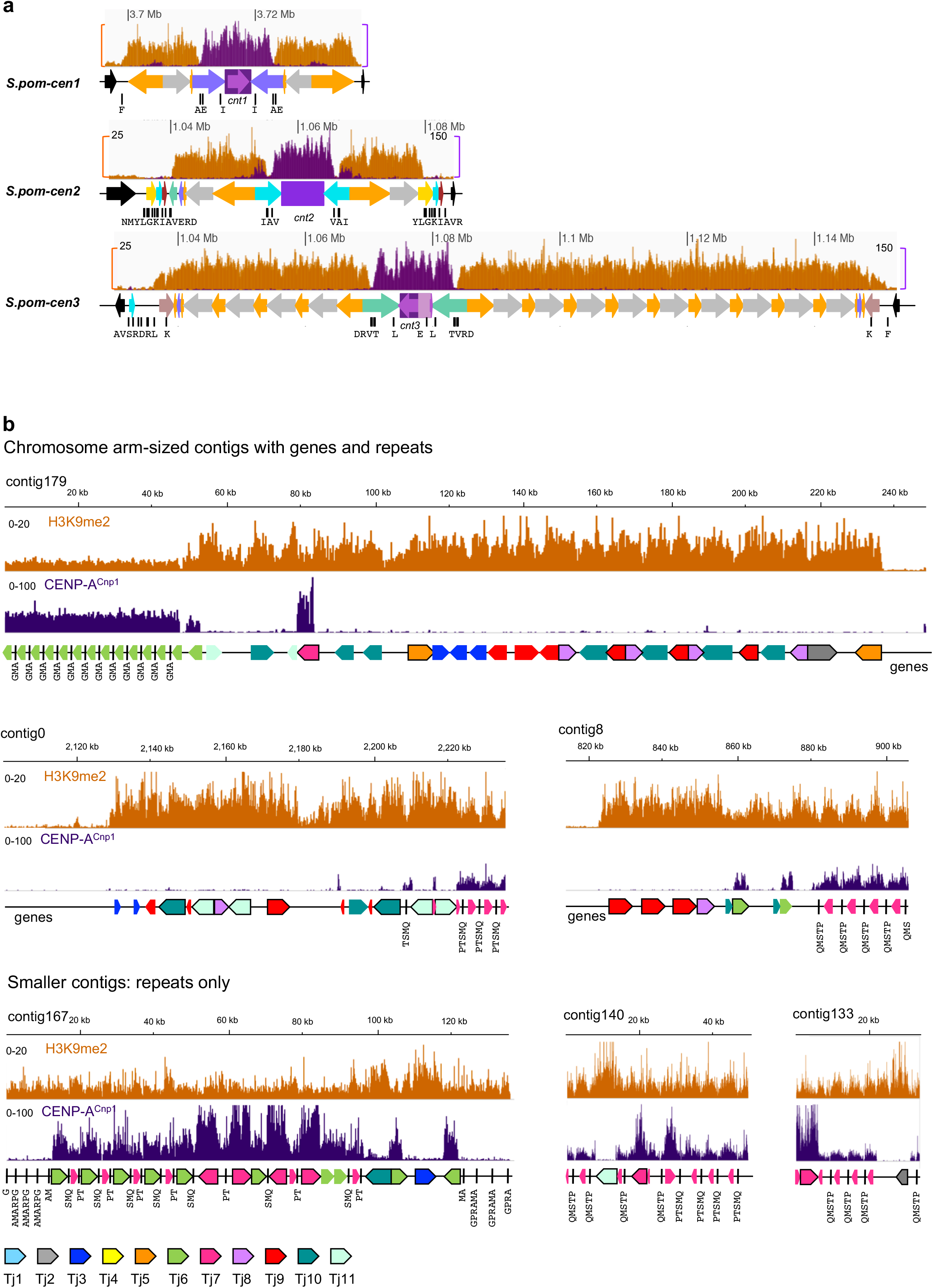
Domain organisation of *S. pombe* centromeres and transposon-rich centromeres of *S. japonicus*. (a) *S. pombe* genome was assembled from Oxford nanopore sequencing. Updated *S. pombe* centromere organisation indicating repeat elements. ChIP-seq profiles for CENP-A^Cnp1^ (purple) and H3K9me2-heterochromatin (orange) are shown above each centromere. Positions of tDNAs (single-letter code of cognate amino acid; black) are indicated. Central cores are shown in shades of purple (TM element common to *cnt1* and *cnt3* shown in mauve); innermost repeats (*imr*) in blues, heterochromatic outer repeat elements are shown in grey (dg) and orange (dh). (b) Representative H3K9me2 and CENP-A-associated *S. japonicus* contigs. ChIP-seq profiles for CENP-A^Cnp1^ (purple) and H3K9me2-heterochromatin (orange) are shown above each contig. Due to their repetitive nature, full assembly of centromere regions was not possible; association with CENP-A^Cnp1^ is strong indicator that a contig is centromere located. Retrotransposons mapping to contigs are indicated; colour-coding as previously published^16^ (key, bottom), full-length or almost full-length retrotransposons are indicated by a black outline. An additional putative retrotransposon was identified which has been named Tj11 (see **Supplementary Table 9**). Positions of tDNA clusters (single-letter code of cognate amino acids; black) are indicated. Top/middle: Chromosome arm-sized contigs which terminate within retrotransposon arrays. Bottom: smaller contigs containing only retrotransposons and other repetitive elements could not be incorporated into genome assembly.

**Figure S6:**
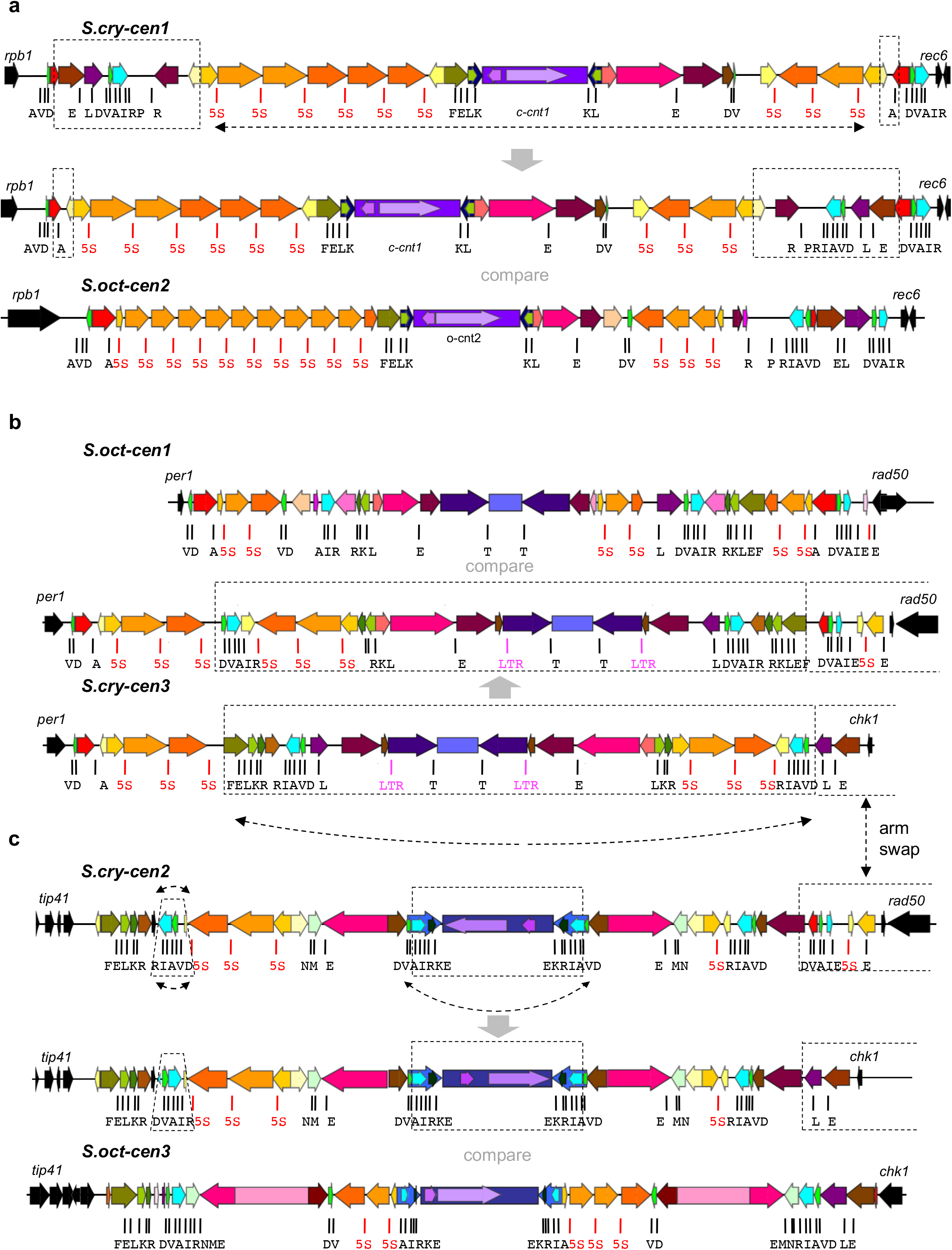
Synteny of tDNA clusters and organisation of repeat elements suggests common ancestry of *S. cryophilus* and *S. octosporus* centromeres, despite lack of sequence conservation. (a) Centromeres of *S. cryophilus* and *S. octosporus* paired with the most similar centromere from the opposite species. In silico rearrangements of *S. cryophilus* centromeres closely recapitulate the organisation of *S. octosporus* centromeres, providing support for their evolution from common ancestral centromeres. Dashed boxes and arrows indicate rearrangement that that would increase the structural similarity between *S.cry-cen1* and *S.oct-cen2*. b and c) A rearrangement involving arm swap of *S.cry-cen2R* and *S.cry-cen3L* would produce synteny of genes on either side of *S.cry-cen2* and *S.cry-cen3*, and *Soct-cen3* and *Soct-cen1* respectively (centre, dashed double-headed arrow). Partial inversions (curved double-headed arrows) would increase similarity between *S. cryophilus* and *S. octosporus* centromeres (labelled compare). See **Figure 4a**.

---

**Supplementary Table 1:**
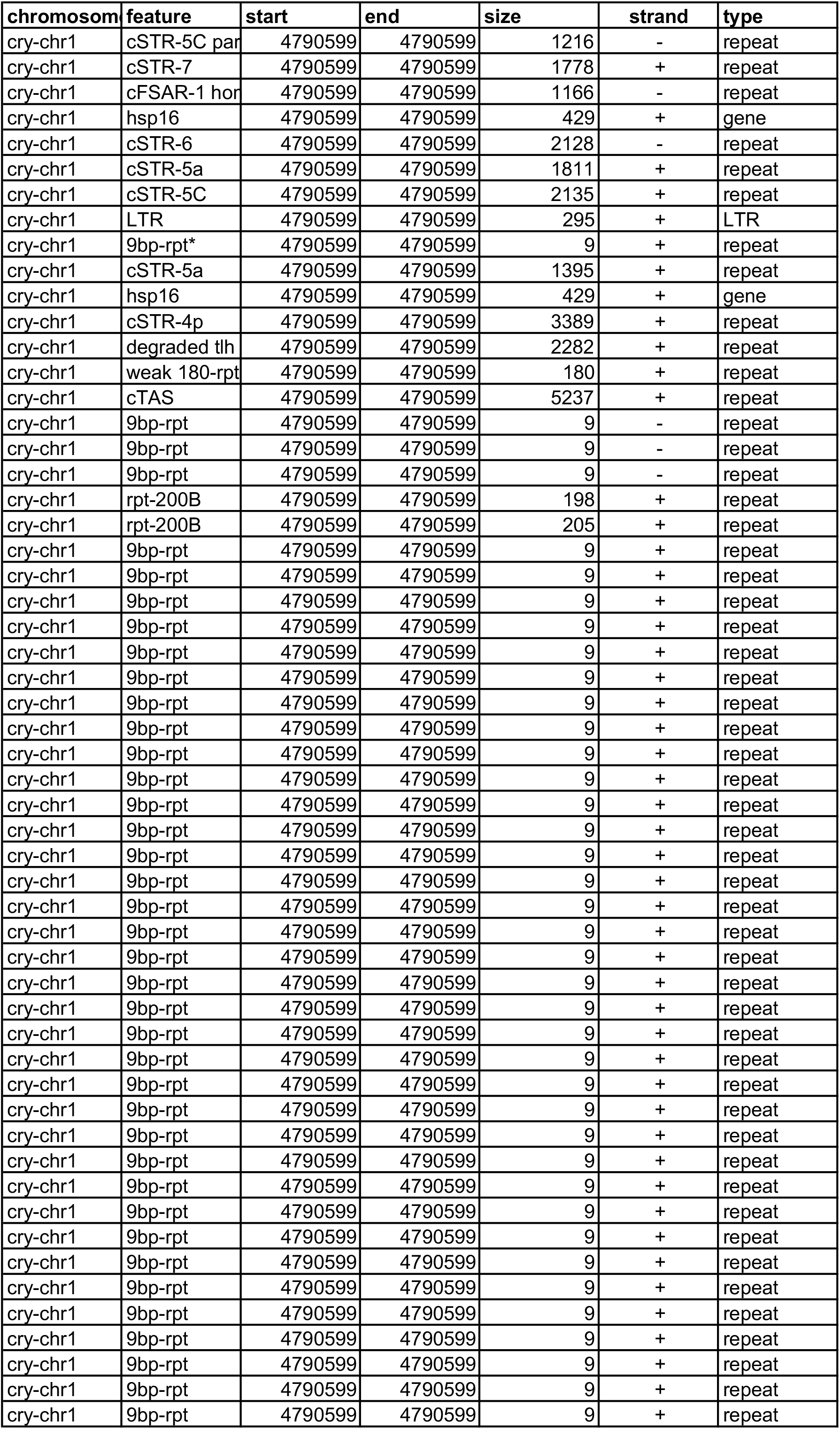
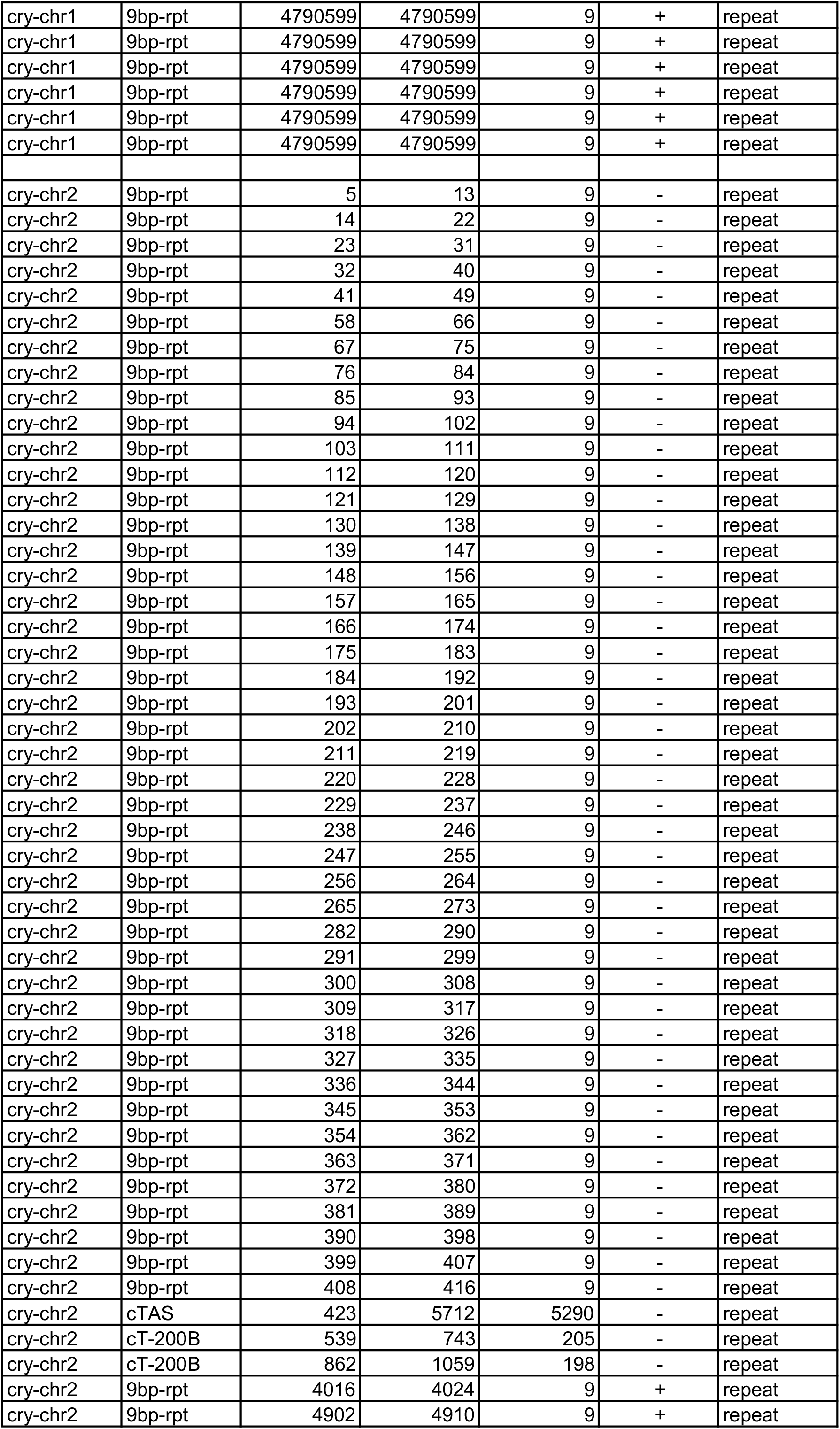
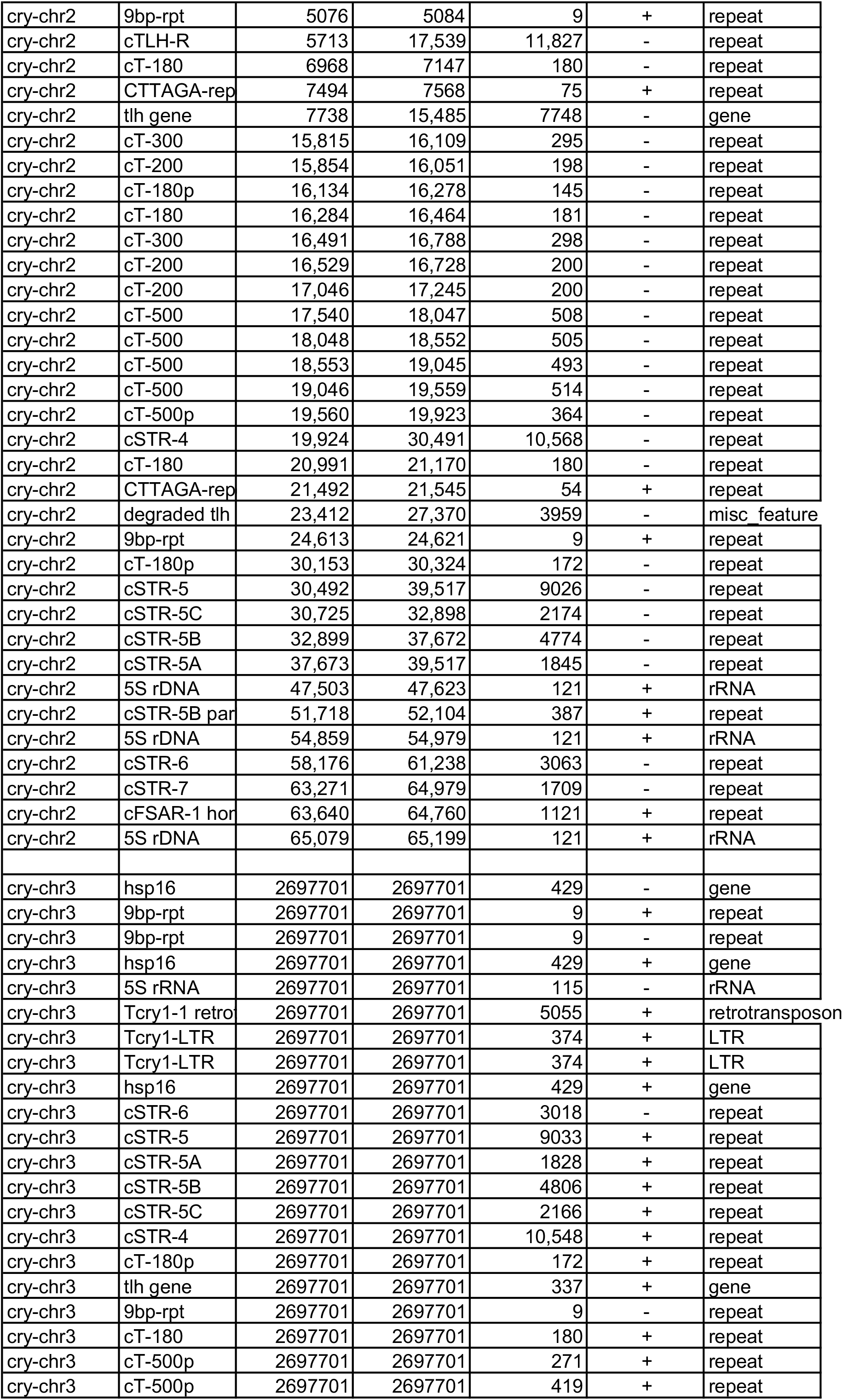
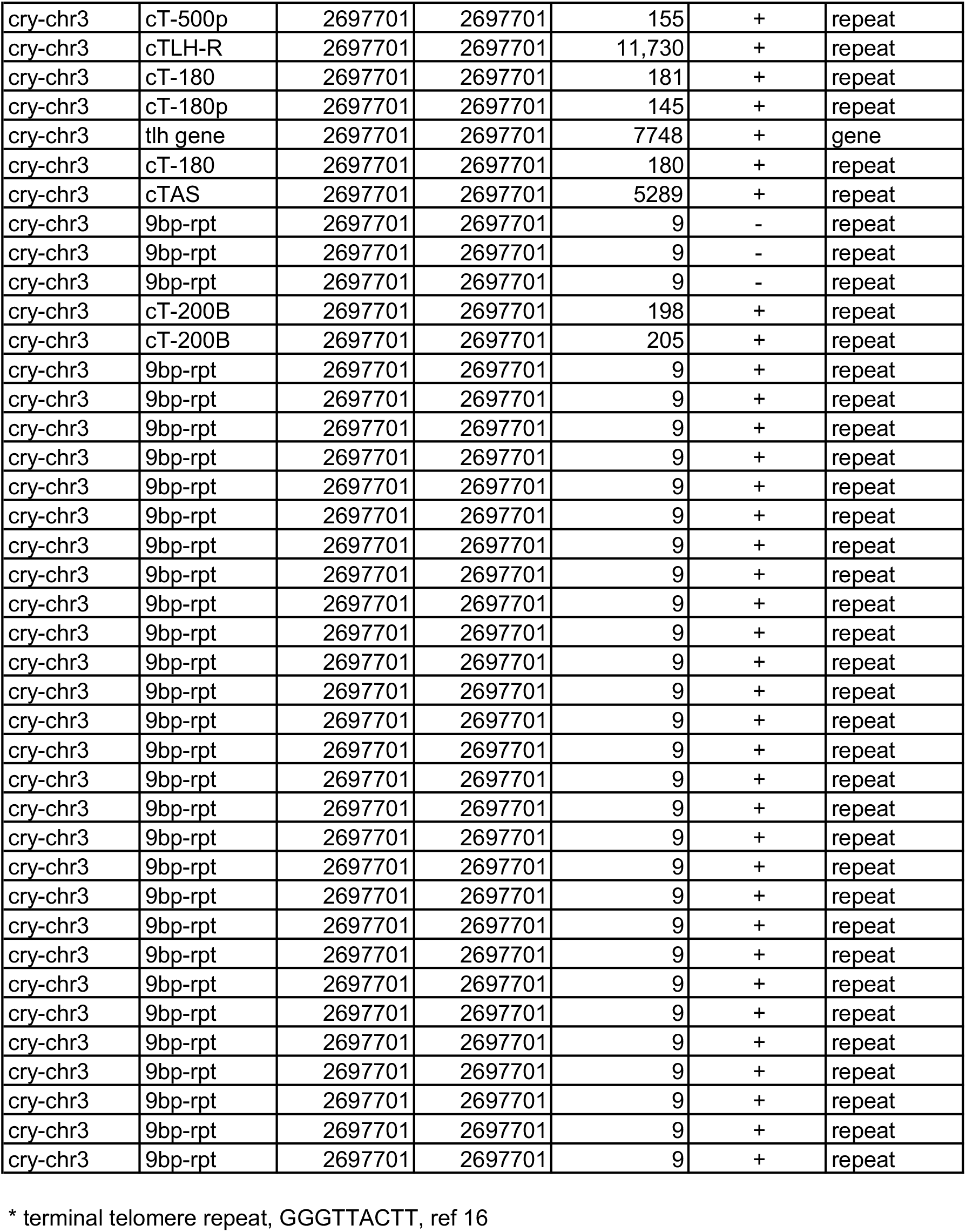
S. cryophilus telomere repeat annotation

**Supplementary Table 2:**
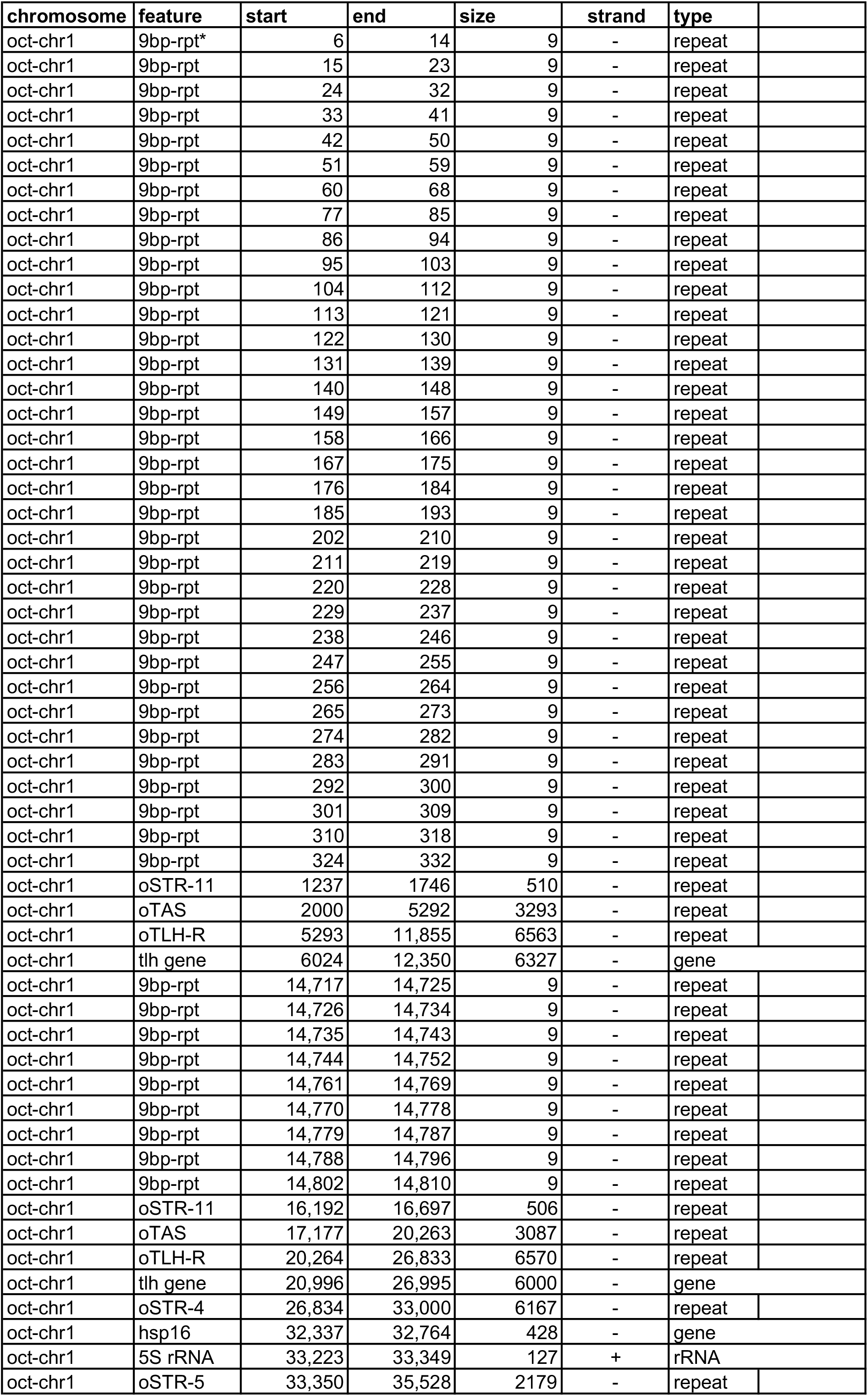
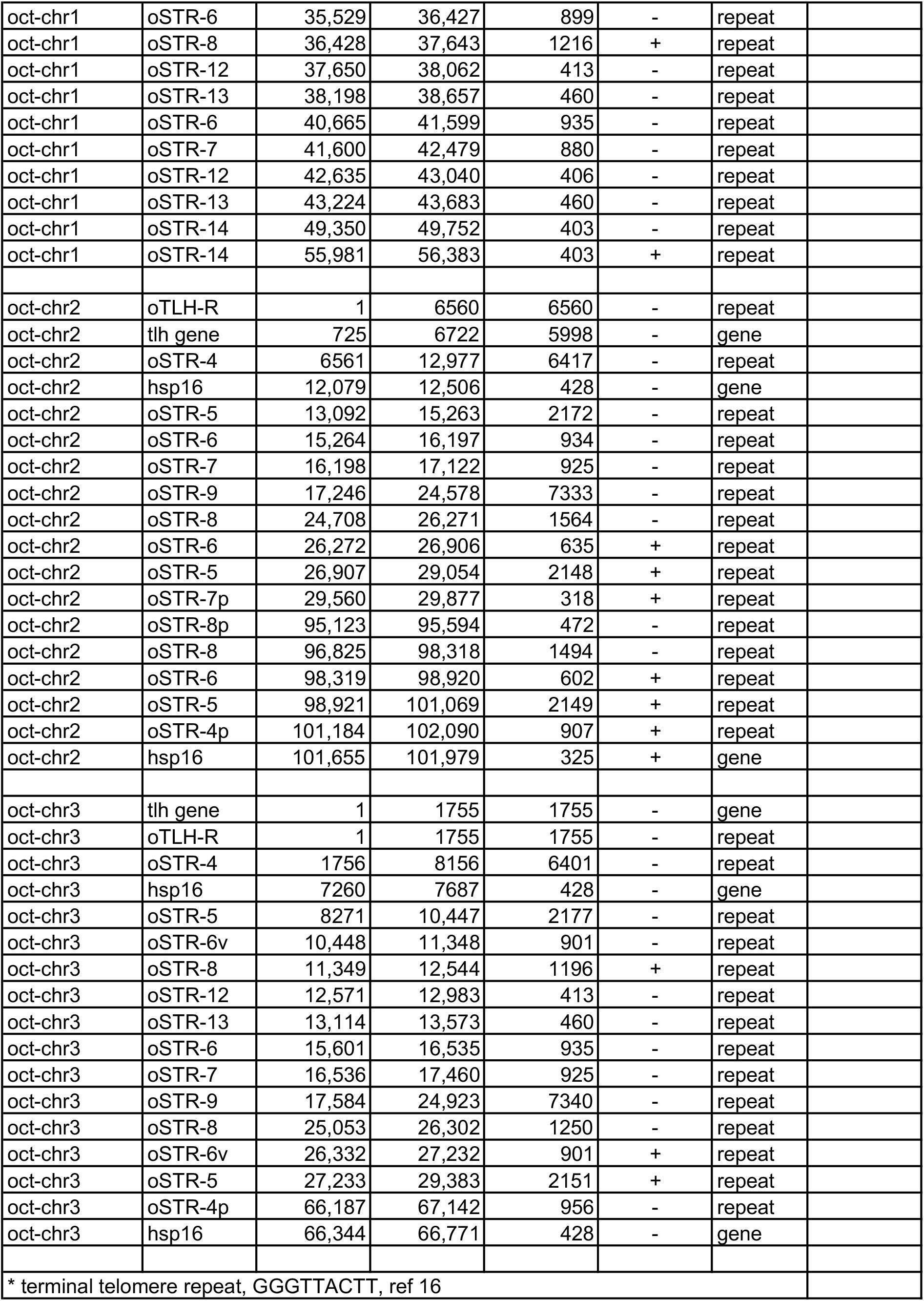
S. octosporus telomere repeat annotation

**Supplementary Table 3:** S. cryophilus centromere repeat annotation

| Repeats | Features | tRNA-anticodon | size (kb) | % GC | comments |
| --- | --- | --- | --- | --- | --- |
| cFSAR-1 | 5S rDNA; ORF |  | 3.7 | 35 | ORF - hypothetical protein |
| cFSAR-2 | 5S rDNA; hsp16 |  | 4.1 | 35 |  |
| cFSAR-3 | 5S rDNA; hsp16 |  | 1.7 | 34 |  |
| cTAR-4 | tDNA: DV | AspGTC, ValAAC | 0.5 | 38 | several variants |
| cTAR-5 | tDNA: AIR | AlaAGC, IleAAT, ArgACG | 1.4 | 33 |  |
| cTAR-6 | tDNA: R | ArgACG | 0.7 | 34 | as RKL usually |
| cTAR-7 | tDNA: EF | GluCTC, PheGAA | 2.3 | 34 |  |
| cTAR-8 | tDNA: KE | LysCTT, GluCTC | 0.7 | 34 | Part of imr3 |
| cTAR-9 | tDNA: LK | LeuCAA, LysCTT | 0.9 | 31 |  |
| cTAR-10 | tDNA: NM | AsnGTT, MetCAT, | 1.3 | 33 | always as NME (but E sometimes alone in cTAR-14) |
| cTAR-11 | tDNA: A | AlaAGC | 1.6 | 31 |  |
| cTAR-12 | tDNA: E | GluTTC | 2.5 | 31 |  |
| cTAR-13 | tDNA: L | LeuAAG | 1.8 | 31 |  |
| cTAR-14 | tDNA: E | GluTTC | 6.2 | 34 | retrotransposon remnant, sometimes alone, sometimes as NME |
| cHR-15 |  |  | 3.8 | 34 |  |
| cHR-16 |  |  | 1.4 | 35 |  |
| cHR-17 |  |  | 1.5 | 34 |  |
| cHR-18 |  |  | 0.7 | 31 |  |
| cHR-19 |  |  | 1.9 | 33 |  |
| cHR-20 |  |  | 1.4 | 33 |  |
| cHR-21 |  |  | 0.7 | 33 |  |
| cCNT-L |  |  | 6 | 33 |  |
| cCNT-S |  |  | 1.3 | 31 |  |
| c-cnt1 |  |  | 10.1 | 33 | Contains cCNT-L and cCNT-S |
| c-cnt2 |  |  | 10.5 | 32 | Contains cCNT-L and cCNT-S |
| c-cnt3 |  |  | 3.9 | 34 |  |
| c-imr1 | tDNA: LK | LeuCAA, LysCTT | 1.3 | 31 | Contains cTAR-9 |
| c-imr2 | tDNA: DVAIRKE | AspGTC, ValAAC, AlaAGC, IleAAT, ArgACG, LysCTT, GluCTC | 1.5 | 32 | Contains cTAR-4, cTAR-5, cTAR-8 |
| c-imr3 | tDNA: T; LTR | ThrAGT | 4.7 | 31 |  |

**Supplementary Table 5:**
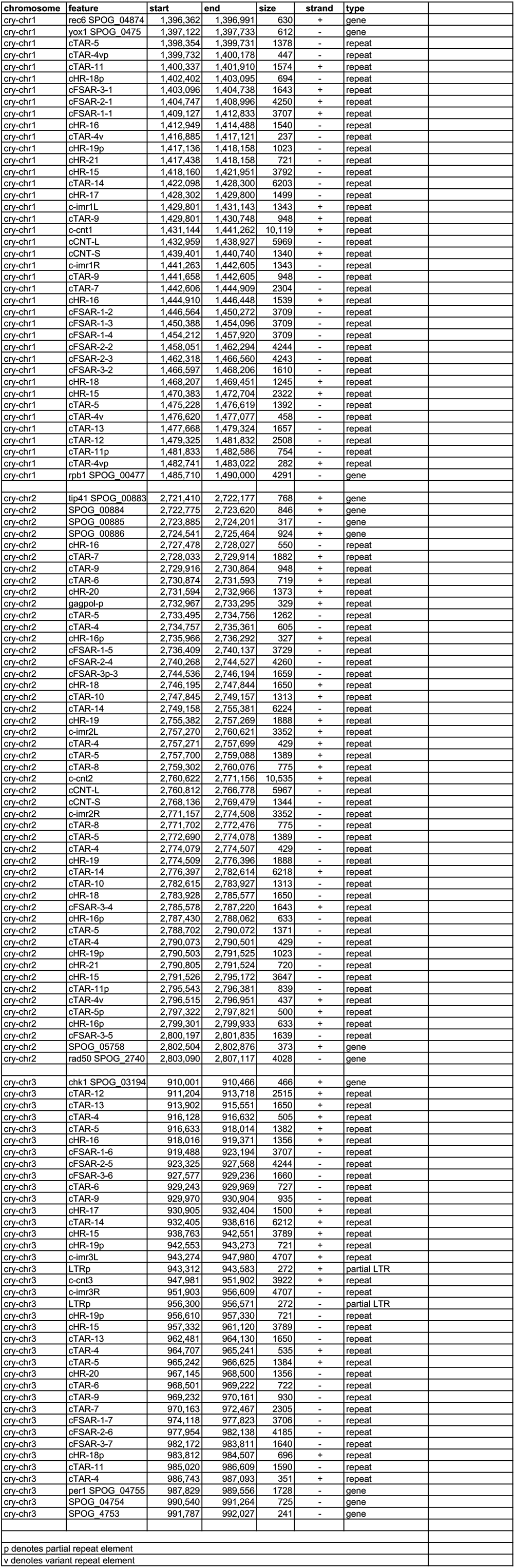
S. cryophilus centromere repeat cordinates

**Supplementary Table 6:**
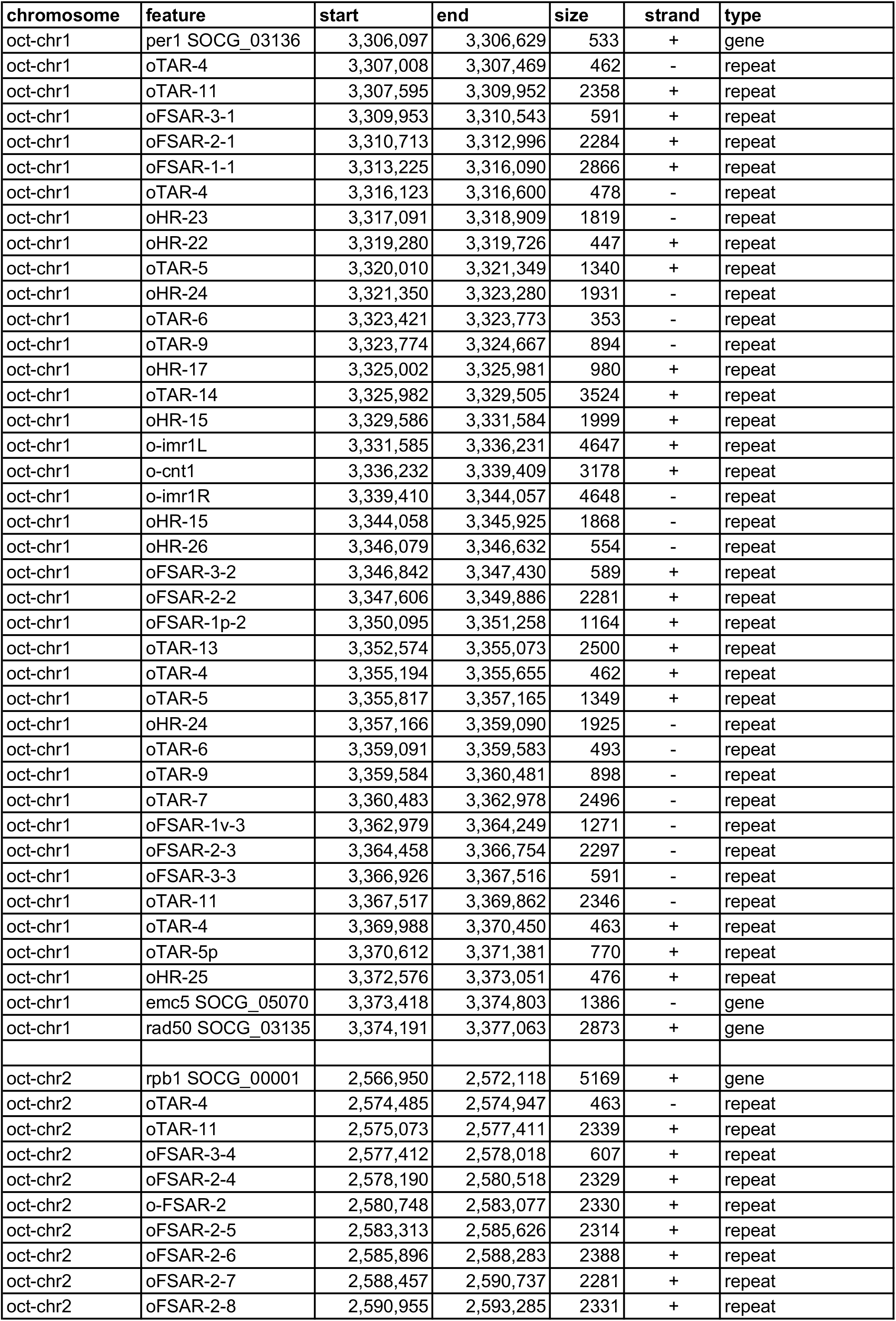
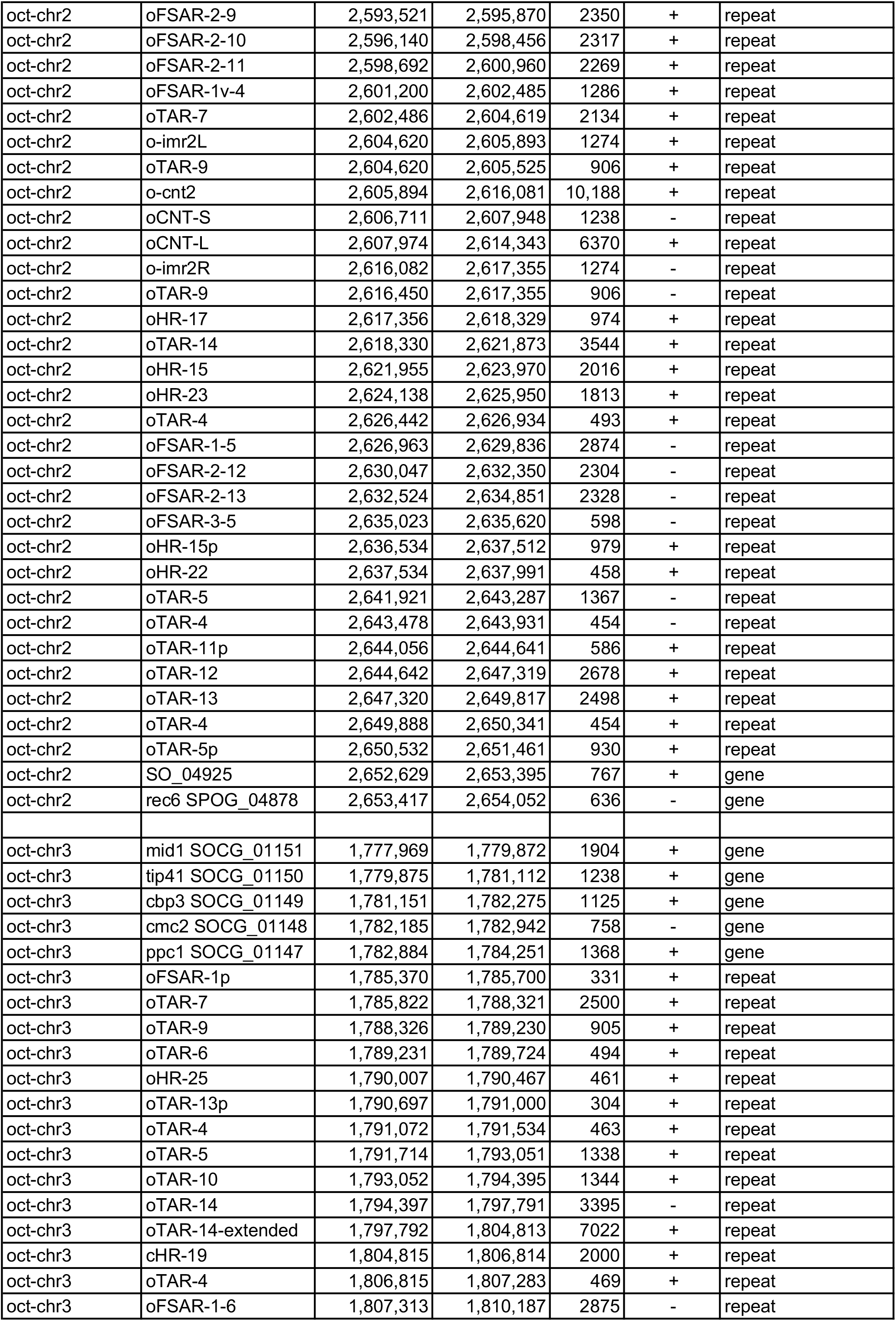
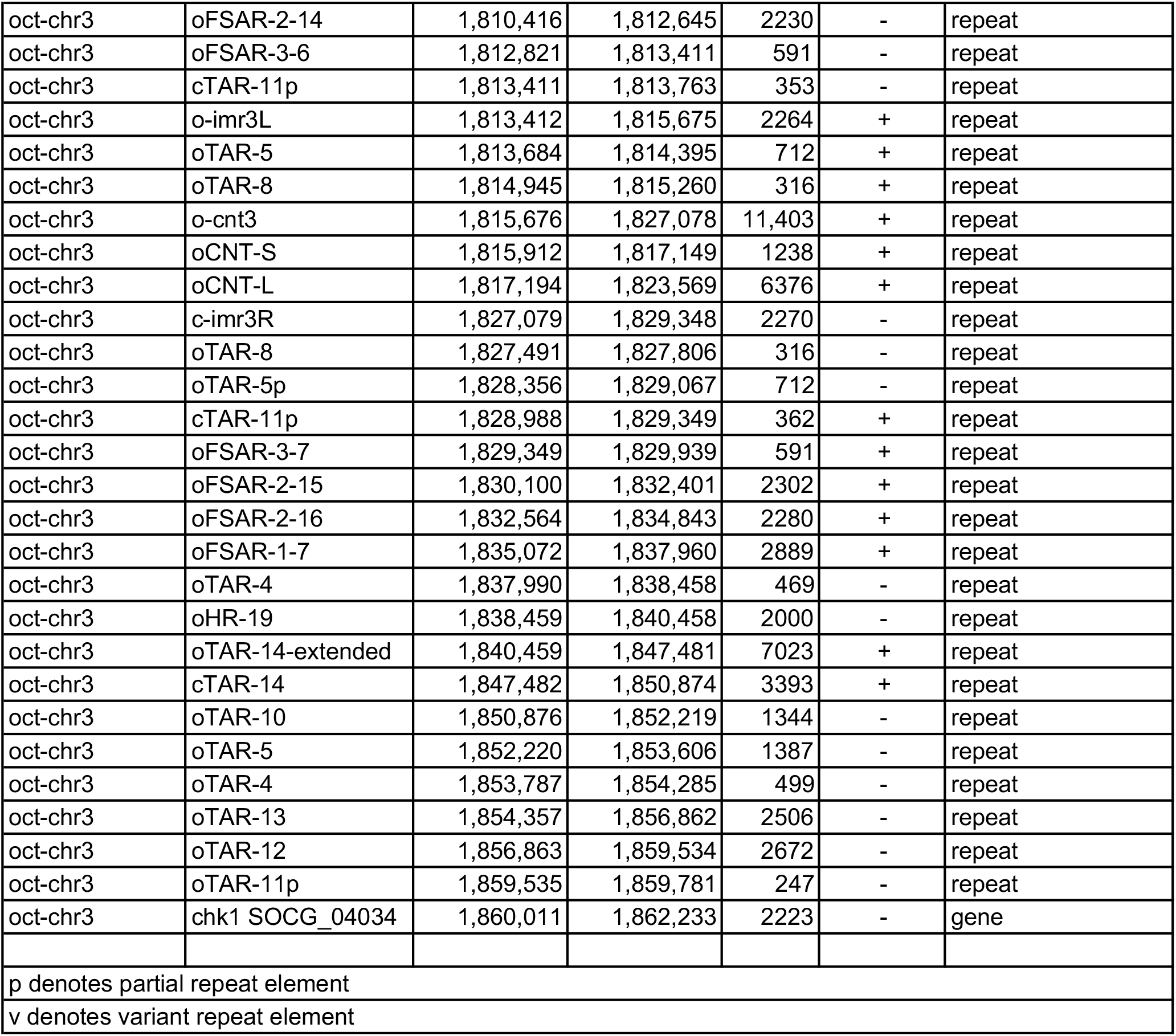
S. octosporus centromere repeat coordinates

**Supplementary Table 7:** S. pombe centromere repeat annotation

| **Feature** | **size (kb)** | **% GC** |
| --- | --- | --- |
| cc1 | 4.0 | 29.1 |
| cc2 | 6.6 | 28.9 |
| cc3 | 4.7 | 29.0 |
| cc1 & cc3 homology region<br>(TM element) | 3.2 | 28.0 |
| dg | 4.1-4.5 | 33-34 |
| dh | 4.1-6.7 | 32-34 |
| dh variant | 2.1 | 36.0 |
| dh partial | 0.3-0.4 | 29-37 |
| imr1 | 5.1 | 28.9 |
| imr2 | 4.1 | 29.1 |
| imr3 | 5.9 | 30.8 |
| imr1 partial | 0.7 | 28.9 |
| imr2 partial | 0.8-0.9 | 29.8 |
| imr3 partial | 1.1 | 33.3 |

**Supplementary Table 8:**
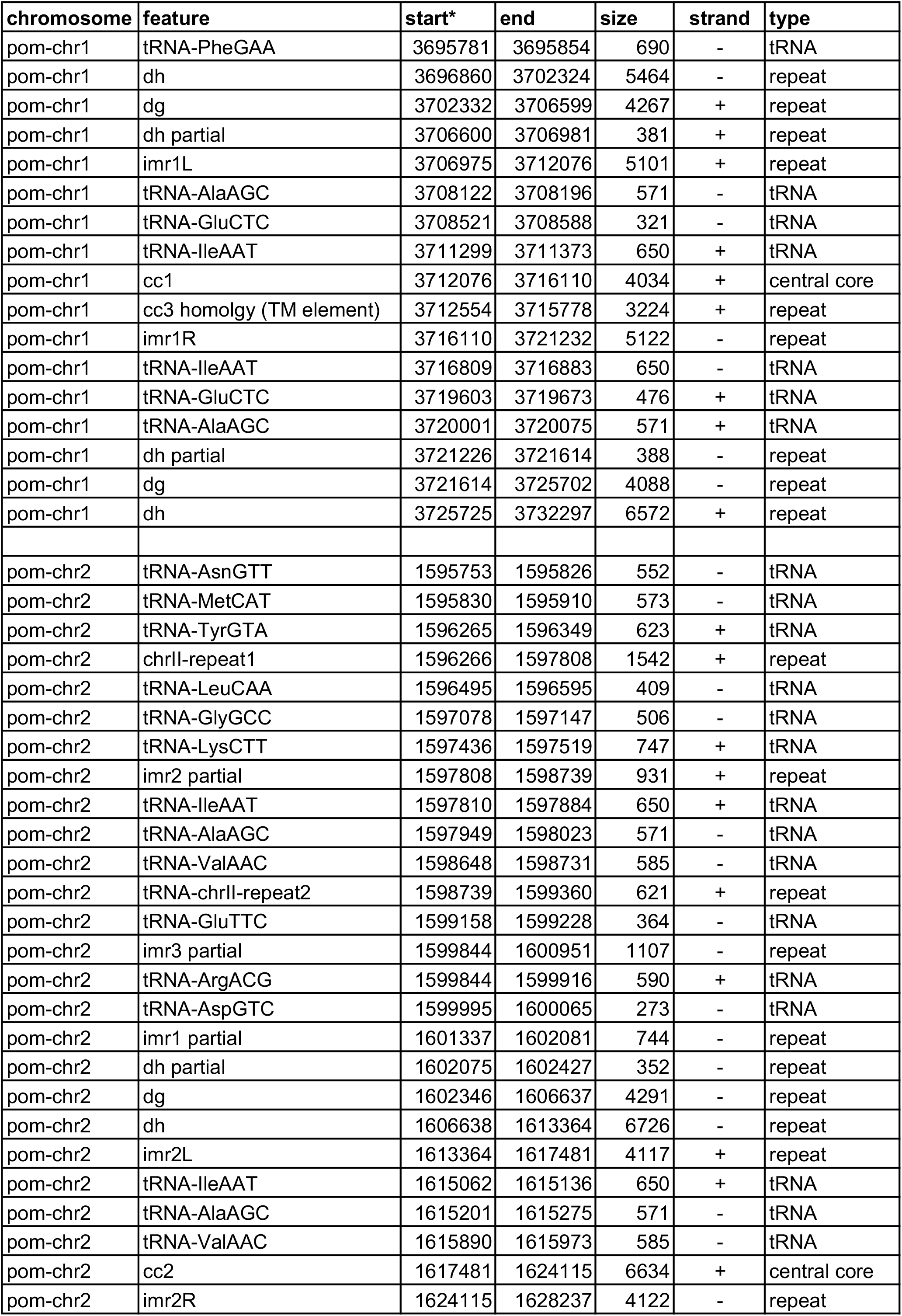
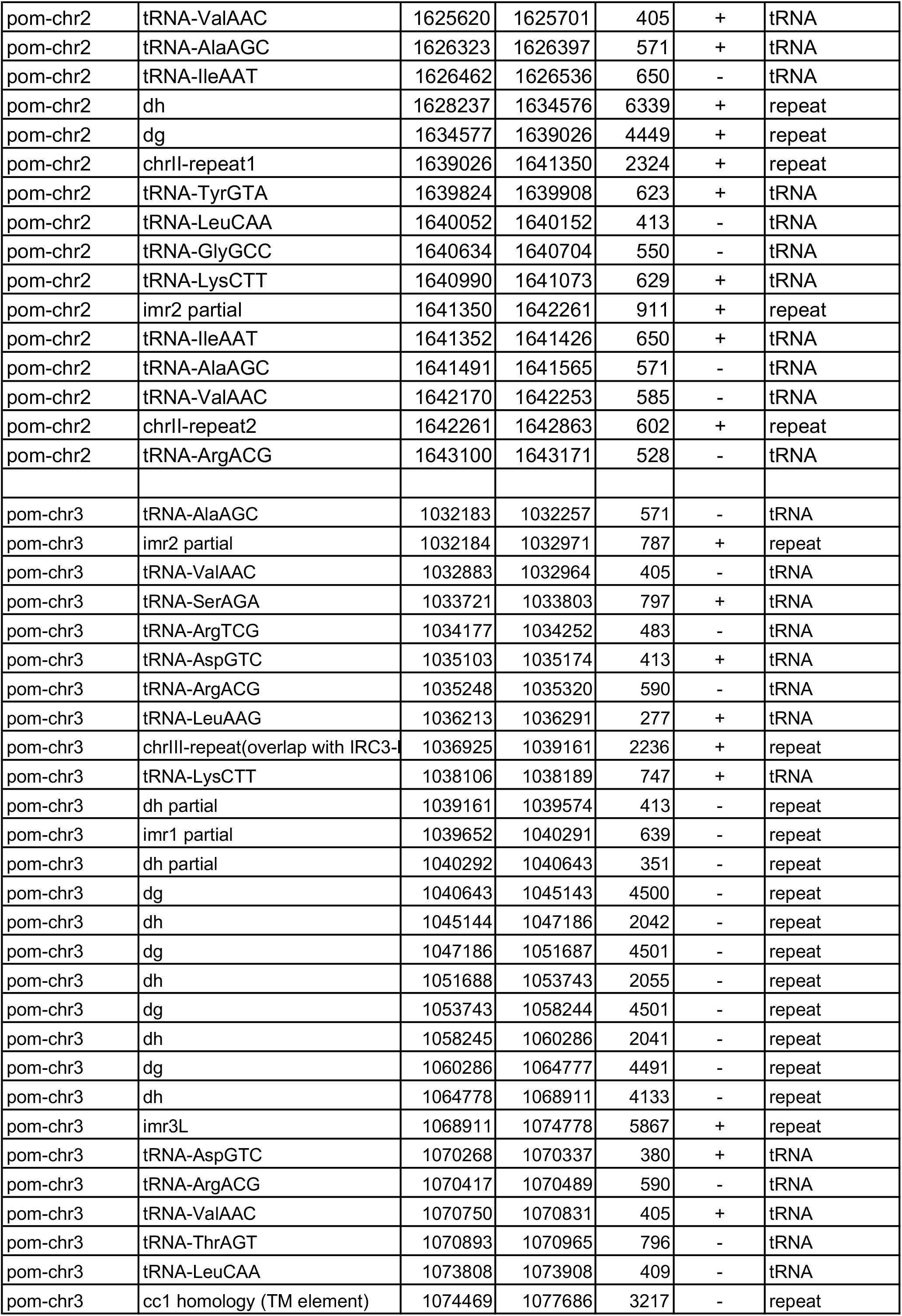
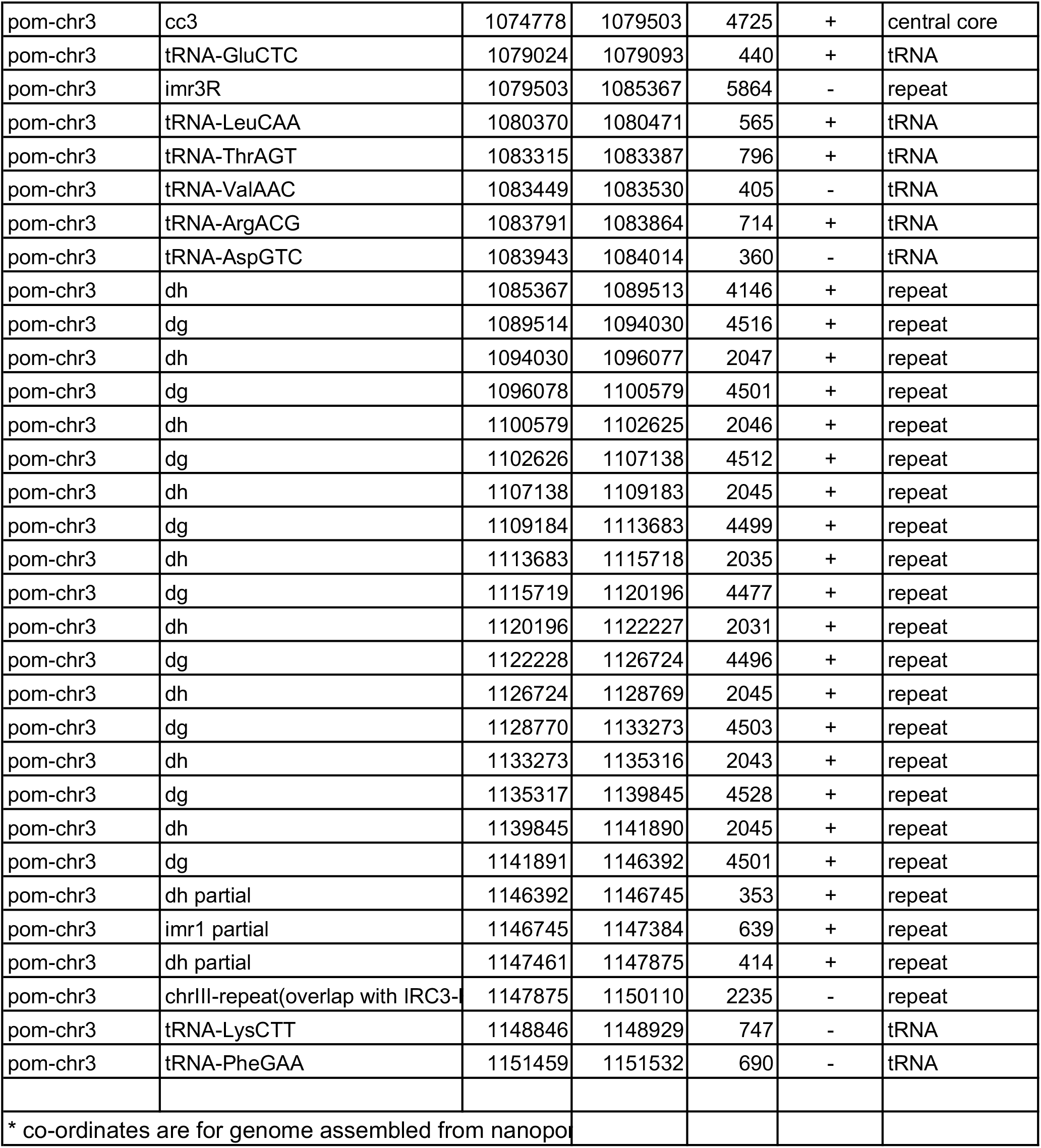
S. pombe centromere repeat coordinates

**Supplementary Table 9:** S. japonicus retrotransposons

| **Retrotransposon** | **Present at putative centromeres?** | **Heterochromatin** | **CENP-A** |
| --- | --- | --- | --- |
| Tj1 | rare | low | - |
| Tj2 | common | High | - |
| Tj3* | common | High | - |
| Tj4 | possibly telomere specific | Intermediate | - |
| Tj5* | common | High | - |
| Tj6 | common | Low-intermediate | Intermediate |
| Tj7 | common | Low-intermediate | High |
|  |  |  | Partials have low levels |
| Tj8* | common | High | - |
| Tj9* | common | High | - |
| Tj10 | common | High | - |
| Tj11** | common | High | - |
|  |  | Partials have low levels |  |

\*Tj3, Tj5, Tj8 and Tj9 have high homology.
\*\*Newly defined putative retrotransposon. 5501 bp. Present in *Schizosaccharomyces japonicus*.GCA\_000149845.2 supercontig 5.5: 40010-45479 (Rhind, 2011)

**Supplementary Table 10:**
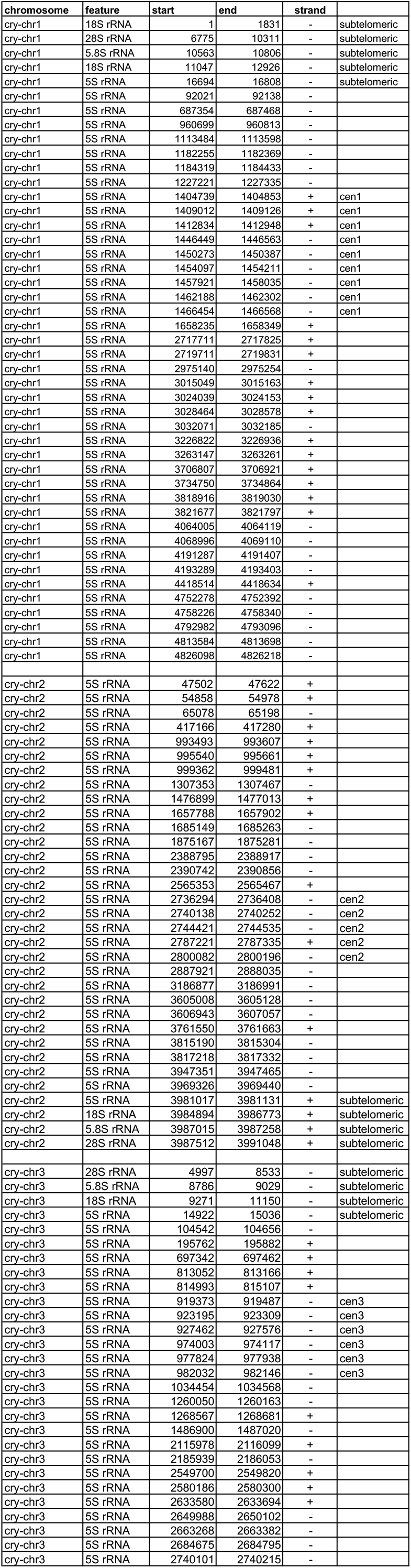
S. cryophilus rDNA annotation

**Supplementary Table 11:** S. octosporus rDNA annotation

| chromosome | feature | start | end | strand |
| --- | --- | --- | --- | --- |
| oct-chr1 | 5S rRNA | 33235 | 33349 | + |
| oct-chr1 | 5S rRNA | 413920 | 414039 | + |
| oct-chr1 | 5S rRNA | 880727 | 880841 | + |
| oct-chr1 | 5S rRNA | 1078204 | 1078318 | - |
| oct-chr1 | 5S rRNA | 1080784 | 1080894 | - |
| oct-chr1 | 5S rRNA | 1264920 | 1265034 | - |
| oct-chr1 | 5S rRNA | 1270014 | 1270128 | - |
| oct-chr1 | 5S rRNA | 1276840 | 1276954 | - |
| oct-chr1 | 5S rRNA | 1787532 | 1787646 | - |
| oct-chr1 | 5S rRNA | 1794347 | 1794461 | - |
| oct-chr1 | 5S rRNA | 2031940 | 2032054 | + |
| oct-chr1 | 5S rRNA | 2829548 | 2829662 | - |
| oct-chr1 | 5S rRNA | 2964472 | 2964586 | - |
| oct-chr1 | 5S rRNA | 3029426 | 3029540 | - |
| oct-chr1 | 5S rRNA | 3258531 | 3258645 | + |
| oct-chr1 | 5S rRNA | 3310547 | 3310661 | + |
| oct-chr1 | 5S rRNA | 3313022 | 3313136 | + |
| oct-chr1 | 5S rRNA | 3347434 | 3347548 | + |
| oct-chr1 | 5S rRNA | 3349892 | 3350006 | + |
| oct-chr1 | 5S rRNA | 3364338 | 3364452 | - |
| oct-chr1 | 5S rRNA | 3366808 | 3366922 | - |
| oct-chr1 | 5S rRNA | 3373028 | 3373142 | + |
| oct-chr1 | 5S rRNA | 3456387 | 3456501 | - |
| oct-chr1 | 5S rRNA | 3753939 | 3754086 | - |
| oct-chr1 | 5S rRNA | 3756499 | 3756613 | - |
| oct-chr1 | 5S rRNA | 4164520 | 4164643 | - |
| oct-chr1 | 5S rRNA | 4167108 | 4167222 | - |
| oct-chr1 | 5S rRNA | 4361472 | 4361586 | + |
| oct-chr1 | 18S rRNA | 4514514 | 4516392 | + |
| oct-chr1 | 5.8S rRNA | 4516668 | 4516914 | + |
| oct-chr1 | 28S rRNA | 4517197 | 4519984 | + |
| oct-chr2 | 5S rRNA | 12977 | 13091 | + |
| oct-chr2 | 5S rRNA | 29055 | 29169 | - |
| oct-chr2 | 5S rRNA | 96705 | 96824 | - |
| oct-chr2 | 5S rRNA | 101070 | 101184 | - |
| oct-chr2 | 5S rRNA | 393309 | 393428 | + |
| oct-chr2 | 5S rRNA | 672545 | 672659 | - |
| oct-chr2 | 5S rRNA | 1041835 | 1041949 | + |
| oct-chr2 | 5S rRNA | 1114943 | 1115057 | - |
| oct-chr2 | 5S rRNA | 1571359 | 1571473 | + |
| oct-chr2 | 5S rRNA | 1739423 | 1739537 | + |
| oct-chr2 | 5S rRNA | 1957207 | 1957323 | - |
| oct-chr2 | 5S rRNA | 1999314 | 1999433 | + |
| oct-chr2 | 5S rRNA | 2249382 | 2249496 | - |
| oct-chr2 | 5S rRNA | 2400626 | 2400740 | - |
| oct-chr2 | 5S rRNA | 2578022 | 2578136 | + |
| oct-chr2 | 5S rRNA | 2580580 | 2580694 | + |
| oct-chr2 | 5S rRNA | 2583145 | 2583259 | + |
| oct-chr2 | 5S rRNA | 2585728 | 2585842 | + |
| oct-chr2 | 5S rRNA | 2588289 | 2588403 | + |
| oct-chr2 | 5S rRNA | 2590787 | 2590901 | + |
| oct-chr2 | 5S rRNA | 2593353 | 2593467 | + |
| oct-chr2 | 5S rRNA | 2595972 | 2596086 | + |
| oct-chr2 | 5S rRNA | 2598524 | 2598638 | + |
| oct-chr2 | 5S rRNA | 2601010 | 2601124 | + |
| oct-chr2 | 5S rRNA | 2629927 | 2630041 | - |
| oct-chr2 | 5S rRNA | 2632404 | 2632518 | - |
| oct-chr2 | 5S rRNA | 2634905 | 2635019 | - |
| oct-chr2 | 5S rRNA | 2896020 | 2896134 | - |
| oct-chr2 | 5S rRNA | 2963574 | 2963688 | - |
| oct-chr2 | 5S rRNA | 3120708 | 3120822 | - |
| oct-chr2 | 5S rRNA | 3607716 | 3607830 | - |
| oct-chr2 | 5S rRNA | 3610298 | 3610412 | - |
| oct-chr2 | 5S rRNA | 3618746 | 3618860 | - |
| oct-chr2 | 18S rRNA | 3789499 | 3791378 | + |
| oct-chr2 | 5.8S rRNA | 3791653 | 3791899 | + |
| oct-chr2 | 28S rRNA | 3792181 | 3794048 | + |
| oct-chr3 | 5S rRNA | 8156 | 8270 | + |
| oct-chr3 | 5S rRNA | 29384 | 29500 | - |
| oct-chr3 | 5S rRNA | 104712 | 104826 | - |
| oct-chr3 | 5S rRNA | 132256 | 132372 | - |
| oct-chr3 | 5S rRNA | 140190 | 140304 | - |
| oct-chr3 | 5S rRNA | 161992 | 162106 | - |
| oct-chr3 | 5S rRNA | 642970 | 643084 | + |
| oct-chr3 | 5S rRNA | 766517 | 766631 | + |
| oct-chr3 | 5S rRNA | 772371 | 772485 | + |
| oct-chr3 | 5S rRNA | 774887 | 775006 | + |
| oct-chr3 | 5S rRNA | 1005175 | 1005289 | - |
| oct-chr3 | 5S rRNA | 1083257 | 1083371 | - |
| oct-chr3 | 5S rRNA | 1107657 | 1107771 | - |
| oct-chr3 | 5S rRNA | 1114356 | 1114470 | - |
| oct-chr3 | 5S rRNA | 1560721 | 1560835 | - |
| oct-chr3 | 5S rRNA | 1600041 | 1600155 | - |
| oct-chr3 | 5S rRNA | 1623740 | 1623854 | + |
| oct-chr3 | 5S rRNA | 1810276 | 1810390 | - |
| oct-chr3 | 5S rRNA | 1812703 | 1812817 | - |
| oct-chr3 | 5S rRNA | 1829943 | 1830057 | + |
| oct-chr3 | 5S rRNA | 1832407 | 1832521 | + |
| oct-chr3 | 5S rRNA | 1834869 | 1834983 | + |
| oct-chr3 | 5S rRNA | 1952114 | 1952228 | - |
| oct-chr3 | 5S rRNA | 2029273 | 2029387 | + |
| oct-chr3 | 5S rRNA | 2894060 | 2894174 | + |
| oct-chr3 | 5S rRNA | 2952779 | 2952893 | + |
| oct-chr3 | 5S rRNA | 2955384 | 2955508 | + |
| oct-chr3 | 18S rRNA | 2966653 | 2968532 | + |
| oct-chr3 | 5.8S rRNA | 2968807 | 2969053 | - |
| oct-chr3 | 28S rRNA | 2969336 | 2971974 | + |

**Supplementary Table 12:** Homology between FSAR repeats within S. cryophilus and S. octosporus

| **S.cry FSAR identity %** |  |  |  |  |  |  |  |
| --- | --- | --- | --- | --- | --- | --- | --- |
|  | cFSAR-1-1 |  |  | cFSAR-2-1 |  |  | cFSAR-3-1 |
| cFSAR-1-1 | 100 |  | cFSAR-2-1 | 100 |  | cFSAR-3-1 | 100 |
| cFSAR-1-2 | 99.95 |  | cFSAR-2-2 | 99.67 |  | cFSAR-3-2 | 98.7 |
| cFSAR-1-3 | 99.95 |  | cFSAR-2-3 | 99.6 |  | cFSAR-3-3 | 93.95 |
| cFSAR-1-4 | 99.95 |  | cFSAR-2-4 | 99.44 |  | cFSAR-3-4 | 95.13 |
| cFSAR-1-7 | 97.57 |  | cFSAR-2-5 | 96.83 |  | cFSAR-3-5 | 99.51 |
| cFSAR-1-5 | 95.82 |  | cFSAR-2-6 | 98.78 |  | cFSAR-3-6 | 95.26 |
| cFSAR-1-6 | 95.66 |  |  |  |  | cFSAR-3p-7 | 94 |
| **S.oct FSAR identity %** |  |  |  |  |  |  |  |
|  | oFSAR-1-1 |  |  | oFSAR-2-1 |  |  | oFSAR-3-1 |
| oFSAR-1-1 | 100 |  | oFSAR-2-1 | 100 |  | oFSAR-3-1 | 100 |
| oFSAR-1-2 | 99.07 |  | oFSAR-2-2 | 97.47 |  | oFSAR-3-2 | 99.83 |
| oFSAR-1-3 | 98.12 |  | oFSAR-2-3 | 97.05 |  | oFSAR-3-3 | 98.98 |
| oFSAR-1-4 | 98.37 |  | oFSAR-2-4 | 97 |  | oFSAR-3-4 | 98.98 |
| oFSAR-1p-5 | 97.25 |  | oFSAR-2-5 | 98 |  | oFSAR-3-5 | 98.98 |
| oFSAR-1v-6 | 96.57 |  | oFSAR-2-6 | 97.78 |  | oFSAR-3-6 | 96.38 |
| oFSAR-1v-7 | 95.75 |  | oFSAR-2-7 | 95.38 |  | oFSAR-3-7 | 97.66 |
|  |  |  | oFSAR-2-8 | 96.16 |  |  |  |
|  |  |  | oFSAR-2-9 | 95.78 |  |  |  |
|  |  |  | oFSAR-2-10 | 97.52 |  |  |  |
|  |  |  | oFSAR-2-11 | 95.4 |  |  |  |
|  |  |  | oFSAR-2-12 | 98.2 |  |  |  |
|  |  |  | oFSAR-2-12 | 96.21 |  |  |  |
|  |  |  | oFSAR-2-13 | 96.16 |  |  |  |
|  |  |  | oFSAR-2-14 | 97.99 |  |  |  |
|  |  |  | oFSAR-2-15 | 97.73 |  |  |  |
|  |  |  | oFSAR-2-16 | 96.75 |  |  |  |

**Supplementary Table 13:** S. cryophilus and S. octosporus Hsp16 gene annotation and coordinates

| **chromosome** | **hsp16 ORF** | **start** | **end** | **strand** |
| --- | --- | --- | --- | --- |
| cry-chr1 | cry-ORF1-cFSAR-3-1 | 1403407 | 1403798 | - |
| cry-chr1 | cry-ORF2-cFSAR-2-1 | 1405538 | 1405959 | - |
| cry-chr1 | cry-ORF3-cFSAR-2-2 | 1461082 | 1461503 | + |
| cry-chr1 | cry-ORF4-cFSAR-2-3 | 1465348 | 1465769 | + |
| cry-chr1 | cry-ORF5-cFSAR-3-2 | 1467506 | 1467895 | + |
| cry-chr1 | cry-ORF6 | 2716245 | 2716666 | - |
| cry-chr1 | cry-ORF7 | 3738213 | 3738633 | - |
| cry-chr1 | cry-ORF8 | 4065105 | 4065526 | + |
| cry-chr1 | cry-ORF9 | 4787816 | 4788541 | - |
| cry-chr1 | cry-ORF10 | 4819528 | 4819949 | + |
| cry-chr1 | cry-ORF11 | 4876023 | 4876444 | + |
| cry-chr2 | cry-ORF12-cFSAR-2-4 | 2743315 | 2743736 | + |
| cry-chr2 | cry-ORF13-cFSAR-3p-3 | 2745491 | 2745883 | + |
| cry-chr2 | cry-ORF14-cFSAR-3-4 | 2785889 | 2786281 | - |
| cry-chr2 | cry-ORF15-cFSAR-3-5 | 2801168 | 2801560 | + |
| cry-chr3 | cry-ORF16-cFSAR-2-5 | 926356 | 926777 | + |
| cry-chr3 | cry-ORF17-cFSAR-3-6 | 928536 | 928925 | + |
| cry-chr3 | cry-ORF18-cFSAR-2-6 | 980964 | 981385 | + |
| cry-chr3 | cry-ORF19-cFSAR-3-7 | 983108 | 983500 | + |
| cry-chr3 | cry-ORF20 | 2539982 | 2540707 | - |
| cry-chr3 | cry-ORF21 | 2555876 | 2556297 | - |
| cry-chr3 | cry-ORF22 | 2706305 | 2706726 | - |
| cry-chr3 | cry-ORF23 | 2734174 | 2734595 | + |
| cry-chr3 | cry-ORF24 | 2754385 | 2754806 | + |
| **chromosome** | **ORF** | **start** | **end** | **strand** |
| oct-chr1 | oct-ORF1 | 32330 | 32773 | - |
| oct-chr2 | oct-ORF2 | 12072 | 12515 | - |
| oct-chr2 | oct-ORF3 | 101646 | 102089 | + |
| oct-chr3 | oct-ORF4 | 7253 | 7696 | - |
| oct-chr3 | oct-ORF5 | 66337 | 66780 | - |
| oct-chr3 | oct-ORF6 | 140764 | 141207 | + |
| oct-chr3 | oct-ORF7 | 162568 | 163011 | + |
| oct-chr3 | oct-ORF8 | 2892999 | 2893652 | - |

**Supplementary Table 14:** S. cryophilus retrotansposon and LTR annotation and coordinates

| **chromosome** | **start** | **end** | **Element** | **strand** | **size** |
| --- | --- | --- | --- | --- | --- |
| cry-chr1 | 26090 | 28101 | Tcry1 partial retrotransposon | - | 2011 |
| cry-chr1 | 30927 | 31247 | Tcry1-type LTR | - | 320 |
| cry-chr1 | 30927 | 31288 | Tcry1-type LTR | - | 361 |
| cry-chr1 | 965190 | 965585 | Tcry1 partial retrotransposon | + | 395 |
| cry-chr1 | 965223 | 965585 | Tcry1-type LTR | + | 362 |
| cry-chr1 | 2024308 | 2024682 | Tcry1-type LTR | - | 374 |
| cry-chr1 | 2024308 | 2024714 | Tcry1-type LTR | - | 406 |
| cry-chr1 | 4844387 | 4844681 | Tcry1-type LTR | - | 294 |
| cry-chr1 | 4844387 | 4844714 | Tcry1-type LTR | - | 327 |
| cry-chr2 | 1431930 | 1434972 | Tcry1-type LTR | - | 3042 |
| cry-chr2 | 1434981 | 1436403 | Tcry1-type LTR | - | 1422 |
| cry-chr3 | 24898 | 25200 | Tcry1-type LTR | - | 302 |
| cry-chr3 | 24898 | 25238 | Tcry1-type LTR | - | 340 |
| cry-chr3 | 943270 | 943583 | Tcry1-type LTR | + | 313 |
| cry-chr3 | 943312 | 943583 | Tcry1-type LTR | + | 271 |
| cry-chr3 | 956300 | 956571 | Tcry1-type LTR | - | 271 |
| cry-chr3 | 956300 | 956613 | Tcry1-type LTR | - | 313 |
| cry-chr3 | 2743314 | 2748368 | Tcry1-1 | - | 5054 |

**Supplementary Table 15:** S. octosporus retrotransposon remnat annotation

| chromosome | homology with<br>locus | Tcry1 |  |  | transposon remnant S.oct-mat |  |  | oTAR-14ex |  |  |
| --- | --- | --- | --- | --- | --- | --- | --- | --- | --- | --- |
|  |  | start | end | size | start | end | size | start | end | size |
| S.oct-chr1 | mat locus | 834081 | 835897 | 1816 | 834236 | 836137 | 1901 | 833988 | 835179 | 1191 |
|  |  |  |  |  | 821484 | 821925 | 441 | 835889 | 836945 | 1056 |
| S.oct-chr2 | retrotransposon homology |  |  |  |  |  |  | 761733 | 762238 | 505 |
| S.oct-chr3 | retrotransposon homology | 134707 | 135239 | 532 | 134071 | 135804 | 1733 | 132948 | 134195 | 1247 |
|  |  |  |  |  |  |  |  | 134834 | 136047 | 1213 |
|  | centromere: oTAR-14ex | 1801780 | 1802233 | 453 | 1801805 | 1802703 | 898 | 1797792 | 1804814 | 7022 |
|  | centromere: oTAR-14ex | 1842409 | 1843493 | 1084 | 1842570 | 1843468 | 898 | 1840459 | 1847481 | 7022 |

**Supplementary Table 16:** S. pombe, S. octosporus and S. cruophilus tDNA frequencies within centromeres and genome wide

| species | % tDNAs at centromeres | mean frequency centromeres | mean frequency rest of genome | fold frequency centromeres vs genome |
| --- | --- | --- | --- | --- |
| *S. pombe* | 32 (55/171) | 3.8 kb | 105.6 kb | 27.8 |
| *S. octosporus* | 32 (96/298) | 2.3 kb | 54.8 kb | 23.6 |
| *S. cryophilus* | 32 (95/294) | 2.6 kb | 57.5 kb | 22.5 |

**Supplementary Table 17:** S. cryophilus tDNA coordinates

| Chromosome | tDNA-anticodon* | start | end | strand | cen located? |
| --- | --- | --- | --- | --- | --- |
| cry-chr1 | GlyGCC | 91817 | 91887 | - |  |
| cry-chr1 | PheGAA | 130564 | 130636 | + |  |
| cry-chr1 | PheGAA | 132151 | 132223 | + |  |
| cry-chr1 | ArgACG | 312388 | 312460 | + |  |
| cry-chr1 | GlyGCC | 505819 | 505889 | - |  |
| cry-chr1 | GlyGCC | 508234 | 508304 | + |  |
| cry-chr1 | GlyGCC | 509140 | 509210 | - |  |
| cry-chr1 | SerGCT | 535112 | 535206 | - |  |
| cry-chr1 | ThrAGT | 679941 | 680012 | + |  |
| cry-chr1 | GlyGCC | 687139 | 687209 | - |  |
| cry-chr1 | LysTTT | 690360 | 690434 | + |  |
| cry-chr1 | IleTAT | 731048 | 731146 | + |  |
| cry-chr1 | MetCAT | 916227 | 916298 | - |  |
| cry-chr1 | SerTGA | 916303 | 916399 | - |  |
| cry-chr1 | LysCTT | 949167 | 949249 | - |  |
| cry-chr1 | ProAGG | 949343 | 949414 | - |  |
| cry-chr1 | SerAGA | 959364 | 959445 | - |  |
| cry-chr1 | MetCAT | 983804 | 983884 | + |  |
| cry-chr1 | AsnGTT | 983891 | 983964 | + |  |
| cry-chr1 | LysCTT | 1000051 | 1000133 | + |  |
| cry-chr1 | ProAGG | 1000466 | 1000537 | + |  |
| cry-chr1 | HisGTG | 1034546 | 1034617 | + |  |
| cry-chr1 | ThrTGT | 1130756 | 1130827 | - |  |
| cry-chr1 | ArgTCG | 1161587 | 1161659 | + |  |
| cry-chr1 | LysTTT | 1222481 | 1222555 | + |  |
| cry-chr1 | GlnTTG | 1226243 | 1226314 | - |  |
| cry-chr1 | HisGTG | 1245191 | 1245262 | - |  |
| cry-chr1 | HisGTG | 1246203 | 1246274 | + |  |
| cry-chr1 | ArgACG | 1398356 | 1398428 | - | cen1 |
| cry-chr1 | IleAAT | 1398802 | 1398875 | + | cen1 |
| cry-chr1 | AlaAGC | 1399216 | 1399289 | - | cen1 |
| cry-chr1 | ValAAC | 1399733 | 1399815 | + | cen1 |
| cry-chr1 | AspGTC | 1400121 | 1400191 | - | cen1 |
| cry-chr1 | AlaAGC | 1401996 | 1402069 | - | cen1 |
| cry-chr1 | ValAAC | 1416701 | 1416783 | + | cen1 |
| cry-chr1 | AspGTC | 1417064 | 1417134 | - | cen1 |
| cry-chr1 | GluTTC | 1422294 | 1422365 | - | cen1 |
| cry-chr1 | LeuCAA | 1429970 | 1430074 | - | cen1 |
| cry-chr1 | LysCTT | 1430651 | 1430736 | + | cen1 |
| cry-chr1 | LysCTT | 1441670 | 1441755 | - | cen1 |
| cry-chr1 | LeuCAA | 1442332 | 1442436 | + | cen1 |
| cry-chr1 | GluCTC | 1442934 | 1443005 | + | cen1 |
| cry-chr1 | PheGAA | 1443462 | 1443534 | - | cen1 |
| cry-chr1 | ArgTCG | 1472926 | 1472998 | + | cen1 |
| cry-chr1 | ProAGG | 1474860 | 1474931 | - | cen1 |
| cry-chr1 | ArgACG | 1475230 | 1475302 | - | cen1 |
| cry-chr1 | IleAAT | 1475694 | 1475767 | + | cen1 |
| cry-chr1 | AlaAGC | 1476104 | 1476177 | - | cen1 |
| cry-chr1 | ValAAC | 1476621 | 1476703 | + | cen1 |
| cry-chr1 | AspGTC | 1477020 | 1477090 | - | cen1 |
| cry-chr1 | LeuAAG | 1478388 | 1478466 | - | cen1 |
| cry-chr1 | GluTTC | 1479575 | 1479646 | - | cen1 |
| cry-chr1 | AspGTC | 1482589 | 1482659 | + | cen1 |
| cry-chr1 | ValAAC | 1482934 | 1483016 | - | cen1 |
| cry-chr1 | AlaAGC | 1483401 | 1483474 | + | cen1 |
| cry-chr1 | ThrTGT | 1566588 | 1566659 | + |  |
| cry-chr1 | TrpCCA | 1600889 | 1600961 | + |  |
| cry-chr1 | LeuTAA | 1662939 | 1663038 | + |  |
| cry-chr1 | AlaAGC | 1699268 | 1699341 | + |  |
| cry-chr1 | IleAAT | 1770660 | 1770733 | + |  |
| cry-chr1 | ArgCCT | 1879809 | 1879913 | + |  |
| cry-chr1 | PheGAA | 1925186 | 1925258 | + |  |
| cry-chr1 | HisGTG | 2071565 | 2071636 | - |  |
| cry-chr1 | LeuCAA | 2193818 | 2193922 | - |  |
| cry-chr1 | GlnTTG | 2275238 | 2275309 | - |  |
| cry-chr1 | ValTAC | 2328129 | 2328201 | - |  |
| cry-chr1 | AsnGTT | 2475236 | 2475309 | - |  |
| cry-chr1 | GlyGCC | 2475535 | 2475605 | - |  |
| cry-chr1 | SerAGA | 2477035 | 2477116 | + |  |
| cry-chr1 | GluTTC | 2542169 | 2542240 | + |  |
| cry-chr1 | ProAGG | 2970237 | 2970308 | - |  |
| cry-chr1 | ProAGG | 2970837 | 2970908 | + |  |
| cry-chr1 | SerAGA | 3014773 | 3014854 | - |  |
| cry-chr1 | GlyGCC | 3028768 | 3028838 | + |  |
| cry-chr1 | SerAGA | 3226546 | 3226627 | - |  |
| cry-chr1 | TyrGTA | 3562905 | 3562988 | + |  |
| cry-chr1 | SerCGA | 3586114 | 3586210 | + |  |
| cry-chr1 | MetCAT | 3586221 | 3586292 | + |  |
| cry-chr1 | ValAAC | 3630054 | 3630135 | + |  |
| cry-chr1 | LeuAAG | 3659732 | 3659810 | - |  |
| cry-chr1 | SerGCT | 3661564 | 3661658 | + |  |
| cry-chr1 | AsnGTT | 3780368 | 3780441 | + |  |
| cry-chr1 | GluCTC | 3851396 | 3851467 | + |  |
| cry-chr1 | LysCTT | 3958742 | 3958824 | - |  |
| cry-chr1 | LysTTT | 3973358 | 3973432 | - |  |
| cry-chr1 | TyrGTA | 3984988 | 3985071 | + |  |
| cry-chr1 | IleAAT | 4010738 | 4010811 | + |  |
| cry-chr1 | GlnTTG | 4060210 | 4060281 | - |  |
| cry-chr1 | ThrCGT | 4108849 | 4108920 | - |  |
| cry-chr1 | LysCTT | 4109634 | 4109716 | - |  |
| cry-chr1 | AlaTGC | 4115466 | 4115537 | - |  |
| cry-chr1 | HisGTG | 4193751 | 4193822 | - |  |
| cry-chr1 | SerAGA | 4227149 | 4227230 | + |  |
| cry-chr1 | GlnTTG | 4383616 | 4383687 | + |  |
| cry-chr1 | ValCAC | 4383894 | 4383965 | + |  |
| cry-chr1 | ThrAGT | 4394451 | 4394522 | - |  |
| cry-chr1 | LeuTAA | 4458580 | 4458679 | + |  |
| cry-chr1 | GlnTTG | 4669217 | 4669288 | - |  |
| cry-chr1 | LysCTT | 4669566 | 4669648 | + |  |
| cry-chr1 | ArgTCT | 4690337 | 4690409 | + |  |
| cry-chr1 | GlyTCC | 4690555 | 4690625 | - |  |
| cry-chr1 | MetCAT | 4747095 | 4747166 | + |  |
| cry-chr1 | LeuAAG | 4747410 | 4747488 | - |  |
| cry-chr1 | ArgTCT | 4762521 | 4762593 | + |  |
| cry-chr1 | ProAGG | 4802986 | 4803057 | + |  |
| cry-chr1 | GlyTCC | 4803434 | 4803504 | + |  |
| cry-chr1 | GlyTCC | 4864713 | 4864783 | + |  |
| cry-chr2 | AsnGTT | 105306 | 105379 | - |  |
| cry-chr2 | ProAGG | 105557 | 105628 | + |  |
| cry-chr2 | AspGTC | 215535 | 215605 | + |  |
| cry-chr2 | AsnGTT | 426497 | 426570 | + |  |
| cry-chr2 | CysGCA | 551460 | 551531 | - |  |
| cry-chr2 | AlaCGC | 563920 | 564002 | + |  |
| cry-chr2 | ProTGG | 617444 | 617515 | + |  |
| cry-chr2 | GlyGCC | 992813 | 992883 | + |  |
| cry-chr2 | GlyGCC | 999636 | 999706 | + |  |
| cry-chr2 | LysTTT | 1024900 | 1024974 | - |  |
| cry-chr2 | LeuTAA | 1062616 | 1062714 | - |  |
| cry-chr2 | MetCAT | 1075030 | 1075110 | + |  |
| cry-chr2 | TrpCCA | 1100377 | 1100449 | + |  |
| cry-chr2 | SerAGA | 1122870 | 1122951 | - |  |
| cry-chr2 | GluCTC | 1195689 | 1195760 | + |  |
| cry-chr2 | ThrAGT | 1206451 | 1206522 | + |  |
| cry-chr2 | AsnGTT | 1273506 | 1273579 | - |  |
| cry-chr2 | MetCAT | 1273586 | 1273668 | - |  |
| cry-chr2 | GlnCTG | 1327870 | 1327941 | + |  |
| cry-chr2 | LysCTT | 1351024 | 1351106 | + |  |
| cry-chr2 | GlnCTG | 1351701 | 1351772 | + |  |
| cry-chr2 | GlnTTG | 1352057 | 1352128 | + |  |
| cry-chr2 | GlnTTG | 1353859 | 1353930 | + |  |
| cry-chr2 | GlyGCC | 1398779 | 1398849 | + |  |
| cry-chr2 | ThrAGT | 1437157 | 1437228 | + |  |
| cry-chr2 | GlyCCC | 1486779 | 1486849 | - |  |
| cry-chr2 | LeuAAG | 1581580 | 1581658 | + |  |
| cry-chr2 | ValCAC | 1594163 | 1594234 | - |  |
| cry-chr2 | PheGAA | 1674504 | 1674576 | - |  |
| cry-chr2 | GlyGCC | 1683338 | 1683408 | + |  |
| cry-chr2 | GlyGCC | 1684016 | 1684086 | - |  |
| cry-chr2 | TyrGTA | 1793743 | 1793826 | - |  |
| cry-chr2 | LeuCAG | 1794580 | 1794673 | + |  |
| cry-chr2 | MetCAT | 1832550 | 1832630 | + |  |
| cry-chr2 | GlyTCC | 1858872 | 1858942 | + |  |
| cry-chr2 | ProAGG | 1862800 | 1862871 | + |  |
| cry-chr2 | GluTTC | 1893328 | 1893399 | - |  |
| cry-chr2 | CysGCA | 1893890 | 1893961 | - |  |
| cry-chr2 | TyrGTA | 1991231 | 1991314 | - |  |
| cry-chr2 | ProCGG | 1991852 | 1991951 | + |  |
| cry-chr2 | AlaTGC | 2105091 | 2105162 | + |  |
| cry-chr2 | SerAGA | 2155606 | 2155687 | - |  |
| cry-chr2 | MetCAT | 2190173 | 2190244 | - |  |
| cry-chr2 | SerCGA | 2190255 | 2190351 | - |  |
| cry-chr2 | AlaAGC | 2288920 | 2288993 | + |  |
| cry-chr2 | GluTTC | 2316793 | 2316864 | - |  |
| cry-chr2 | ArgACG | 2428841 | 2428913 | - |  |
| cry-chr2 | ProTGG | 2532325 | 2532396 | - |  |
| cry-chr2 | SerAGA | 2566313 | 2566394 | + |  |
| cry-chr2 | LysCTT | 2571329 | 2571411 | + |  |
| cry-chr2 | SerGCT | 2623190 | 2623284 | - |  |
| cry-chr2 | CysGCA | 2637527 | 2637598 | + |  |
| cry-chr2 | ValTAC | 2667854 | 2667935 | + |  |
| cry-chr2 | PheGAA | 2728985 | 2729057 | + | cen2 |
| cry-chr2 | GluCTC | 2729514 | 2729585 | - | cen2 |
| cry-chr2 | LeuCAA | 2730082 | 2730186 | - | cen2 |
| cry-chr2 | LysCTT | 2730767 | 2730852 | + | cen2 |
| cry-chr2 | ArgACG | 2731068 | 2731140 | - | cen2 |
| cry-chr2 | ArgACG | 2733498 | 2733570 | - | cen2 |
| cry-chr2 | IleAAT | 2733945 | 2734018 | + | cen2 |
| cry-chr2 | AlaAGC | 2734350 | 2734423 | - | cen2 |
| cry-chr2 | ValAAC | 2734868 | 2734950 | + | cen2 |
| cry-chr2 | AspGTC | 2735291 | 2735361 | - | cen2 |
| cry-chr2 | AsnGTT | 2748139 | 2748212 | - | cen2 |
| cry-chr2 | MetCAT | 2748219 | 2748299 | - | cen2 |
| cry-chr2 | GluTTC | 2749357 | 2749428 | - | cen2 |
| cry-chr2 | AspGTC | 2757271 | 2757341 | + | cen2 |
| cry-chr2 | ValAAC | 2757616 | 2757698 | - | cen2 |
| cry-chr2 | AlaAGC | 2758143 | 2758216 | + | cen2 |
| cry-chr2 | IleAAT | 2758569 | 2758642 | - | cen2 |
| cry-chr2 | ArgACG | 2759016 | 2759088 | + | cen2 |
| cry-chr2 | LysCTT | 2759303 | 2759388 | - | cen2 |
| cry-chr2 | GluCTC | 2760005 | 2760076 | - | cen2 |
| cry-chr2 | GluCTC | 2771702 | 2771773 | + | cen2 |
| cry-chr2 | LysCTT | 2772390 | 2772475 | + | cen2 |
| cry-chr2 | ArgACG | 2772690 | 2772762 | - | cen2 |
| cry-chr2 | IleAAT | 2773136 | 2773209 | + | cen2 |
| cry-chr2 | AlaAGC | 2773562 | 2773635 | - | cen2 |
| cry-chr2 | ValAAC | 2774080 | 2774162 | + | cen2 |
| cry-chr2 | AspGTC | 2774437 | 2774507 | - | cen2 |
| cry-chr2 | GluTTC | 2782344 | 2782415 | + | cen2 |
| cry-chr2 | MetCAT | 2783473 | 2783553 | + | cen2 |
| cry-chr2 | AsnGTT | 2783560 | 2783633 | + | cen2 |
| cry-chr2 | ArgACG | 2788705 | 2788777 | - | cen2 |
| cry-chr2 | IleAAT | 2789151 | 2789224 | + | cen2 |
| cry-chr2 | AlaAGC | 2789557 | 2789630 | - | cen2 |
| cry-chr2 | ValAAC | 2790074 | 2790156 | + | cen2 |
| cry-chr2 | AspGTC | 2790431 | 2790501 | - | cen2 |
| cry-chr2 | AspGTC | 2796516 | 2796586 | + | cen2 |
| cry-chr2 | ValAAC | 2796869 | 2796951 | - | cen2 |
| cry-chr2 | AlaAGC | 2797336 | 2797409 | + | cen2 |
| cry-chr2 | IleAAT | 2797745 | 2797818 | - | cen2 |
| cry-chr2 | GluTTC | 2798548 | 2798619 | + | cen2 |
| cry-chr2 | GluTTC | 2801937 | 2802008 | - | cen2 |
| cry-chr2 | LeuAAG | 3030776 | 3030854 | - |  |
| cry-chr2 | MetCAT | 3146936 | 3147007 | + |  |
| cry-chr2 | SerTGA | 3212860 | 3212956 | + |  |
| cry-chr2 | MetCAT | 3212961 | 3213032 | + |  |
| cry-chr2 | LysTTT | 3250962 | 3251036 | - |  |
| cry-chr2 | HisGTG | 3374376 | 3374447 | + |  |
| cry-chr2 | AsnGTT | 3453220 | 3453293 | - |  |
| cry-chr2 | AspGTC | 3473162 | 3473232 | - |  |
| cry-chr2 | TrpCCA | 3487278 | 3487350 | + |  |
| cry-chr2 | GlnTTG | 3509685 | 3509756 | + |  |
| cry-chr2 | SerGCT | 3512052 | 3512146 | - |  |
| cry-chr2 | SerGCT | 3543510 | 3543604 | + |  |
| cry-chr2 | LysCTT | 3551555 | 3551637 | + |  |
| cry-chr2 | GlyGCC | 3603153 | 3603223 | - |  |
| cry-chr2 | ProAGG | 3610949 | 3611020 | + |  |
| cry-chr2 | GlyTCC | 3612247 | 3612317 | + |  |
| cry-chr2 | GlyTCC | 3616931 | 3617001 | + |  |
| cry-chr2 | ThrAGT | 3762741 | 3762812 | + |  |
| cry-chr2 | LeuCAA | 3771210 | 3771314 | + |  |
| cry-chr2 | ArgACG | 3776236 | 3776308 | + |  |
| cry-chr2 | HisGTG | 3819860 | 3819931 | - |  |
| cry-chr2 | GlyGCC | 3914569 | 3914639 | + |  |
| cry-chr2 | SerAGA | 3915190 | 3915271 | + |  |
| cry-chr2 | ValAAC | 3946555 | 3946638 | + |  |
| cry-chr3 | LeuAAG | 40194 | 40272 | + |  |
| cry-chr3 | LysCTT | 83679 | 83761 | - |  |
| cry-chr3 | AlaAGC | 103678 | 103751 | + |  |
| cry-chr3 | GlyGCC | 106935 | 107005 | - |  |
| cry-chr3 | ThrAGT | 186980 | 187051 | - |  |
| cry-chr3 | PheGAA | 242481 | 242553 | + |  |
| cry-chr3 | SerAGA | 389010 | 389091 | + |  |
| cry-chr3 | SerAGA | 413100 | 413181 | - |  |
| cry-chr3 | TyrGTA | 417806 | 417889 | - |  |
| cry-chr3 | TyrGTA | 418634 | 418717 | + |  |
| cry-chr3 | ArgTCT | 486107 | 486179 | + |  |
| cry-chr3 | ThrAGT | 601152 | 601223 | - |  |
| cry-chr3 | TyrGTA | 687190 | 687273 | - |  |
| cry-chr3 | GlyGCC | 697625 | 697695 | + |  |
| cry-chr3 | IleAAT | 700223 | 700296 | - |  |
| cry-chr3 | ProAGG | 700471 | 700542 | - |  |
| cry-chr3 | GlyGCC | 701100 | 701170 | - |  |
| cry-chr3 | ArgACG | 797292 | 797364 | - |  |
| cry-chr3 | AsnGTT | 818057 | 818130 | + |  |
| cry-chr3 | ThrAGT | 821031 | 821102 | - |  |
| cry-chr3 | AlaAGC | 900948 | 901021 | - |  |
| cry-chr3 | GluTTC | 913484 | 913555 | + | cen3 |
| cry-chr3 | LeuAAG | 914748 | 914826 | + | cen3 |
| cry-chr3 | AspGTC | 916129 | 916199 | + | cen3 |
| cry-chr3 | ValAAC | 916550 | 916632 | - | cen3 |
| cry-chr3 | AlaAGC | 917079 | 917152 | + | cen3 |
| cry-chr3 | IleAAT | 917485 | 917558 | - | cen3 |
| cry-chr3 | ArgACG | 917939 | 918011 | + | cen3 |
| cry-chr3 | ArgACG | 929697 | 929769 | + | cen3 |
| cry-chr3 | LysCTT | 929983 | 930068 | - | cen3 |
| cry-chr3 | LeuCAA | 930628 | 930732 | + | cen3 |
| cry-chr3 | GluTTC | 938349 | 938420 | + | cen3 |
| cry-chr3 | ThrAGT | 947532 | 947603 | + | cen3 |
| cry-chr3 | ThrAGT | 952280 | 952351 | - | cen3 |
| cry-chr3 | LeuAAG | 963327 | 963405 | + | cen3 |
| cry-chr3 | AspGTC | 964708 | 964778 | + | cen3 |
| cry-chr3 | ValAAC | 965159 | 965241 | - | cen3 |
| cry-chr3 | AlaAGC | 965690 | 965763 | + | cen3 |
| cry-chr3 | IleAAT | 966096 | 966169 | - | cen3 |
| cry-chr3 | ArgACG | 966550 | 966622 | + | cen3 |
| cry-chr3 | ArgACG | 968956 | 969028 | + | cen3 |
| cry-chr3 | LysCTT | 969244 | 969329 | - | cen3 |
| cry-chr3 | LeuCAA | 969889 | 969993 | + | cen3 |
| cry-chr3 | GluCTC | 970492 | 970563 | + | cen3 |
| cry-chr3 | PheGAA | 971020 | 971092 | - | cen3 |
| cry-chr3 | AlaAGC | 984839 | 984912 | + | cen3 |
| cry-chr3 | AspGTC | 986744 | 986814 | + | cen3 |
| cry-chr3 | ValAAC | 987140 | 987222 | - | cen3 |
| cry-chr3 | ValAAC | 1022461 | 1022543 | + |  |
| cry-chr3 | PheGAA | 1030689 | 1030761 | - |  |
| cry-chr3 | CysGCA | 1031226 | 1031297 | - |  |
| cry-chr3 | TrpCCA | 1174528 | 1174600 | + |  |
| cry-chr3 | ProAGG | 1210736 | 1210807 | - |  |
| cry-chr3 | ProAGG | 1264287 | 1264358 | + |  |
| cry-chr3 | LeuTAG | 1525949 | 1526027 | - |  |
| cry-chr3 | TrpCCA | 1548446 | 1548518 | + |  |
| cry-chr3 | AlaTGC | 1600309 | 1600380 | - |  |
| cry-chr3 | ArgCCG | 1654835 | 1654915 | - |  |
| cry-chr3 | LysCTT | 2030138 | 2030221 | + |  |
| cry-chr3 | ProAGG | 2059834 | 2059905 | + |  |
| cry-chr3 | ThrAGT | 2090541 | 2090612 | + |  |
| cry-chr3 | LysCTT | 2246722 | 2246804 | - |  |
| cry-chr3 | IleAAT | 2269370 | 2269443 | + |  |
| cry-chr3 | IleTAT | 2328647 | 2328745 | - |  |
| cry-chr3 | GluCTC | 2364126 | 2364197 | + |  |
| cry-chr3 | CysGCA | 2394500 | 2394571 | - |  |
| cry-chr3 | GlyTCC | 2546749 | 2546819 | - |  |
| cry-chr3 | ProAGG | 2547208 | 2547279 | - |  |
| cry-chr3 | GlyTCC | 2688401 | 2688471 | + |  |
| * Centromeric tDNAs are shaded purple |  |  |  |  |  |

**Supplementary Table 18:**
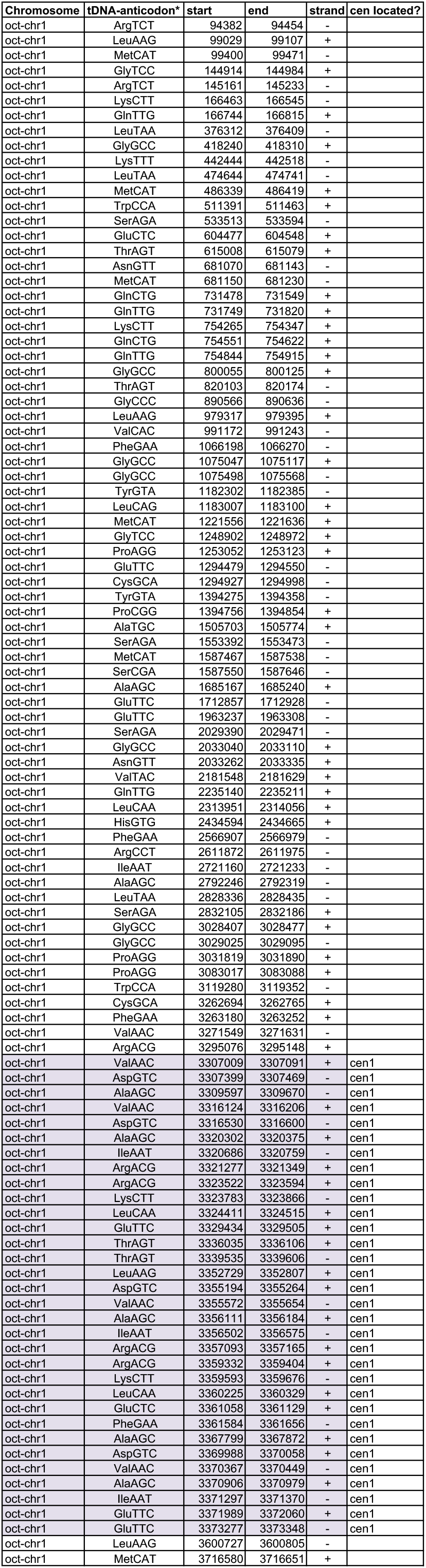
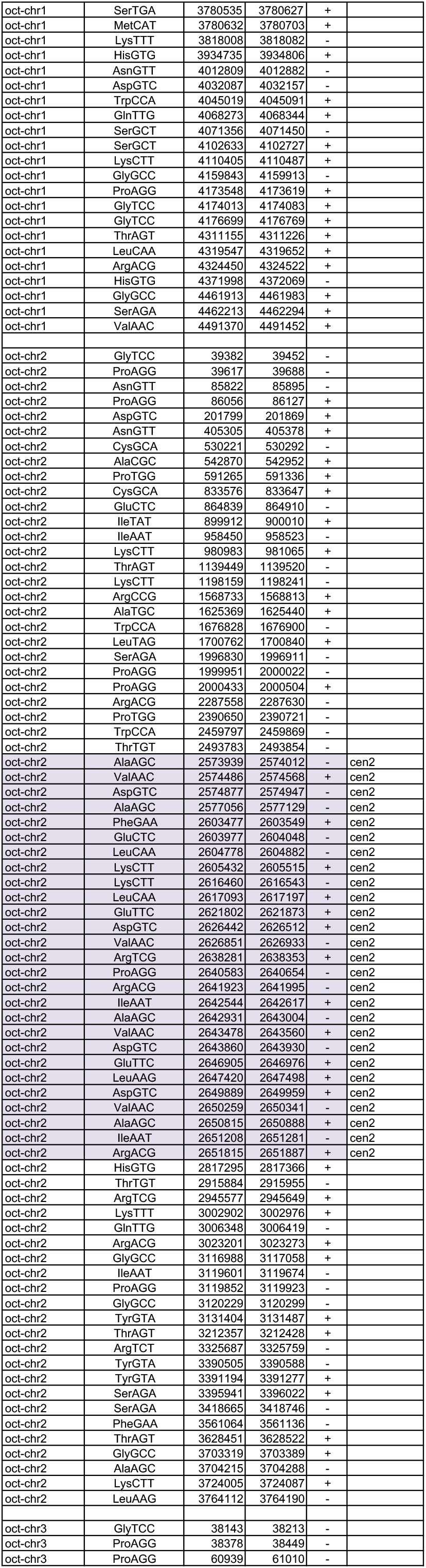
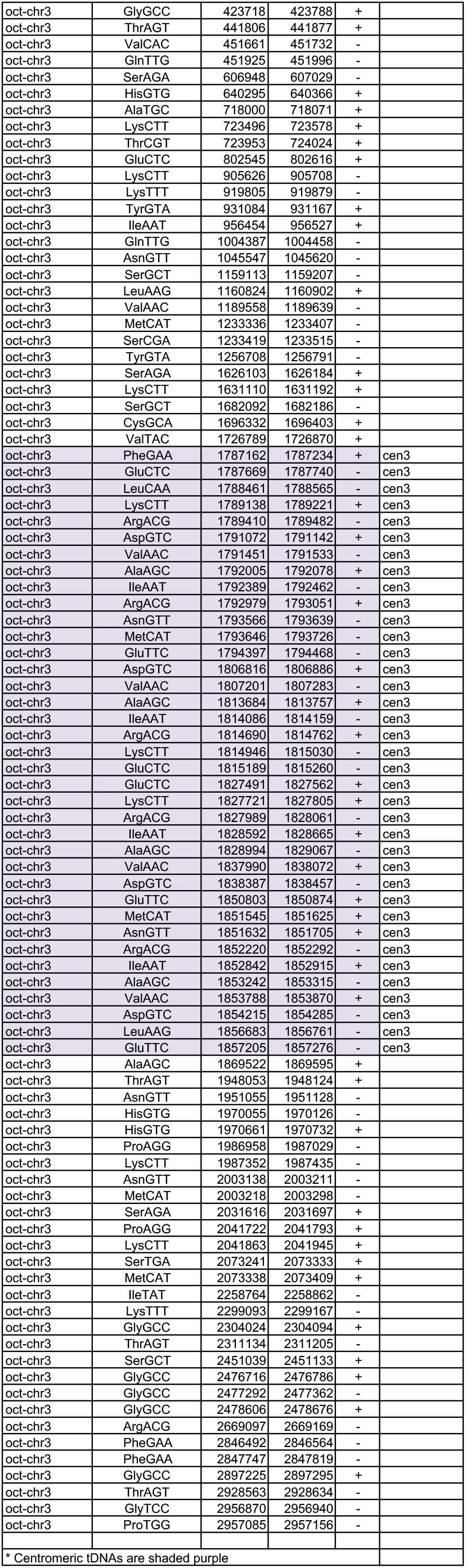
S. octosporus tDNA coordinates

**Supplementary Table 19:** Features associated with S. cryophilus and S. octosporus centromere DNA elements

| *S. cryophilus* | *S. octosporus* | Chromatin, features, comments |
| --- | --- | --- |
| cCNT-L | oCNT-L | CENP-A chromatin<br>Contain CNT-L and CNT-S elements<br>Contain CNT-L and CNT-S elements<br>Short central core, with long imrs<br>c-imr3 has LTRs which may act as boundaries |
| cCNT-S | oCNT-S |  |
| c-cnt1 | o-cnt2 |  |
| c-cnt2 | o-cnt3 |  |
| c-cnt3 | o-cnt1 |  |
| c-imr3 | o-imr1 |  |
| cFSAR-1 | oFSAR-1 | Heterochromatin. 5S-associated repeats |
| cFSAR-2 | oFSAR-2 |  |
| cFSAR-3 | oFSAR-3 |  |
| cTAR-11 | oTAR-11 | tDNA-associated repeats (TAR) - elements that are always associated with particular single tDNAs in both species and occur in equivalent positions. cTAR-14 and oTAR-14-ex contain retrotransposon remnants. |
| cTAR-12 | oTAR-12 |  |
| cTAR-13 | oTAR-13 |  |
| cTAR-14 | oTAR-14 |  |
| cHR-15 | oHR-15 | Heterochromatin. Not associated with tRNAs but occur in equivalent positions in the two species. |
| cHR-19 | oHR-19 |  |
| cTAR-4 | oTAR-4 | tDNA-associated repeats (TAR) elements that are always associated with multiple specific tDNAs. No/little CENP-A or heterochromatin (except TAR-7s which have heterochromatin on ~2 kb non-tDNA part of repeat). TARs may act as boundaries. |
| cTAR-5 | oTAR-5 |  |
| cTAR-6 | oTAR-6 |  |
| cTAR-7 | oTAR-7 |  |
| cTAR-8 | oTAR-8 |  |
| cTAR-9 | oTAR-9 |  |
| cTAR-10 | oTAR-10 |  |
| cTAR-11 | oTAR-11 |  |

**Supplementary Table 20:** List of Schiosaccharomyces strains used.

| Species | Strain ID | Genotype | Used for | Figure | Source Laboratory |
| --- | --- | --- | --- | --- | --- |
| *S. pombe* | A7408 | *h- cc2D6kb:cc1 ars1:nmt41-GFP-cnp1-NAT ade6-704-HYGMX6 his3-D1 leu1-32 ura4-DSE/D18 arg3?* | Centromere establishment | Fig 6 | Allshire (ref 15) |
| *S. pombe* | A7373 | *h- ade6-704-HYGMX6 his3-D1 leu1-32 ura4-DSE/D18? arg3? cc2D6kb:cc1* | Centromere establishment | not shown | Allshire (ref 15) |
| *S. pombe* | 6960 | *h- lys1+::cnp1-1 cnp1::ura4+ leu1-32 ura4-* | Complementation | Fig 5 | Takahashi (ref 30) |
| *S. pombe* | 1645 | *h+ ade6-210 arg3-D4 his3-D1 leu1-32 ura4-D18* | Localisation | Fig 5 | Allshire |
| *S. pombe* | 968 | *h<sup>90</sup>* | H3K9me2 ChIP-seq for mating-type region plot (h90) | Supp Fig S1 | Fantes / Sawin |
| *S. pombe* | 972 | *h-* | H3K9me2 and CENP-A ChIP-seq | Figs 2, Supp Fig S5 | Allshire |
| *S. pombe* | AMC501 | *h+ ade6-704 ura4-D18 leu1-32 rhn201 cdc6-L591G* | Minion nanopore sequencing, assembly of *S. pombe* genome | Supp Fig S5 | Nieduszynski / Murray |
| *S. pombe* | A6372 | *h- leu1 ura4 Δcen1::pADH1-loxP-KanR cd39 (tel1L neocentromere)* | CENP-A ChIP-seq for neocentromere regions | Fig 4 | Ishii / Takahashi (ref 29) |
| *S. pombe* | A6374 | *h- leu1 ura4 Δcen1::pADH1-loxP-KanR cd60 (tel1R neocentromere)* | CENP-A ChIP-seq for neocentromere regions | Fig 4 | Ishii / Takahashi (ref 29) |
| *S. octosporus* | A6969 | *h<sup>90</sup>* | PacBio sequencing, H3K9me2 and CENP-A ChIP-seq | Figs 1, 2, Supp Fig S2 | Rhind (ref 16) |
| *S. cryophilus* | A6972 | *h<sup>90</sup>* | PacBio sequencing, H3K9me2 and CENP-A ChIP-seq | Fig 1, 2, Supp Fig S3 | Rhind (ref 16) |
| *S. japonicus* | A1856 | *h<sup>90</sup>* | PacBio sequencing, H3K9me2 and CENP-A ChIP-seq | Supp Fig S5 | Rhind (ref 16) |

**Supplementary Table 21:** Summary of sequencing platforms used.

| Type of Sequencing | Species/ChIP | Sequencing Facility | Instrument / Chemistry /info etc |
| --- | --- | --- | --- |
| PacBio 1 | S. pombe<br>S. octosporus<br>S. cryophilus<br>S. japonicus | Biomedical Research Core Facilities,<br>University of Michigan | 2 SMRT cells each (no bluepippin)<br>PacBio RSII instrument, P4-XL |
| PacBio 2 with BluePippin | S. pombe<br>S. octosporus<br>S. cryophilus<br>S. japonicus | CSHL Cancer Center Next Generation<br>Genomics Shared Resource | no BluePippin: 1 SMRT cells each<br>With BluePippin technology:<br>8 SMRT cells each for Oct and Cry;<br>15 SMRT cells for Jap<br>PacBio RSII instrument with P4/C3 chemistry |
| Minion nanopore | S. pombe<br>AMC501 | Oxford | MinION nanopore sequencer |
| ChIP-seq | S. japonicus<br>CENP-A<br>H3K9me2 | BGI | Illumina GAII Single end |
| ChIP-seq | S. octosporus<br>CENP-A<br>H3K9me2 | Ark Genomics | HiSeq2000 Paired end |
| ChIP-seq | S. cryophilus<br>CENP-A<br>H3K9me2 | Ark Genomics | HiSeq2000 Paired end |
| ChIP-seq | S. pombe<br>CENP-A<br>H3K9me2 | Ark Genomics | HiSeq2000 Paired end |
| ChIP-seq | S. pombe<br>972h-H3K9me2 | Edinburgh Genomics | HiSeq2000 Paired end |
| ChIP-seq | S. pombe<br>h90<br>H3K9me2 | Allshire Lab | Miniseq Paired end |

**Supplementary Table 22:**
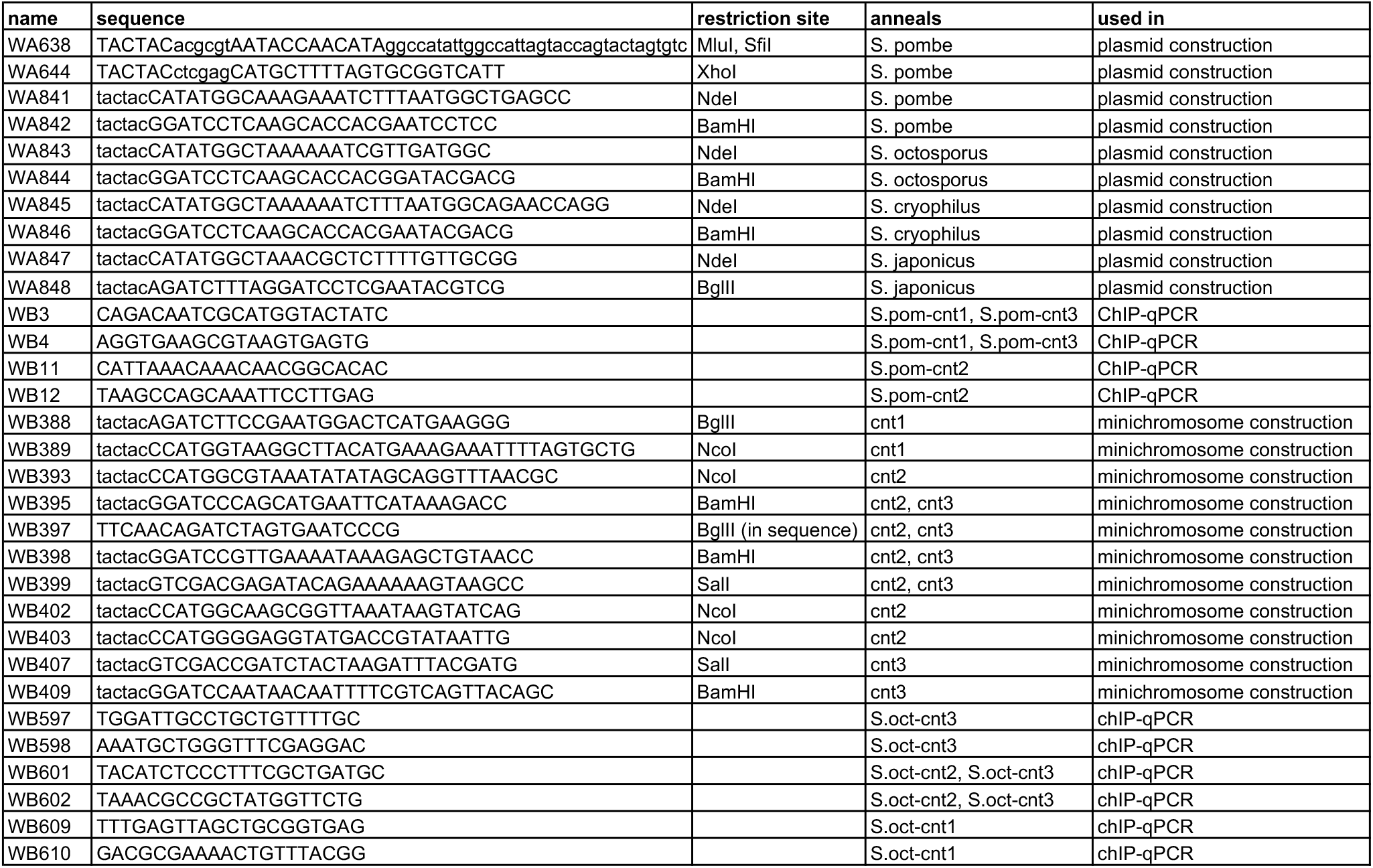
List of oligonucleotide primers used.

**Supplementary Table 23:** List of plasmids constructed and used.

| **plasmid / minichromosome** | **vector** | **source of insert/plasmid** | **primer F** | **primer R** |
| --- | --- | --- | --- | --- |
| pK(5.6kb)-MCS-ΔBam |  | S.pombe K repeat |  |  |
| pKp (pMC91) | pMC1 | S.pombe K repeat | WA638 | WA644 |
| pK-So-cnt1-3.2kb | pK(5.6kb)-MCS-DBam | S.oct-cen1 | WB388 | WB389 |
| pK-So-cnt2-6.5kb | pK(5.6kb)-MCS-DBam | S.oct-cen2 | WB402 | WB397 |
| pK-So-cnt2-4.7kb | pK(5.6kb)-MCS-DBam | S.oct-cen2 | WB393 | WB395 |
| pK-So-cnt2-10kb | pK-So-cnt2-6.5kb | S.oct-cen2 | WB398 | WB399 |
| pK-So-cnt3-6.5kb | pK(5.6kb)-MCS-DBam | S.oct-cen3 | WB407 | WB409 |
| pK-So-cnt3-3.6kb | pK(5.6kb)-MCS-DBam | S.oct-cen3 | WB397 | WB403 |
| pK-So-cnt3-2.6kb | pK(5.6kb)-MCS-DBam | S.oct-cen3 | WB395 | WB403 |
| pKp-So-cnt3-6.5kb | pKp | S.oct-cen3 | WB407 | WB409 |
| pKp-So-cnt3-3.6kb | pKp | S.oct-cen3 | WB397 | WB403 |
| pREP41-GFP-Sp-Cnp1 | pREP41-GFP (Craven et al) | S. pombe gDNA | WA841 | WA842 |
| pREP41-GFP-Sc-Cnp1 | pREP41-GFP | S. cryophilus genomic DNA | WA843 | WA844 |
| pREP41-GFP-So-Cnp1 | pREP41-GFP | S. octosporus genomic DNA | WA845 | WA846 |
| pREP41-GFP-Sj-Cnp1 | pREP41-GFP | S. japonicus genomic DNA | WA847 | WA848 |

